# A genome-wide epistatic network underlies the molecular architecture of continuous color variation of body extremities: a rabbit model

**DOI:** 10.1101/2021.10.07.463146

**Authors:** Julie Demars, Yann Labrune, Nathalie Iannuccelli, Alice Deshayes, Sophie Leroux, Hélène Gilbert, Patrick Aymard, Florence Benitez, Juliette Riquet

## Abstract

Deciphering the molecular architecture of coat coloration for a better understanding of the biological mechanisms underlying pigmentation still remains a challenge. We took advantage of a rabbit French experimental population in which both a pattern and a gradient of coloration from white to brown segregated within the himalayan phenotype. The whole experimental design was genotyped using the high density Affymetrix® AxiomOrcun™ SNP Array and phenotyped into 6 different groups ordered from the lighter to the darker. Genome-wide association analyses pinpointed an oligogenic determinism, under recessive and additive inheritance, involving genes already known in melanogenesis (*ASIP, KIT, MC1R, TYR*), and likely processed pseudogenes linked to ribosomal function, *RPS20* and *RPS14*. We also identified *(i)* gene-gene interactions through *ASIP:MC1R* affecting light cream/beige phenotypes while *KIT*:*RPS* responsible of dark chocolate/brown colors and *(ii)* a genome-wide epistatic network involving several others coloration genes such as *POT1* or *HPS5*. Finally, we determined the recessive inheritance of the English spotting phenotype likely involving a copy number variation affecting at least the end of the coding sequence of the *KIT* gene. Our analyses of coloration as a continuous trait allowed us to go beyond much of the established knowledge through the detection of additional genes and gene-gene interactions that may contribute to the molecular architecture of the coloration phenotype. Moreover, the characterization of a network including genes that contribute to melanogenesis and pigmentation, two processes affected in various human disorders, shows the potential interest of our rabbit model for transversal studies.

## Introduction

Understanding the molecular mechanism of coloration has been the goal of many genetic and evolutionary studies in a broad number of species [1–3]. More than a hundred of genes have been involved in coloration traits in model species such as drosophila or mice but also in wild species [4–7]. Specific color-producing cells contribute to animal coloration and patterns. The so-called “dermal chromotophore unit” [8] involves several types of chromatophores, including pterin and carotenoid-containing xantophors, iridophores with reflecting guanine platelets, and melanophores or melanocytes producing chemically melanin pigments. Phenotypic characteristics of animal coloration may be classified based on the patterning and/or type and amount of pigment produced through melanogenesis pathway [9]. The main genes that alter the development of melanocytes, corresponding to « spotting » phenotypes, are the Proto-Oncogene Receptor Tyrosine Kinase (*KIT*) and the Microphthalmia associated Transcription Factor (*MITF*) and the endothelin axis [10, 11]. Other genes such as Tyrosinase (*TYR*), Tyrosinase related Protein 1 (*TYRP1*), the Oculocutaneous albinism 2 (*OCA2*) and the Membrane-Associated Transporter Protein (*MATP*) affect melanin synthesis. Another group of genes related to pigment synthesis are those that control the switch between eumelanin and pheomelanin production; those with the strongest effect in this change are the Melanocortin 1 Receptor (*MC1R*) and the Agouti Signaling Protein (*ASIP*).

Most of our current knowledge is restricted to color traits exhibiting relatively simple variation and inheritance patterns. As example, mice carrying the viable or lethal yellow mutation (2 dominant mutations in the *Asip* gene) exhibit a phenotype that includes yellow fur called Agouti [12, 13]. Mutations in the *ASIP* gene have been highlighted in many other species displaying the Agouti coat color phenotype [14]. Similarly, red coat pigmentation in several mammals comes from mutations in the *MC1R* gene [14]. While many studies considered color traits as complex phenotypes analysing skin or hair colors as categories [15–17], most of recent analyses evaluated pigmentation as continuous variations [18–21]. Although GWAS have allowed for a greater understanding of the genetic component of many complex traits, the genetic effects highlighted are largely small and often focused on common SNP and additive genetic models. More and more studies explore alternative heritable components such as genetic interactions but there are still some challenges for identifying significant epistasis [22]. Skin, hair/coat pigmentation represent then pertinent phenotypes since the genetic determinism of coloration determined is likely polygenic involving a few genes with large effects [17, 23]. Typically, in the case of coat coloration, molecular interactions are known between MC1R and its antagonist ASIP peptide since gain-of-function *ASIP* mutations block MC1R signalling and lead to the production of red pheomelanin [24, 25]. Epistatic interaction of those two genes modulates wool color in an creole sheep breed [26]. Moreover, strong synergistic interactions have also been highlighted for other color traits such as skin/hair pigmentation in humans for which an interaction between *HERC2* and *MC1R* has been shown significantly associated [23].

In the European rabbit (*Oryctolagus cuniculus*), different coat colors have been selected through domestication and are nowadays fixed in specific breeds. Therefore, candidate gene approaches have allowed the identification of various mutations responsible of different phenotypes such as the dilution of the coat color [27] or the brown phenotype [28]. In rabbits, six loci (called A for Agouti, B for Brown, C for Color, D for Dilution, E for Extension and En for English Spotting) are involved in the coloration of the coat (Table 1). Notably, allelic heterogeneity with dominance/recessivity relations exists between different mutations within the single loci, resulting in distinct phenotypes. For instance, at the C locus (*TYR*), the *C*^*ch*^ mutation, responsible of the chinchilla phenotype, is dominant over the *C*^*h*^ mutation, itself associated with himalayan coat coloration which is dominant over the *c* mutation leading to albino phenotype [29] (Table 1). In addition, epistatic effects between both Extension and Agouti loci have been shown from a cross between a Champagne d’Argent buck and a Thuringian doe [30].

**Table 1:**
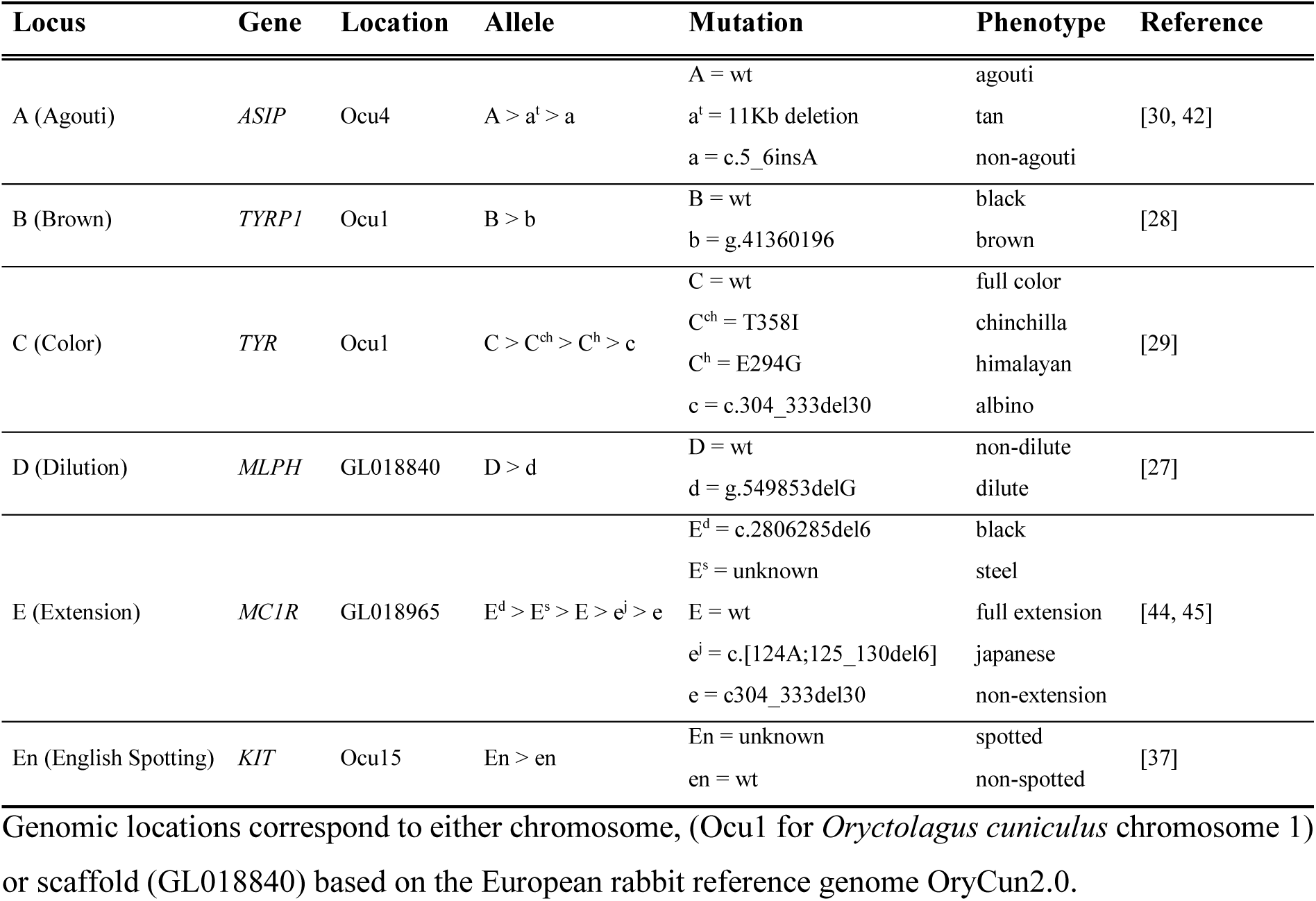
Genes and mutations responsible of coat coloration phenotypes in rabbits.

Although the mutation responsible of the himalayan phenotype has been identified, this trait has always been described in a simple way without considering both the gradient and the pattern of coloration occurring within the phenotype. A better insight of the coloration variability requires a fine characterisation of the phenotype to highlight dominance and/or recessive effects and epistatic interactions. Here, we propose a genome-wide investigation of coat color of body extremities using the high-density SNP rabbit beadchip (Affymetrix® AxiomOrcun™ SNP Array) in an experimental familial design. We *(i)* identified several significant loci including key genes involved in melanogenesis (*ASIP, MC1R, TYR* and *KIT*) but also atypical candidate genes which are processed pseudogenes linked to ribosomal proteins (*RPS20* and *RPS14*), *(ii)* highlighted how epistatic phenomena contribute to the genetic determinism of color variation of body extremities through *ASIP:MC1R* and *KIT:RPS* interactions regulating light and dark phenotypes, respectively and *(iii)* determined the recessive inheritance of the English spotting phenotype likely involving a copy number variation within the *KIT* gene. Altogether, our results bring new insights into the genetic determinism of the coat coloration variability emphasizing the key role played by interactions in the establishment of this complex trait.

## Results

### Several loci are significantly associated with coat color of body extremities

Coat color of body extremities was analysed as a quantitative trait with phenotypes numbered from 1 to 6 (called P1 to P6) and ordered from lighter to darker (Fig. 1a). A first exploration of the genetic determinism of coat color of body extremities was performerd using a simple a linear mixed model as outlined in the Methods section. A group of more than one hundred markers located on chromosome 1 (called Ocu1 for *Oryctolagus cuniculus* 1) showed significant associations, with the best signal for the SNP AX-146986391 (125,766,001 bp on Ocu1, p-value = 2.36*10^−56^) (Fig. 1b, and Additional file 1: Table S1). Additional significant and suggestive signals located on Ocu3 (AX-147059932, 131,847,470 bp, p-value = 9.70*10^−06^), Ocu4 (AX-147169681, 7,186,175 bp, p-value = 4.84*10^−06^), and Ocu15 (AX-146983797, 93,913,201 bp, p-value = 6.80*10^−11^) were obtained (Fig. 1b and Additional file 1: Table S1). In addition, groups of variants located on scaffolds GL018754 (AX-147179313, 18,452 bp, p-value = 2.83*10^−06^) and GL018965 (AX-147173908, 86,908 bp, p-value = 5.49*10^−05^), here regrouped for convenience in chromosome Unknown (an arbitrary chromosome that groups together all the scaffolds), also showed a suggestive association (Fig. 1b and Additional file 1: Table S1).

**Figure 1:**
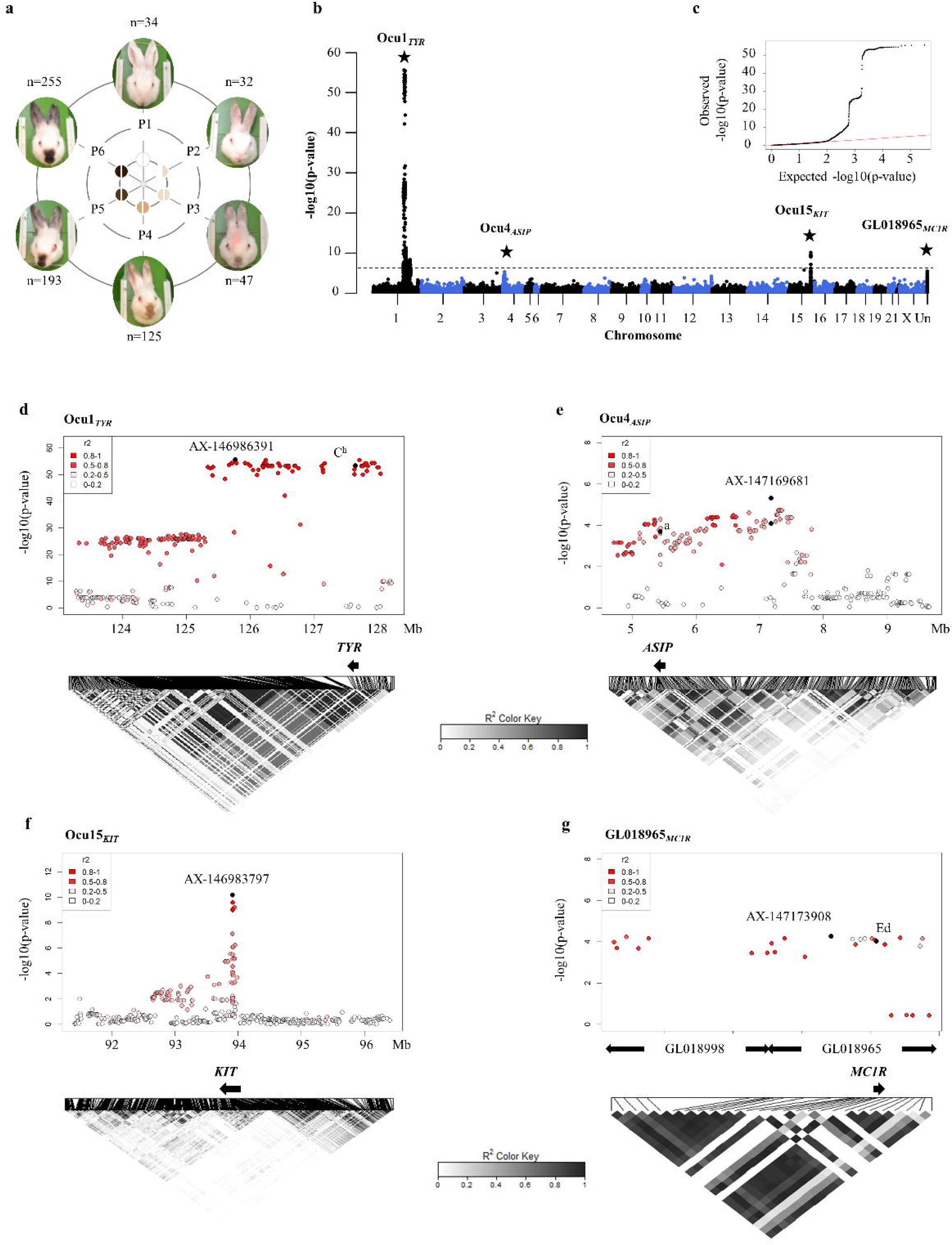
Coat coloration association results for the whole experimental design. **a**. Phenotypic classification. All rabbits were phenotyped in order to give them a phenotypic value. In total, 6 distinct coat pigmentations (P1 to P6) were considered and ordered in a gradient ranging from white to light and dark as quantitative phenotypes from 1 to 6. **b**. Manhattan plot. GWAS was performed using quantitative 1 to 6 phenotypes under a linear mixed model. Location of SNPs on the x-axis is based on the reference OryCun2.0 genome and all scaffolds were regrouped under an extra chromosome Unknown. The dashed line represents the 5% genome-wide threshold. **c**. Q-Q plot corresponding to the GWAS analysis. Highlights on a few regions of interest **d** for *TYR* locus on Ocu1 (Ocu1_*TYR*_), **e** for *ASIP* locus on Ocu4 (Ocu4_*ASIP*_), **f** for *KIT* locus on Ocu15 (Ocu15_*KIT*_) and **g** for *MC1R* locus on scaffold GL01865 (GL018965_*MC1R*_)) with regional Manhattan plots showing the best associated marker and already known mutations, and local linkage disequilibrium heatmap.

#### Two major genes with recessive effects are associated with white and spotting traits

We then focused on best associated markers to decipher how detected genomic regions contribute to the different coat color of body extremities. Analysis of the genotypic classes for variant Ocu1_*AX-146986391*_ among the different phenotypic groups showed that all individuals homozygous for the minor allele were P1 animals. Moreover, among the P1 class, only 5% of the individuals carry an allele 2, in a heterozygous manner suggesting that this locus, associated to white coat color, segregated with a recessive inheritance pattern (Additional file 2: Fig. S1a). To test this assumption, we performed association analyses comparing phenotype 1 (P1) versus the 5 remaining ones under different genetic models. A unique significant signal on Ocu1 with the best signal using the recessive model (AX-147087415, 127,829,702 bp, p-value = 1.30*10^−300^) was highlighted (Additional file 2: Fig. S1b). The *TYR* gene is 2 Mb downstream the best associated marker in a region showing high linkage disequilibrium (LD) (Fig. 1d). Already known mutations located in the *TYR* gene, *c* and *C*^*h*^, responsible of albino and himalayan phenotypes, respectively, did not show highest significant signals (Fig. 1d and Additional file 1: Table S2).

In a similar way, an excess of homozygote for the minor allele was observed for the best associated marker (Ocu15_*AX-146983797*_) located on Ocu15 (Additional file 2: Fig. S2a), suggesting that the light spotted color corresponding to phenotype P2 is mainly a Mendelian trait. We carried out a GWAS comparing phenotype 2 (P2) to combined light to dark brown extremities-colored rabbits (P3 to P6), excluding phenotype 1 (P1). The association signals for phenotype 2 was explained by variants of the chromosome 15 and scaffold GL018754 with the best p-values under a recessive model (Ocu15_*AX-146983797*_, 93,913,201 bp, p-value = 1.26*10^−144^ and GL018754_*AX-147115616*_, 74,205 bp, p-value = 1.72*10^−37^) (Additional file 2: Fig. S2b). The best associated marker on Ocu15 is located within the *KIT* gene (Fig. 1f). For the AX-147115616 SNP from scaffold GL018754, it is close to the *GSX2* and *PDGFRA* genes which are neighbours to the *KIT* gene in many species. This suggests that GL018754 is likely linked to Ocu15 as also suggested on the LD heatmap (Additional file 2: Fig. S3), and only one signal should be considered.

#### Five additional loci account for the remaining coat color of body extremities

To better decipher the molecular architecture of the other four phenotypic groups, we only considered those individuals (n=620) in further analyses. We used a Bayesian sparse linear mixed model, a more appropriate method for polygenic traits, allowing to estimate the number of quantitative trait loci (QTL) explaining the remaining coat color of body extremities. An estimation of 5 to 7 QTLs contributed to coloration phenotypes 3 to 6 (Additional file 2: Fig. S4a). Importantly, while 50% of the variance in phenotypes was explained by this model (Additional file 2: Fig. S4b), most of the genetic variance seemed due to QTLs (Additional file 2: Fig. S4c). We summed the sparse probabilities for the SNP inclusion on sliding windows containing 20 SNPs to amplify the identified signals from single variants (Additional file 2: Fig. S4d).

Two loci with large effects of approximately 0.6, located on Ocu1 and scaffold GL018965, showed high probabilities of being QTLs (70% and 78%, respectively). The *MC1R* gene belongs to the scaffold GL018965. Some known mutations within *MC1R* did not segregate in this population (japanese and extension alleles), but the best signal (AX-147194100) on GL018965 corresponded to the black dominant E^d^ mutation of the *MC1R* gene (Fig. 1g and Additional file 1: Table S2). Concerning the novel position on Ocu1 (AX-146995791), it matches to a gene-poor region on the OryCun2.0 genome assembly, and does not seem linked to any other region of interest. The closest genes are *RPS14* pseudogene (Ribosomal Protein S14, ENSOCUG00000026323), *RPS27* pseudogene (Ribosomal Protein S27, ENSOCUG00000024168), and *RORB* (RAR Related Orphan Receptor B). Two additional QTLs, located on Ocu1 and Ocu4, showed intermediate probabilities (36% and 55%, respectively), but also large effects of 0.9 and 0.4, respectively. The highlighted region on Ocu1 spanned the *TYR* locus and signal located on Ocu4 is approximately 1.5 Mb downstream the *ASIP* gene with a long structure of LD as previously shown for Ocu1 (Fig. 1e). Although several variants within *ASIP*, including the agouti *a* marker, were genotyped, they did not have the best p-values (Fig. 1e and Additional file 1: Table S2). Finally, 2 novel QTLs, located on Ocu13 and Ocu14, showed a trend of being QTLs with a probability above 15% for both and a large effect of 0.7 for the QTL on Ocu14. A few annotated genes (ENSOUG00000027919, ENSOUG00000000698, ENSOUG00000025838 and ENSOUG00000032896) including *RPS20* pseudogene (Ribosomal Protein S20, ENSOCUG00000025838) belong to the Ocu14 genomic region.

To fine-map intervals of interest, we used a Bayesian method considering the sum of the single-effects particularly well-suited to settings where variables are highly correlated and detectable effects are sparse. We validated 5 out of the 6 identified regions (exception of the Ocu13 locus) (Fig. 2a) and fine-mapped them in minimal Credible Set (CS) (Additional file 1: Table S3). One CS contained one SNP (AX-146995791) and the 4 others contained between 16 and 61 markers, with interval sizes ranging from 430 Kb to 2.2 Mb. The scaffold GL018965 was highlighted within a CS including also the scaffold GL018998 (Additional file 1: Table S3). While the region on GL018965 contained the Extension locus characterised by the *MC1R* gene, the interval on GL018998 pinpointed towards the *ANKRD11* gene. Homologous regions in human or mice highlighted the *MC1R* gene 600 Kb downstream the *ANKRD11* gene confirming that both scaffolds GL018998 and GL018965 might be linked, as confirmed by the LD heatmap (Fig. 1g). A unique CS of markers was identified on Ocu14 regrouping 2 groups of markers located more than 25 Mb away on the chromosome (Additional file 1: Table S3). Although these two groups of markers are located apart on the draft, linkage analysis using the familial meiosis of our pedigree indicates that these two groups are linked. A local genetic map could thus be established (Additional file 1: Table S4).

**Figure 2:**
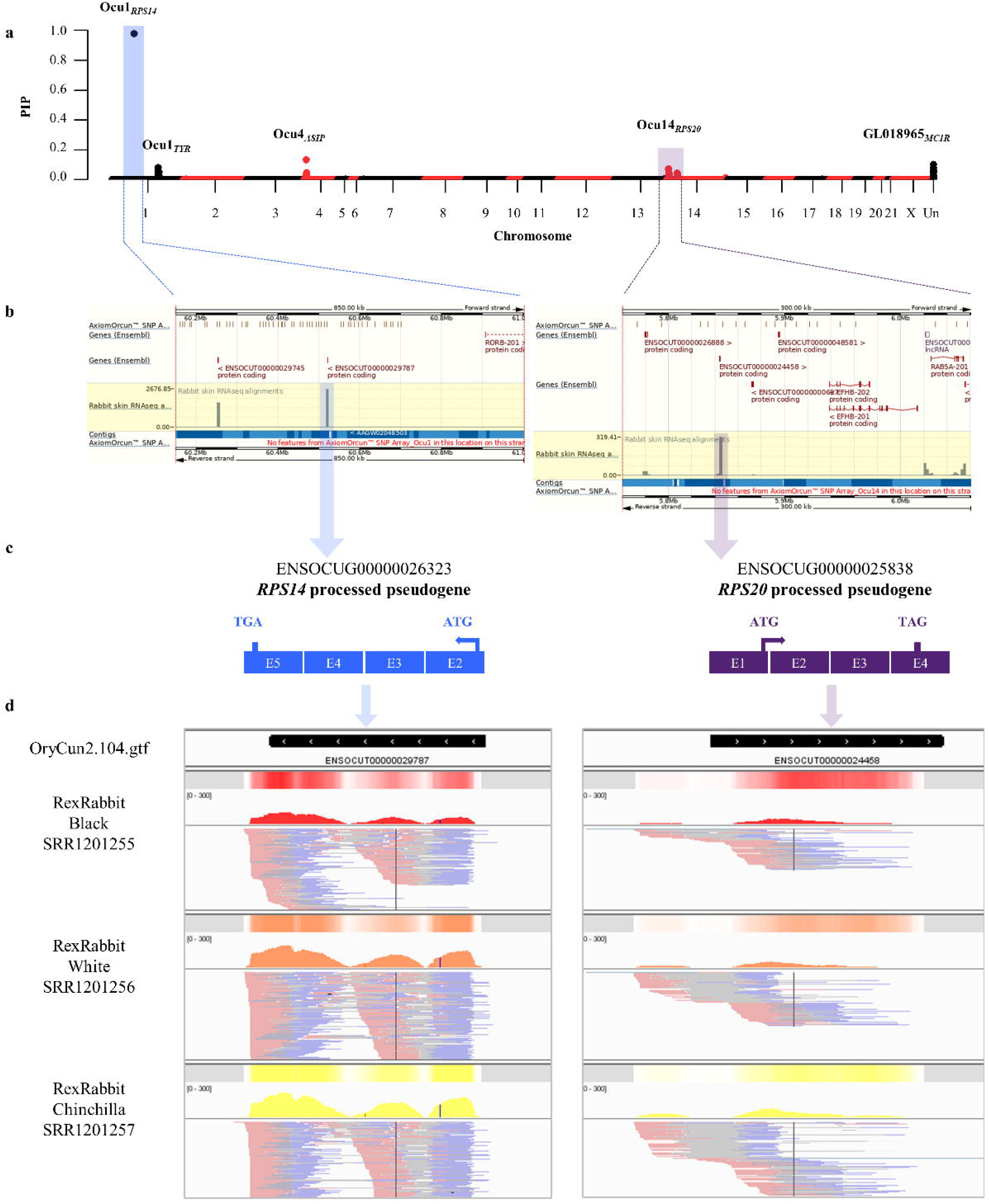
Highlights on processed pseudogenes of ribosomal proteins (*RPS14* and *RPS20*). **a**. Manhattan plot depicting posterior inclusion probabilities (PIP), which pinpoint fine mapping of genomic regions involved in cream to dark brown color variability including P3 to P6 colored rabbits. Location of SNPs on the x-axis is based on the reference OryCun2.0 genome and all scaffolds were regrouped under an extra chromosome Unknown. **b**. Screenshots of both Ocu1_*RPS14*_ and Ocu14_*RPS20*_ Ensembl regions. The first track represents the position of SNP on the Affymetrix® AxiomOrcun™ SNP Array, the second track represents the annotated genes and the third track highlighted in yellow shows the quantification of transcripts in the generic rabbit sample (accession number SAMN00013655). **c**. Representation of both *RPS14* and *RPS20* processed pseudogenes in the OryCun2.0 reference genome. **d**. Screenview of Intregrative Genome Viewer showing alignments of reads obtained after RNA-seq experiments of skin samples from 3 Rex rabbits carrying black coat (accession number SRR1201255), white coat (accession number SRR1201256) or chinchilla coat (accession number SRR1201255).

To likely identify candidate genes belonging to novel highlighted Ocu1 and Ocu14 intervals, we analysed publicly available RNA-seq data extracted from skin of rabbits including a generic sample (accession number SAMN00013655), Rex black rabbit (accession number SAMN02693835), Rex white rabbit (accession number SAMN02693836) and Rex chinchilla rabbit (accession number SAMN02693834). Only 5 annotated genes (2 and 3 in Ocu1 and Ocu14 genomic regions, respectively) were quantified in skin (Fig. 2b). It occurred that the three *RPS* pseudogenes (*RPS14* - ENSOCUG00000026323, *RPS27* - ENSOCUG00000024168 and *RPS20* - ENSOCUG00000025838) looked like processed pseudogenes since they all carried both START and STOP codons in the OryCun2.0 genomic reference sequence. While 8 and 10 pseudogenes of *RPS14* and *RPS20*, respectively are sparse in the rabbit genome, only the two copies located on Ocu1 and Ocu14, respectively are processed pseudogenes carrying the transcription initiation and ending codons (Fig. 2c). All of them were expressed in the skin tissue of the generic rabbit sample (Fig. 2b). Despite the lack of statistics between the three samples of skin of Rex rabbits (black *vs*. white *vs*. chinchilla), less reads mapped to both *RPS14* and *RPS20* processed pseudogenes in the black Rex rabbit (Fig. 2d).

### Gene-gene interactions contribute to the determinism of coat color of body extremities

To assess whether specific interactions accounted for the variability of coat color of body extremities, we first evaluated pairwise genotypic distribution across the 6 phenotypic groups (P1 to P6) between the 7 selected markers from previous analyses (AX-146995791 (Ocu1_*RPS14*_), AX-147087415 (Ocu1_*TYR*_ [P1]), AX-147073566 (Ocu1_*TYR*_ [P3 to P6]), AX-147097074 (Ocu4_*ASIP*_), AX-147006836 (Ocu14_*RPS20*_), AX-146983797 (Ocu15_*KIT*_), and AX-147194100 (GL018965_*MC1R*_). Significant interactions between variants located on Ocu1 likely reflecting linkage were identified (Fig. 3a and Additional file 1: Table S5). Significant epistasis were also highlighted between best markers of Ocu15_*KIT*_ and Ocu14_*RPS20*_ (p-value = 0.0156) and a trend was observed for Ocu4_*ASIP*_:Ocu14_*RPS20*_ (p-value = 0.08127) and Ocu4_*ASIP*_:GL018965_*MC1R*_ (p-value = 0.1056) (Fig. 3a and Additional file 1: Table S5).

**Figure 3:**
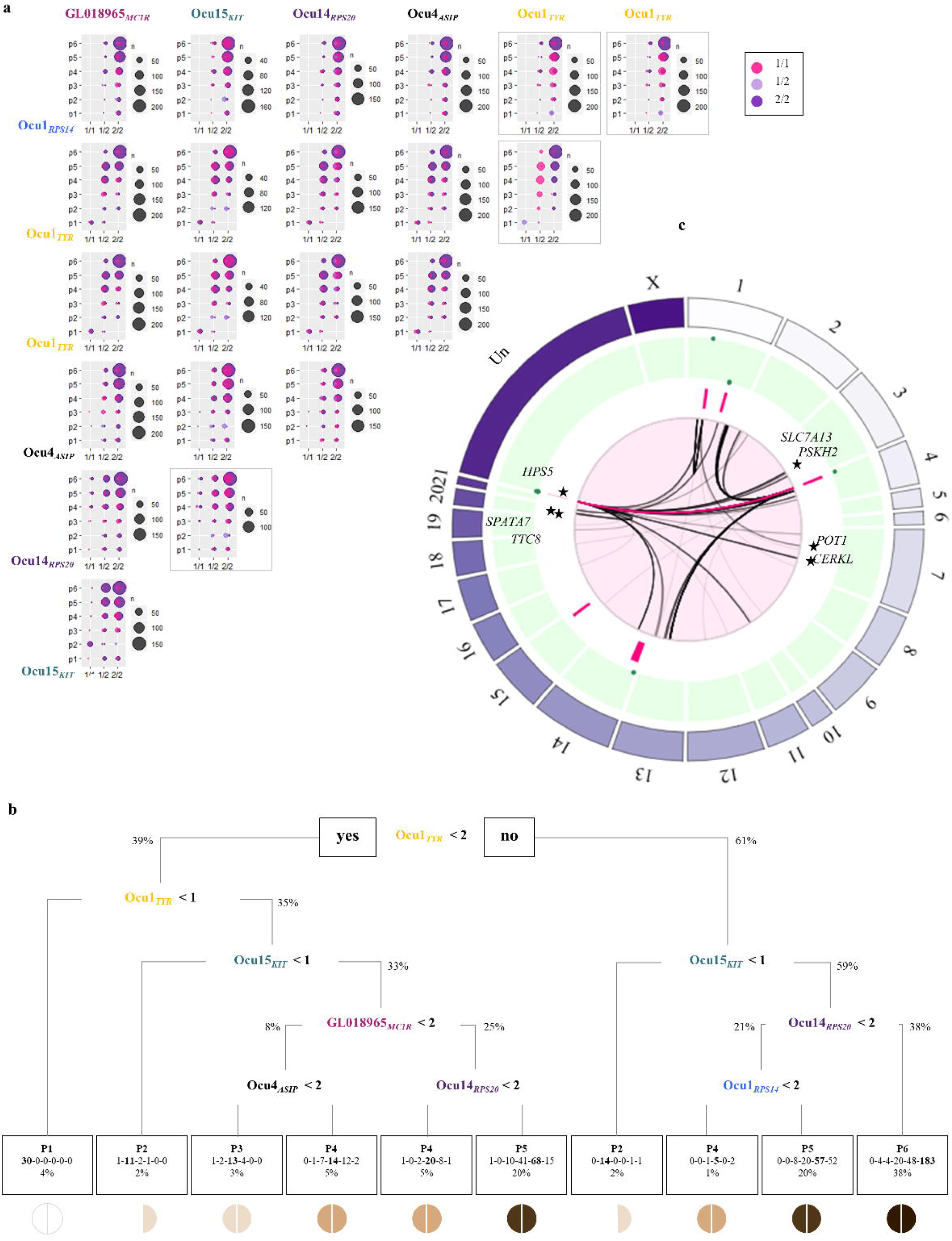
Epistatic interactions underlying the molecular architecture of coat coloration. **a**. Pairwise epistatic interaction. Each count plot represents the genotypic distribution between the most significant markers of two out of the 7 associated loci previously identified. The x-axis represents genotypes at the locus mentioned on the line while the colored circles (pink, light purple and dark purple) represent genotypes at the locus mentioned on the column. The y-axis represents the 6 coat coloration phenotypic groups. The framed boxes correspond to significant interaction using the linear regression model. **b**. Decision tree. All rabbits were considered for all highlighted loci. The nodes discriminate genotypes at each involved marker with 0, 1 and 2 genotypes corresponding to the presence of 0, 1 or 2 minor alleles, respectively. The final leaves are the distinct colored phenotypes with the repartition of rabbits within each group from 1 to 6. **c**. Epistatic network. Only rabbits with colored phenotypes P3 to P6 were considered. Circular plot shows from external to internal tracks: chromosomes track (purple track), posterior inclusion probability (PIP) track ranging from 0 to 1 (green track), 6 identified regions track (Ocu1_*RPS14*_, Ocu1_*TYR*_, Ocu4_*ASIP*_, Ocu14_*RPS20*_, Ocu15_*KIT*_ and GL018965_*MC1R*_) (pink track) and interactions track (central circle). Stars represent significant interactions with genes already known to contribute to coloration process.

We then built a classification tree based on the genotypes at each marker of this set of 7 markers, to apprehend epistasis between the different loci and genotypes-phenotypes relationships (Fig. 3b). As expected, Ocu1_*TYR*_ and Ocu15_*KIT*_, were found as major genes responsible of white (P1) and spotted (P2) phenotypes, respectively. For the remaining phenotypic groups, approximately 28% (13/47), 31% (39/125), 65% (125/193) and 72% (183/255) of individuals seemed correctly classified for phenotypes P3, P4, P5 and P6, respectively (Fig. 3b). While P3 and P4 phenotypes seem mostly explained by interaction between Ocu4_*ASIP*_ and GL018965_*MC1R*_, interactions between both loci involving *RPS* processed pseudogenes, Ocu1_*RPS14*_ and Ocu14_*RPS20*_, seemed involved in darker P5 and P6 phenotypes (Fig. 3b).

To disentangle the most significant genetic components including gene-gene interactions that contribute to the determinism of coat color of body extremities, we used the Bayesian selection criterion BIC to select significant interactions in a stepwise procedure applied to linear regression models. Since phenotypes P1 and P2 seemed exclusively explained by Ocu1_*TYR*_ and Ocu15_*KIT*_ major genes, we considered only phenotypes P3 to P6 but incorporated in our genetic model all possible combinations between the set of 7 selected variants. The best returned model included the different markers as main effect with the most significative positive effects for Ocu4_*ASIP*_, Ocu14_*RPS20*_ and GL018965_*MC1R*_. Significant epistasis were highlighted with the most significant effect for the Ocu4_*ASIP*_:GL018965_*MC1R*_ interaction with a negative effect on phenotypes (Additional file 2: Fig. S5). In addition, Ocu15_*KIT*_ showed significant epistatic effects with both Ocu1_*TYR*_ and ribosomal genes *RPS* processed pseudogenes (Additional file 2: Fig. S5).

Finally, we considered epistasis between 6 of the 7 markers and the rest of the genome to analyse with a wide angle the coloration genes network. Only one variant, AX-147073566, was considered for Ocu1_*TYR*_ since both selected markers (AX-147073566 and AX-147087415) are very close. We performed pairwise epistasis tests using an adaptive shrinkage method estimating both local false sign rates (lfsr) and effect sizes, adapted to limited sample size for increasing statistical power [31]. The best interactions with a lfsr < 10^−03^ are shown on the Fig. 3c. Approximately 570 significant interactions were obtained with more than 85% of those involving Ocu4_*ASIP*_ and GL018965_*MC1R*_ markers with clusters of variants between each other (Additional file 1: Tables S6-S9). The best significant effect was observed for Ocu4_*ASIP*_:GL018965_*MC1R*_ interaction (5,857,504 bp on Ocu4 and 400 Kb away to the *ASIP* gene, lfsr = 1.33*10^−10^) with a similar effect to the previously one observed (Additional file 1: Table S8 and Additional file 2: Fig. S5). Interestingly, several novel highlighted epistatic interactions pinpointed to genomic regions spanning genes involved in coloration pathways or pigmentation linked disorders, such as *HPS5, POT1, TTC8, SPATA7* or likely *SLC24A4, CERKL, PSKH2* and *SLC7A13* that are less than 500 Kb away (Fig. 3c and Additional file 1: Tables S6-S9).

### A copy number variation likely overlaps the Ocu15_*KIT*_ locus

Since several known mutations, located within genomic regions of interest, did not show best significant signals, we searched for CNV. We focused our research of CNV to regions spanning previously intervals identified as associated to phenotypes. Instead of the characterization of individual structural variants, we analysed the mean value of LRR (Log R Ratio) and BAF (B Allele Frequency) at each SNP of the region in the different phenotypic groups. For Ocu15_*KIT*_, aberrant values of means of LRR, ranging below −0.97 and above 0.57, were observed for spotted rabbits (phenotype P2) in an interval containing the *KIT* gene, which referred to several markers that can then be considered as involved in a CNV (Fig. 4a). In addition, the distribution of BAF values for all individuals of the experimental design, not only the spotted colored rabbits, showed a typical profile of a CNV since we detected many values outside the expected 0, 0.5 and 1 categories (Fig. 4a). This suggests that the CNV affecting the *KIT* gene segregated within the whole protocol. The CNV seemed to affect the *KIT* gene as shown for 1 marker located within the gene for which the signal intensity seemed different between both alleles and additional groups of genotypes might be deduced (Fig. 4b). A similar pattern was observed for 5 markers located within the *KIT* gene suggesting a CNV of at least 5 Kb (93,911,613 bp − 93,916,801 bp) affecting the 5 last exons of the gene including the STOP codon. A GWAS using individual LRR values as markers and comparing spotted colored animals (P2) to combined all others extremities-colored rabbits except P1 showed a significant signal on Ocu15 in the interval containing the *KIT* gene (Fig. 4c).

**Figure 4:**
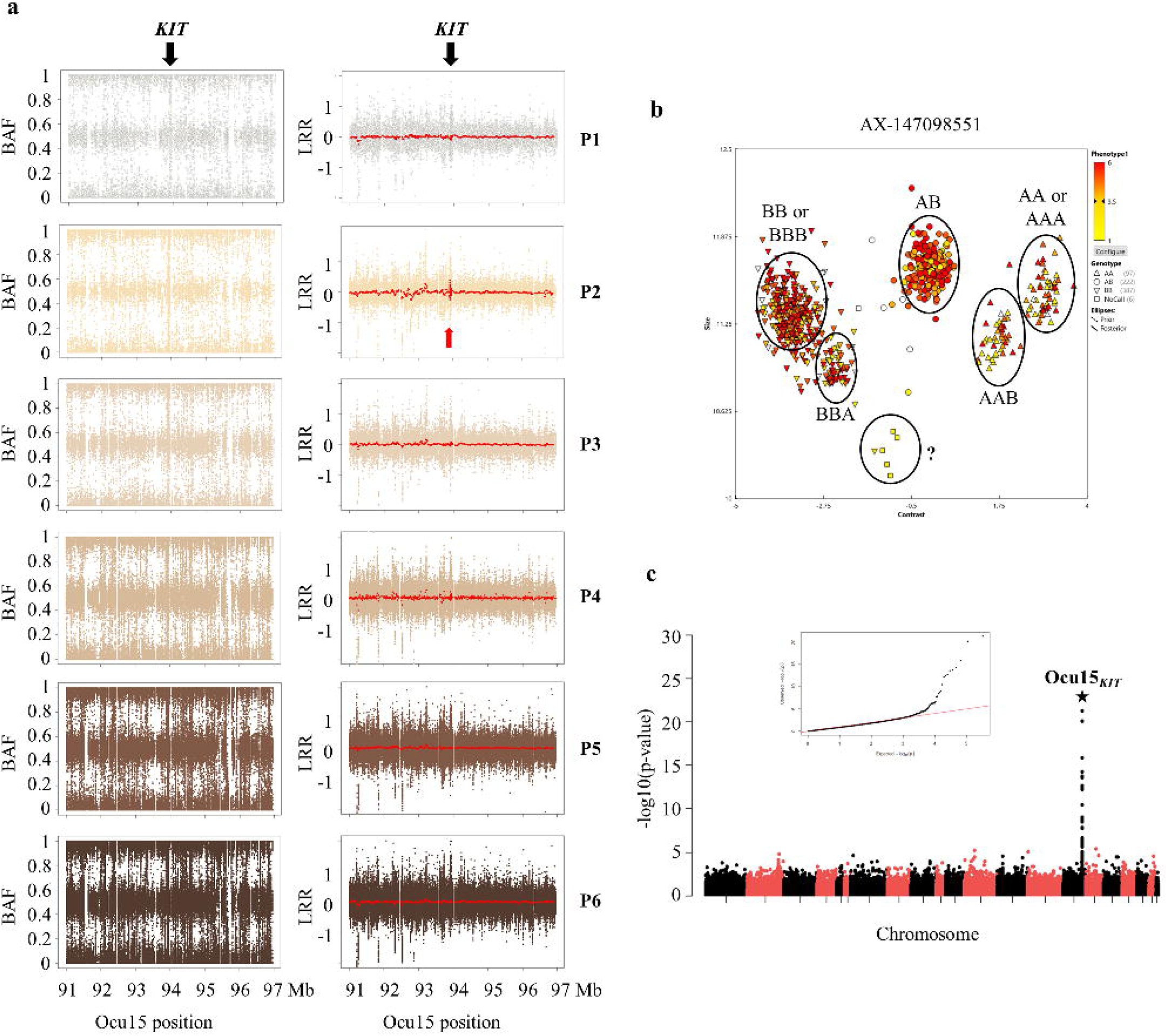
Copy number variation within *KIT* associated with white spotting phenotype. **a**. B Allele Frequency (BAF) and Log R Ratio (LRR) plots. BAF and LRR values were plotted per phenotype (P1 to P6) for all individuals for a region of 6 Mb centered around the *KIT* gene. For each marker, the average LRR per phenotype was evaluated and a sliding average using a window of 10 markers was shown in red on LRR plots. **b**. Genotyping data extracted from the Axiom™ Analysis Suite Software 4.0.3.3 for the SNP AX-147098951 (93,916,801 bp) located within the *KIT* gene. Additional groups of genotypes likely reflecting a copy number variation spanning this marker. **c**. Manhattan plot. GWAS was performed using LRR values for each SNP and by a comparison of rabbits with phenotype P2 *vs*. all others groups of colored animals (P3 to P6).

## Discussion

### Effect of known genes and/or mutations on the coat color of body extremities

The interval located on Ocu1 contains the *TYR* gene, which is an essential enzyme of the melanin biosynthesis from the tyrosine within melanosomes [9]. Regarding the Himalayan C^*h*^ allele, it perfectly discriminated white rabbits from all other individuals with coat color at their body extremities. Surprisingly, for the white phenotype (called P1), the best association signal under the recessive model was obtained for AX-147087415 instead of the Albino c allele (AX-146982536). Indeed, 4 individuals classified in phenotype P1 are heterozygous for this known allele. Additional manual genotyping for this variant showed genotyping errors from the SNP array with 2 out of 4 animals homozygous for the Albino c allele. The remaining 2 incoherent rabbits are likely phenotypic errors. This may confirm the causal effect of this mutation on the genetic determinism of the white coat coloration [29].

Interestingly, markers located within the genomic region spanning the *TYR* locus also showed association signals when only phenotypes P3 to P6 were considered. The LD structure measured in this region suggested the segregation of two distinct haplotypes with significant p-values. The haplotype carrying the region upstream the *TYR* gene may likely been involved in the variability of coat color of body extremities under an additive determinism. As observed on the classification tree, heterozygosity at the Ocu1_*TYR*_ seemed more correlated with lighter P3 and P4 phenotypes while homozygosity for the major allele seemed more represented within darker P5 and P6 groups of rabbits. In a conditional and reversible gene expression knockdown mouse model, the authors showed that TYR was necessary not only for the synthesis of melanin, but also for the complete maturation of the stage IV melanosome [32]. This system where the TYR protein was depleted at a level that was sufficient to alter coat color but not sufficient to significantly alter melanin accumulation, likely suggested the potential effect of an additional genetic variation at the *TYR* locus [32]. In accordance with these observations, our results complicate a little more the molecular basis and allelic series imbalance of the coloration C locus.

The KIT receptor is also a key regulator activating the synthesis of eumelanin through the MAPK signalling pathway [9]. We figured out a structural variant spanning the *KIT* gene, a CNV of at least 5 Kb affecting the 5 last exons of the gene including the STOP codon, might contribute to the P2 phenotype (white rabbits with coloration at their body extremities except their noses, also considered as spotted phenotype). The *KIT* gene has been described in several species associated with coloration traits, especially with white spotting phenotype in cats [33], donkeys [34], camels [35], horses [36] and English spotting phenotype in rabbits [37]. In addition, structural variants involving *KIT* have been identified and associated with white spotting phenotypes such as in horses in which a heterozygous 1.9 Kb deletion affecting exons 10-13 of the *KIT* gene represented a true null allele responsible of the depigmentation phenotype [38]. Although an accurate characterization of the structural variant affecting the *KIT* gene is needed, our results strongly suggest that a CNV within the *KIT* gene is the causal mutation of the English spotting phenotype in rabbits [37].

The region on Ocu4 spans the *ASIP* gene, a signalling ligand initiating the synthesis of pheomelanin pigment through its binding to the MC1R receptor [9]. In many species, the *ASIP* gene is involved in coloration traits with the most known is the agouti phenotype [39] In rabbits, the causal mutation disrupting the protein discriminates between full and a dual coloration due to the expression of pheomelanin [30]. Here, results focusing on the *ASIP* locus are less clear since the best association signal was located more than 1 Mb downstream the agouti *a* marker with an intermediate level of LD suggesting another variants involved in the light coat color of body extremities (phenotypes P3 and P4). However, aberrant BAF values focused on the *ASIP* gene might suggest a CNV spanning the *ASIP* gene but LRR values did not seem confirming it (data not shown). More and more studies have highlighted structural variants encompassing or close to the *ASIP* gene and associated with coat colored phenotypes in different domestic species [40, 41]. Indeed, a 11 Kb deletion affecting the *ASIP* gene was the most likely variant for the black and tan phenotype in rabbits [42]. In addition, populations analyses performed in livestock have shown that copy number variants underlying breed-defining coat color phenotypes revealed selection signatures [43]. Although complementary experimentations are needed to deeper characterize the mutation at the *ASIP* locus, our results suggested additional allelic heterogeneity at the coloration A locus in rabbits.

The genomic region carrying the *MC1R* gene is also associated with coat color of body extremities in our experimental design. While the binding of the α-MSH peptide on MC1R allows the synthesis of eumelanin *via* the cAMP signaling pathway, the ASIP peptide has an antagonist effect blocking the biosynthesis of eulemanin in favour of pheomelanin production [24, 25]. Although black and japanese alleles were genotyped [44, 45], only the black dominant E^d^ allele segregated within our experimental design and was the best associated marker with coat color traits. Only 2 haplotypes spanning the whole GL018965 scaffold have been identified throughout the experimental design (data not shown) clearly suggesting either the causal role of the E^d^ allele or an additional mutation within the same haplotype. In contrary to both *ASIP* and *KIT* loci, very few examples of CNV have been identified involving the *MC1R* gene [46]. Here, searching for structural variants was not appropriate given that *MC1R* is located within a scaffold containing a low number of variants. However, given the LD structure within the region, identifying and discriminating between several variants may remain challenging for the *MC1R* locus.

### Processed pseudogenes of ribosomal proteins are likely involved into the coat color of body extremities

More importantly, two regions, Ocu1_*RPS14*_ and Ocu14_*RPS20*_, also account for the molecular architecture of the coat color of body extremities. Both regions contain few annotated genes on the OryCun2.0 genome and only some of them are expressed in the skin. Three of them belong to the 40S ribosomal proteins which are likely the *RPS14, RPS27* and *RPS20* genes, respectively, by sequence homology. The three ribosomal genes located within intervals of interest looked like pseudogenes which is very common for ribosomal genes in several species [47, 48]. Approximately 30 *RPS* genes and 100 *RPS* pseudogenes are dispersed in mammalian genomes [49, 50]. In addition, analyses performed from publicly available RNA-seq data obtained from skin including a generic sample, a Rex black, a Rex white and a Rex chinchilla rabbit highlighted quantification of messengers from both *RPS14* and *RPS20* processed pseudogenes. The study of Tonner et al. detected transcription of ribosomal protein pseudogenes in diverse human tissues from RNA-seq data [51]. Unlike *RPS* genes that are constitutively expressed in almost all tissues, *RPS* pseudogenes are differentially expressed, suggesting that they may contribute to tissue-specific biological processes [50]. Two studies carried in mice [31] and zebrafish [53] showed coloration defects when mutations in *Rps20* and *Rps14*, respectively, were induced. Indeed, a study in mice reported 2 mouse dark skin (*Dsk*) loci caused by mutations in *Rps19* and *Rps20* with a common signalling pathway through the stimulation of Kit ligand (kitl) expression by p53 [54]. Hence, a ribosome defect in keratinocytes may mimic ultraviolet response to keratinocytes resulting in a p53 induction in these cells that may drive melanocytes proliferation/migration *via* kitl signalling; this may lead to an hyperpigmentation tanning response [55, 56]. In addition, deficiency in rps14 in zebrafish led to a delayed pigmentation through an increase of p53 activity [53]. Moreover, the comparison of the transcriptional profiles of human cell lines of dark and light melanocytes under basal conditions and following ultraviolet-B irradiation showed an interaction between ribosomal proteins and the p53 signalling pathway [52]. Although deeper sequencing analyses of both DNA and RNA are needed to consolidate our assumption, our results in the light of the literature suggested ribosomal genes especially *RPS20* and *RPS14* processed pseudogenes, as pertinent and novel candidate genes likely involved in the genomic basis of coat coloration.

### Epistatic network contributes to the genetic determinism of the coat color of body extremities

Here, we analysed the color variation of body extremities as a continuous trait allowing a better understanding of gene-gene interactions involved in the molecular architecture of the trait. The global overview of the coat color of body extremities determinism in our rabbit model showed the existence of an epistatic interaction network involving core genes but also likely additional genes with small effects (Fig. 5). Our results suggested that *(i)* Ocu1_*TYR*_ seemed to dictate whether pigmentation is produced or not and when the coloration occurred, it is restricted to body extremities, *(ii)* interactions among the others genes (Ocu1_*RPS14*_, Ocu4_*ASIP*_, Ocu14_*RPS20*_ and GL018965_*MC1R*_) seemed to dictate the amount of pigment produced and *(iii)* Ocu15_*KIT*_ seemed to control where pigment is deposited, all body extremities or restricted to some extremities (Fig. 5).

**Figure 5:**
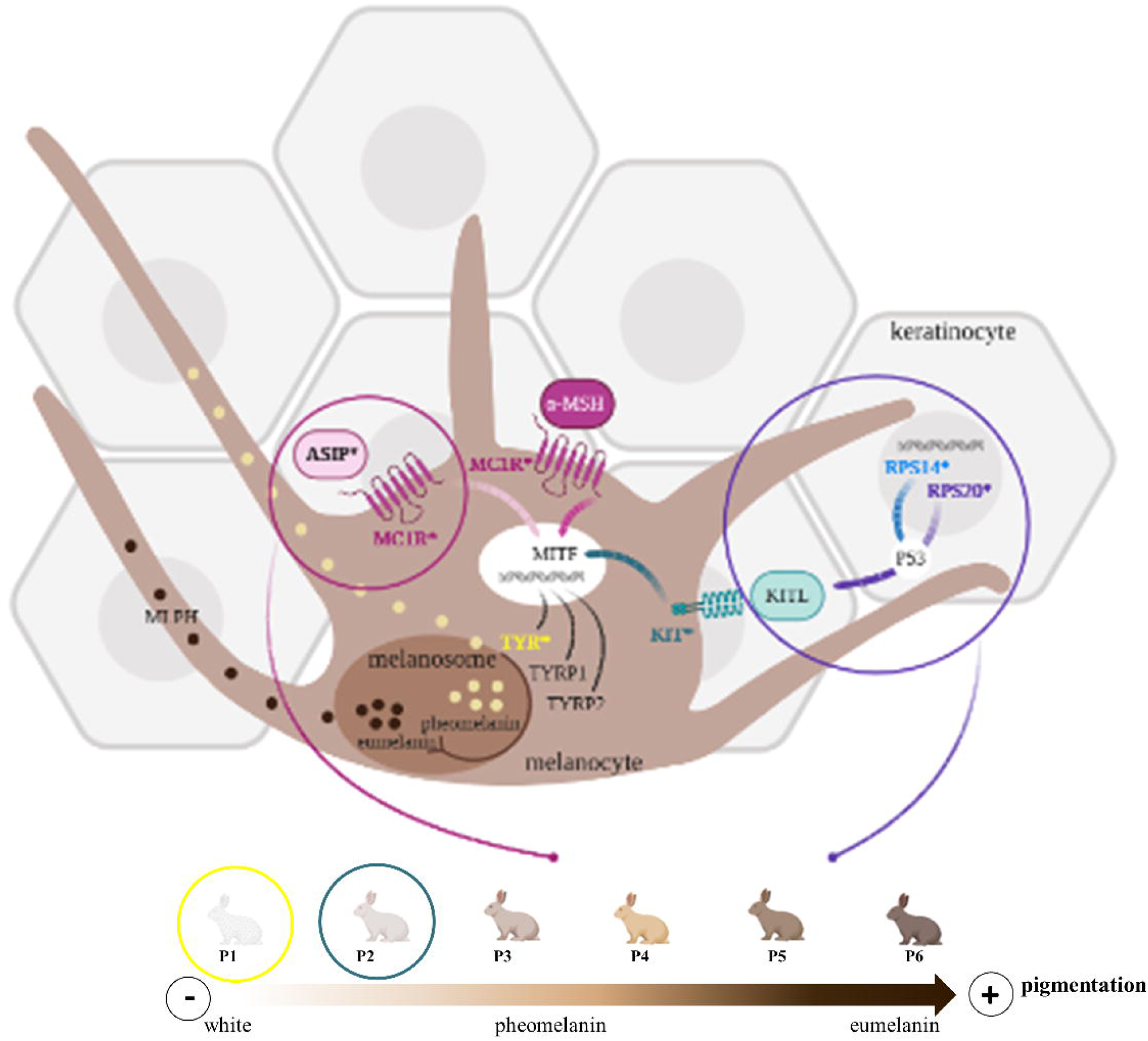
Model of pigmentation molecular architecture through melanogenesis pathway. Both core genes and gene-gene interactions are represented. Core genes identified here to account for coat color variability are in bold, marked with a star and colored in yellow (*TYR*), green (*KIT*), dark pink (*MC1R*), light pink (*ASIP*), purple (*RPS20*) and blue (*RPS14*). Major genes involving *TYR* and *KIT* are responsible of white (P1) (yellow circle) and spotted (P2) (green circle) phenotypes, respectively. The remaining cream to dark brown phenotypes (P3 to P6) are explained by *ASIP, MC1R, TYR* and *RPS* genes as main effects but also through epistatic interactions. While *ASIP*:*MC1R* interactions contribute mainly to light colored rabbits (P3 and P4) (pink circle), *KIT:RPS* (including both *RPS14* and *RPS20*) epistasis seem account for darker colorations through tanning response (purple circle).

The most significant epistatic interaction identified from the various analyses is between both markers tagging *ASIP* and *MC1R* loci. Epistasis involving those genes has already been described in mice [13, 57], human [17], sheep [58] but also in rabbits from a cross between a Champagne d’Argent buck and a Thuringian doe [30]. Interactions have been associated with color variation particularly in Creole sheep where it has clearly been showed from analyses of phenotypes segregation through families crosses that a functional wild-type genotype at *MC1R* locus is needed for the manifestation of the effects of the duplicated allele at the *ASIP* gene [58]. Although highlighting epistasis remains challenging, understanding the functional role of the interaction at the molecular level affecting phenotypes is still more complex. Consequence of *ASIP*:*MC1R* genetic interaction is straightforward since both proteins act within the same signalling pathway with ASIP being an antagonist ligand which competes with α-MSH for binding on its MC1R receptor. Here, significant positive effects from both individual AX-147097074 (Ocu4_*ASIP*_) and AX-147194100 (GL018975_*MC1R*_) markers were identified suggesting likely gain of function mutations. In contrary, epistasis seem to have a negative effect on coat coloration since combination of both variants is mainly associated with light colored rabbits harbouring cream/beige coat pigmentation (P3 and P4 phenotypes) (Fig. 5).

Our results highlighted suggestive interactions between ribosomal processed pseudogenes themselves and *KIT* genomic regions. Although the involvement of ribosomal genes especially *RPS20* and *RPS14* processed pseudogenes needs to be assessed with further experiments, epistasis with those genes and its impact on pigmentation seemed consistent with knowledge [53, 54]. As previously mentioned, deregulation of Rps20 in mice has been shown to activate the Kit signalling pathway through p53 activation mimicking the tanning response responsible of the hyperpigmentation of animals [55] and inactivation of rps14 in zebrafish delayed pigmentation process also via an increase of p53 activity [53]. Our results demonstrating that both individual and combined effects of *RPS* and *KIT* loci affected rabbits with dark coat color of body extremities (P5 and P6 phenotypes) (Fig.5).

Beside epistasis highlighted between the hub genes involved in the determinism of coat coloration, we also pinpointed a denser interaction network including several genes, that are known to affect the pigmentation process. A significant interaction was identified between Ocu4_*ASIP*_ locus and a part of the scaffold GL018733, in which the Heat Shock Protein 5 (*HPS5*) gene is located. Mutations within this gene altered melanosome biogenesis and have been associated with hypopigmentation specific of oculocutaneous albinism [59, 60] and Hermansky-Pudlak Syndrome [60, 61]. The built network also figured out *POT1* (The Protection Of Telomeres 1 protein) and *CERKL*, (Ceramide Kinase Like) located on Ocu7, significantly interacting with GL018965_*MC1R*_ locus. POT1 encodes a nuclear protein involved in telomere maintenance. Several human genetics study carried out in different ethnic groups have characterised mutations responsible of skin melanoma [62–64] making POT1 a major driver of this human disease as reviewed in [65, 66]. The remaining genes that showed epistasis with one of the significant locus (Ocu1_*RPS14*_, Ocu1_*TYR*_, Ocu4_*ASIP*_, and GL018965_*MC1R*_) are retinal pigment related proteins such as SLC7A13 (Solute Carrier Family 7 Member 13) [67, 68]. But more importantly, deregulation of many of them have been associated with retinal pigmentation disorders. Indeed, CERKL [69–71], SPATA7 (Spermatogenesis Associated 7) [72–75] and TTC8 (Tetratricopeptide Repeat Domain 8) [76, 77] have all been involved in retinitis pigmentosa which corresponds to a dysfunction and degeneration of both photoreceptors and retinal pigment endothelial cells.

## Conclusions

To conclude, our results bring new insights into the molecular architecture of the coat color of body extremities pinpointing the key role played by interactions in the establishment of this complex trait. The characterisation of a genome-wide epistatic network might significantly contribute to a better understanding of underlying mechanisms. Moreover, divergences in the relationships between phenotypes and genotypes have been described in different breeds pointing out the functional effect of specific combination of alleles. Future studies through deeper analyses from sequencing data might lead to an allele-specific network considering also their dominance/recessivity or copy numbers.

## Methods

### Animal data

#### The experimental design

The experimental rabbit populations were issued from the INRA 1001 line [78] and bred in the INRAE experimental farm (UE PECTOUL, Toulouse, France) in accordance with the national regulations for animal care and use of animals in agriculture. The experimental population was a combination of two genetically related lines: the G10 line, selected for 10 generations for decreasing Residual Feed Intake (RFI) [79] and the G0 control line produced from frozen embryos of the ancestor population of the selected line. The 296 G10 and 292 G0 rabbits were produced in the same 3 batches with a 42 days interval. In each batch, half of the kits was fostered to G0 does and the second half was fostered by G10 does. Does adopted alternatively kits from one line and from the other line in successive batches. At weaning (32 days), in each batch, kits were placed in individual cages. More details about the experimental cross can be found in Garreau et al. [80] but briefly, the initial design included 832 rabbits including the 20 bucks, 101 does and 711 offspring. Although the experimental cross was not especially designed for evaluating coloration traits, we took advantage of it since the G0 line originated from Californian rabbits and we observed within the experimental cross a segregation for both color and pattern variability within the himalayan phenotype. The final design, based on phenotypic evaluation is detailed further and on Additional file 2: Fig S6.

#### Phenotypic data and quality control

We first distinguished 5 different rabbit color groups (from white to dark chocolate) by visual inspection of the whole population. This notation was performed by 2 independent experimenters. Colors were classified as A, B, C, D and E. Secondly, we selected a few individuals (n=15 per classified group) that were phenotyped for their nose coloration using a colorimeter to validate our subjective classification. A significant correlation was observed between the luminescence (L*) measurement and the notes (Additional file 2: Fig. S6a), validating the determined groups. Moreover, an additional group was created since some animals from class A had colored ears but white noses. Altogether, 6 ordered phenotypes were defined, sorted from P1 to P6 and numbered 1 to 6 for further quantitative analyses (P1=1, P2=2, P3=3, P4=4, P5=5, P6=6). In total, 686 rabbits out of the 832 of the whole experimental design were assigned to one phenotypic group, including 574 offspring, 20 bucks and 92 does, with 2 to 50 offspring per buck and 1 to 15 offspring per doe (Additional file 2: Fig. S6b). Number of rabbits per phenotype were 34, 32, 47, 125, 193 and 255 for P1, P2, P3, P4, P5 and P6, respectively (Additional file 2: Fig. S6c).

#### Sampling collection and DNA extraction

Ear punch biopsies were collected in Allflex Tissue Sampling Unit tube (Allflex France, Vitré, France) and genomic DNA was extracted from samples with a home-made protocole: proteinase K lysis following by salt-based DNA extraction and ethanol precipitation. Briefly, ear punch biopsies were digested at 56°C for 3h using a 500μL solution including 10mM Tris HCl, 0.1M EDTA pH=8, 0,5% SDS and 0.2mg proteinase K. After overnight incubation at 37°C, 1/3 volume of saturated (6M) NaCl were added and slightly mixed before a centrifugation step (30min at 4°C and 21,000g). The supernatant was mixed with 2 volumes of 100% ethanol. DNA was retrieved and resuspended in classic buffer for 1h at 60°C before an overnight resuspension at 37°C. Total genomic DNA was quantified using the Nanodrop 8000 (ND8000LAPTOP, Thermo Fisher Scientific, USA) and the Qubit2.0 (Q32866, Life Technologies, USA).

#### Genotyping data and quality control

The DNA samples were genotyped at the Centro Nacional de Genotipado (CeGen) platform (Santiago de Compostela, Spain) using the Affymetrix® AxiomOrcun™ SNP Array as recommended by the manufacturer. The SNP array contains 199,692 molecular markers spanning both chromosomes and scaffolds. The order of the SNPs was based on the Rabbit OryCun2.0 assembly released by the Broad Institute of MIT and Harvard [81]. Missing data imputation and haplotype phasing were performed with the software FImpute [82]. The SNP data were then filtered based on minor allele frequencies ≥ 0.005 leading to a final SNP dataset of 162,070 markers for association analyses. The other standard filtering were not applied in primo-analyses to not eliminate markers that could pinpoint structural variants. However, the quality (call rate (95%), call freq (95%), Hardy-Weinberg disequilibrium (10^−06^)) of highlighted variants lying within intervals of interest were checked *a posteriori* to secure our results especially for epistasis analyses. Additional 12,640 SNP were excluded for genome-wise epistasis study. Additional manual genotyping of 5 variants, included 4 known mutations (*ASIP* - allele a, *TYR* - alleles c and C^h^ and *MC1R* - allele Ed) and another variant for ASIP - 5435370 bp. Briefly, variants that were a SNP were genotyped either using RFLP PCR (TYR - alleles c and C^h^) or allele-specific PCR (ASIP - 5435370 bp) while variants corresponding to deletions were genotyped using Capillary Electrophoresis (*ASIP* - allele a and *MC1R* - allele Ed and e). Primers and PCR conditions used are presented in Additional file 1: Table S9. Briefly for RFLP PCR, PCR were performed with the kit GoTaq® Flexi (Promega, USA) using 20 ng DNA, 0.5 mM of primers, 0.2 mM dNTPs (Promega, USA), 1X buffer, 1.5mM MgCl2 et 0.25 U Taq in a final volume of 12 μL. Digestions were performed with NciI I et BsaXI (NEB, USA) for TYR - alleles c and C^h^, respectively, using 2U of enzyme and 1X of their respective buffer before incubation at 37°C for 15 min. The PCR and digestion were performed on thermocycleur Verity (Thermo Fisher Scientific, USA) and PCR products were loaded on a 2.5% agarose gel with ethidium bromide. Briefly for allele-specific PCR, we used the KASPAR (Kompetitive Allele Specific PCR) (KBioscience, United Kingdom) technology. Amplification was perfomed with 10 ng DNA, 1X PCR buffer, 1.8 mM MgCl2, 0.2 mM dNTPs, 0.25 μM of each fluorescent dye (Fam et Vic), 0.5 U of Taq polymerase and 12 μM for allele-specific primers and 30 μM for the common primer in a final volume of 5 μL. We followed provider recommendation for the PCR program, fluorescent reading was made on a Quant Studio 6 (Thermo Fisher Scientific, USA) and results were analysed with the software Quant studio Real Time PCR (Thermo Fisher Scientific, USA). Finally, genotyping using capillary electrophoresis were perfomed on a ABI3730™ (Applied Biosystems, USA). The PCR were performed with 0.1 mM of the extended primer, 0.15 mM of the hybridization primer carrying the dye and 0.15 mM of the reverse primer. The other conditions of PCR and cycle are similar to the RFLP PCR. PCR products were loaded on the ABI3730™ after a first step of 1/20 dilution and 2 μL of the dilution were mixed with formamide and size standard GeneScan-600Liz Size Standard (Applied Biosystems, USA) before a denaturation step at 94°C for 5 min. Analyses were perfomed with GeneMapper™ Software (Applied Biosystems, USA).

#### RNA-seq alignments

Publicly available RNA-seq raw data from back skin of rabbits were uploaded to perform alignment, quantification and transcript discovery with statistics. Three Rex rabbits with black or white or chinchilla back skin were considered. Accession numbers for BioSample (https://www.ncbi.nlm.nih.gov/biosample/) are SAMN02693835, SAMN02693836 and SAMN02693834 for black, white and chinchilla, respectively. Accession number for raw data (https://trace.ncbi.nlm.nih.gov/Traces/sra/) are SRR1201255, SRR1201256 and SRR1201257 for black, white and chinchilla, respectively.

Quality controls, alignments and analyses were performed with the open-source nf-core/rnaseq workflow (https://nf-co.re/rnaseq) using the 3.0 version that implemented fastqc 0.11.9 and qualimap 2.2.2 for quality controls and STAR 2.6.1 for the mapping. Paired-end reads were aligned on the reference OryCun2.0 genome and annotation version Oryctolagus_cuniculus.OryCun2.0.104.gtf was used for analyses.

#### Linkage map construction

The netmap option of the netgwas R package (https://www.rdocumentation.org/packages/netgwas/versions/1.13) [83] was used for building a linkage map of the Ocu14 region since 2 distinct intervals away from 20 Mb were within the same credible set.

#### Copy number variations evaluation

The raw measurements consist of two intensity signals, one for each allele, which are subsequently transformed into the log-scaled ratio of the observed and the expected intensity (LRR), and the B Allele Frequency (BAF) which captures the relative contribution from one allele (B) to the fluorescent signal. While expected values of 0 for LRR reflect normal copy number (n=2 for diploid individuals, log_2_(2/2)), aberrant theoretical values of 0.57 or −1 reflect one copy gain (log_2_(3/2)) or loss (log_2_(1/2)), respectively. From BAF values, a BAF value of 0.5 indicates a heterozygous genotype (AB), whereas 0 and 1 indicate homozygous genotypes (AA and BB, respectively). For example, a single copy number gain is characterized by 4 theoretical distinct BAF values = 0, 0.33, 0.67 and 1, reflecting AA/AAA, AAB, ABB and BB/BBB genotypes, respectively. Log R Ratio (LRR) and B Allele Frequency (BAF) were extracted from the Axiom™ Analysis Suite Software 4.0.3.3 (Thermo Fisher Scientific, USA) using the Axiom^®^ CNV Summary Tools 1.1 (Thermo Fisher Scientific, USA) after a global analysis of the whole experimental design. The 8 96-well genotyping plates were analysed simultaneously for an accurate definition of genotypes clusters since no reference cluster exist for the Affymetrix® AxiomOrcun™ SNP Array. However, a large difference in signal intensities between plates for both LRR and BAF values were observed for 2 of them likely affecting the results for CNV analyses. The LRR values were normalised after confirming that it was not a biological effect since the 6 phenotypes and all families were represented on these 2 plates. Results presented here are adjusted LRR values taking the plate effect into account using the lm function in R

### Statistical analyses

#### Univariate linear mixed models for association analyses

We used the GEMMA (Genome-wide Efficient Mixed Model Association) software to perform association analyses. Briefly, GEMMA fits a univariate linear mixed model (LMM) [84] or a Bayesian sparse linear mixed model using Markov chain Monte Carlo (BSLMM) [85]; both methods control for population structure.

SNP effects were tested with the following univariate animal mixed model LMM [84]:

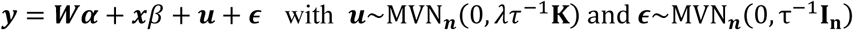

where ***y*** is the vector of phenotypes for a given trait, ***W*** is the incidence matrix of covariates corresponding to fixed effects and ***α*** stands for the effects of these covariates, ***x*** is the vector of allelic dosages of the genotypes (0, 1 or 2) and β stands for marker size effect, ***u*** is the random polygenic effect and ***ϵ*** is the random residual effect. Additive effects are structured after **K**, the centered relatedness matrix computed from the genotypes, *λ* is the ratio between the two variance components and ***τ*** is the variance of the residual errors. GEMMA tests the alternative hypothesis H1: *β* ≠ 0 against the null hypothesis H0: *β* = 0 for each SNP in turn, using one of the three commonly used test statistics (Wald, likelihood ratio or score). In this article, we will only report the p-value associated with the Wald statistic.

In addition to the classical additive model, dominant and recessive models were tested by transforming allelic dosages of the genotypes into binary genotypes based on the minor allele at each SNP. As examples, genotypes coded 0, 1, 2 for a given variant (corresponding to homozygosity for the minor allele, heterozygosity and homozygosity for the major allele, respectively) were recoded either 0, 0, 1 to test for the dominance of the minor allele or 0, 1, 1 to test for the recessivity of the minor allele. Association was concluded as *(i)* significant at genome-wise level after a Bonferroni correction (0.05 / 162,070 = 3.08*10^−07^) and *(ii)* suggestive at chromosome-wise level after a Bonferroni correction (0.05 / n markers on Ocu).

The SNP effects were also tested with the following Bayesian sparse animal mixed model [85]:

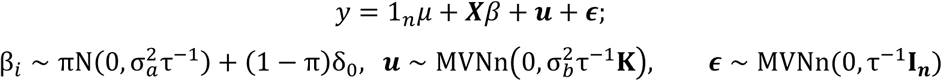

where **1**_***n***_ is a n-vector of 1s, *μ* is a scalar representing the phenotype mean, ***X*** is an n*p matrix of genotypes measured on n individuals at p genetic markers, ***β*** is the corresponding p-vector of the genetic marker effects, and other parameters are the same as defined in the standard linear mixed model. In the special case **K=*XX***^*T*^/*p*, the SNP effect sizes can be decomposed into two parts: **α** that captures the small effects that all SNPs have, and **β** that captures the additional effects of some large effect SNPs. In this case, ***u* = *X*α** can be viewed as the combined effect of all small effects, and the total effect size for a given SNP *i* is *α*_*i*_ *+ β*_*i*_. To pinpoint signals, we summed the sparse probabilities evaluated from the total effect size for a given SNP on sliding windows containing 20 SNPs.

#### Fine-mapping of regions of interest

We also used the SuSiE (Sum of Single Effects) model, which corresponds to a new formulation of the Bayesian variable selection in regression (BVSR) to fine-map the loci [86]. This model fits an Iterative Bayesian Stepwise Selection (IBSS) algorithm that is a Bayesian analogue of traditional stepwise selection methods. SuSie produces Posterior Inclusion Probabibilities (PIPs) and Bayesian Credible Sets (CSs) which capture an effect variable allowing the fine-mapping of significant detected regions. The SuSiE method removes the single causal variant assumption and groups SNPs into distinct association signals in the analysis, such that it aims to find as many CSs of variants that are required so that each set captures an effect variant, whilst also containing as few variants as possible.

Regions showing either -log_10_(p-value) > 6.5 with the LMM (corresponding to 5% genome-wide threshold after a Bonferonni correction) or a PIP > 0.1 were consider for further analyses.

#### Linkage disequilibrium pattern of intervals

Each pairwise linkage disequilibrium measure (r2) was computed both using the PLINK 1.9 software with the --r2 option (www.cog-genomics.org/plink/1.9/) [87] for all the pairs of SNPs of the selected regions.

#### Epistatic interaction analyses

Epistatic interaction analyses were focused on best variants located within selected regions, that were identified from the previously described association analyses.

A first evaluation of epistasis was performed using the linear regression model implemented in PLINK 1.9 [87] with the --epistasis option to fit the model:

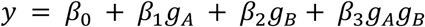

for each inspected variant pair (A, B), where *g*_*A*_ and *g*_*B*_ are allele counts, and *g*_*A*_*g*_*B*_ is the count of common occurrences of the alleles at the two loci; then the *β* coefficients are tested for deviation from zero. Pairwise interaction was tested between each marker of the set of the 7 best associated SNP variants. Interactions with p-value < 0.05 are considered significant.

To determine the best classification of individuals within the 6 phenotypig groups (P1 to P6) given their combined genotypes at the different selected regions, we built a decision tree using the Classification And Regression Trees (CART) algorithm [88]. The evaluation criterion of the CART algorithm is the Gini-index of diversity, which measures how often a random individual in the set would be misranked if its genotypes was randomly assigned according to the distribution of phenotypes in the subset. The Gini-index of diversity can be calculated by summing the probability of each individual being assigned, multiplied by the probability that it would be misranked. It reaches its minimum value (zero) when all individuals in the set are in the same class as the target variable. Moreover, to identify the most likely epistatic interactions between the selected markers, we used the decision criterion BIC metric. The most probable model is therefore the one that minimizes the BIC criterion. The stepAIC function in the MASS package was applied [89].

Pairwise epistatic interactions between the set of selected markers from GWAS and the rest of the genome were also evaluated using the adaptive shrinkage method [31] implemented in the ashr R package (https://www.rdocumentation.org/packages/ashr/versions/2.2-47). This represented approximately (number of selected variants)*200,000 tests. Both interaction effect size and corresponding standard error were thus estimated for each pairwise combination. Those measures were used with an empirical Bayes approach for large-scale hypothesis testing. This method accounts for variation in measurement precision across tests in the computation of the effect sizes, and facilitates their estimation. In addition, instead of p-value, q-value or local FDR, the “Local False Sign Rate” (lfsr) [31], which refers to the probability of getting the sign of an effect wrong, was computed. The authors argue that it is a superior measure of significance than the local FDR:

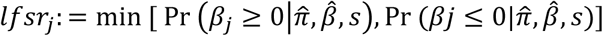

with for effect *j*, lfsr_*j*_, is the probability that we would make an error in the sign of effect *β*_*j*_ if we were forced to declare it either positive or negative. Small values of lfsr_j_ indicate that we can be confident in the sign of *β*_*j*_, which implies that we are confident it has a non-zero value. lfsr is a more conservative measure of significance than local FDR and it is more robust to modeling assumptions.

### Visualisation tools

Several software or R packages were used for visualisation of data. Classical plots were built with the ggplot2 R package (https://www.rdocumentation.org/packages/ggplot2) [90], Manhattan and Quantile-Quantile plots were built using the qqman R package (https://www.rdocumentation.org/packages/qqman) [91]. Decision trees were built with the rpart R package (https://www.rdocumentation.org/packages/rpart) [92]; the Fig. 3b is a concatenation of two independent decision trees. The circular plot was built using the BioCircos R package (https://www.rdocumentation.org/packages/BioCircos) [93]. Linkage disequilibrium profiles were visualized with the LDheatmap R package (https://www.rdocumentation.org/packages/LDheatmap) [94]. Mapping of RNA-seq experiments were showed using the Integrative Genome Viewer (https://software.broadinstitute.org/software/igv/) [95].

## Abbreviations

BAF: B allele frequency
CNV: copy number variation
CS: credible set
GWAS: genome wide association study
LD: linkage disequilibrium
LFSR: local false sign rate
LRR: log R ratio
Ocu: *Oryctolagus cuniculus*
PIP: posterior inclusion probability
RPE: retinal pigment endothelium
SNP: single nucleotide polymorphism

## Declarations

### Ethics approval and consent to participate

The French ministery of higher education, Research and innovation and the local animal research ethics committee (C2EA-115) approved the study (approval number 00903.01). All procedures were conducted in accordance with the French legislation on animal experimentation and ethics. The senior researchers were authorized by the French Ministry of Agriculture to conduct experiments on living animals at the INRAE facilities in Toulouse, France (approval number 312011116).

### Consent for publication

Not applicable

### Availability of data and materials

The datasets (genotypes, phenotypes, pedigrees, LRR values and BAF values) supporting the conclusions of this article are available in the author personal genologin repository (http://genoweb.toulouse.inra.fr/~jdemars/RabbitColoration/) belonging to the Genotoul Bioinformatics facility (http://bioinfo.genotoul.fr/).

### Competing interests

The authors declare that they have no competing interests

### Funding

This study has been funded by the European Union’s H2020 project Feed-a-Gene under grant agreement no. 633531, the Genetic Animal division from INRA, and the GenPhySE laboratory.

### Authors’ contributions

JD analysed data, performed their visualisation and wrote the manuscript, YL performed quality control and analyses, NI organized sampling and genotyping, AD performed additional polymorphisms genotyping, SL performed phenotypic measurements and additional genotyping, HG contribute to statistical analyses and read the manuscript, PA and FB were in charge of animal care and experimental design, JR set up the project, performed phenotypic measurements and quality control of datasets.

## Acknowledgements

We thank all people of the animal facility, who carefully looked after the animals and for their help with the skin biopsies. We warmly thank Laurence Drouilhet, Bertrand Servin and Alain Vignal and all those who have contributed directly or indirectly to this work for discussion and support. We also thank both Centro Nacional de Genotipado (CeGen) (http://www.usc.es/cegen/) and GenoToul bioinformatics (http://bioinfo.genotoul.fr/) facilities.

## Authors’ information (optional)

Not applicable

## Additional files

Additional file 1 corresponds to all supplementary tables

Additional file 2 corresponds to all supplementary figures

## Notes

http://genoweb.toulouse.inra.fr/~jdemars/RabbitColoration/

### Competing Interest Statement

The authors have declared no competing interest.

## References

1. Protas ME, Patel NH (2008) Evolution of coloration patterns. Annu. Rev. Cell Dev. Biol. 24

2. Cieslak M, Reissmann M, Hofreiter M, Ludwig A (2011) Colours of domestication. Biol. Rev. 86

3. Cuthill IC, Allen WL, Arbuckle K, et al (2017) The biology of color. Science (80-.). 357

4. Bennett DC, Lamoreux ML (2003) The color loci of mice - A genetic century. Pigment Cell Res. 16

5. Hoekstra HE (2006) Genetics, development and evolution of adaptive pigmentation in vertebrates. Heredity (Edinb). 97

6. San-Jose LM, Roulin A (2017) Genomics of coloration in natural animal populations. Philos. Trans. R. Soc. B Biol. Sci.

7. Pavan WJ, Sturm RA (2019) The Genetics of Human Skin and Hair Pigmentation. Annu Rev Genomics Hum Genet. https://doi.org/10.1146/annurev-genom-083118-015230

8. Bagnara JT, Taylor JD, Hadley ME (1968) The dermal chromatophore unit. J Cell Biol 38:. https://doi.org/10.1083/jcb.38.1.67

9. Lin JY, Fisher DE (2007) Melanocyte biology and skin pigmentation. Nature

10. Kaelin CB, Xu X, Hong LZ, et al (2012) Specifying and sustaining pigmentation patterns in domestic and wild cats. Science (80-) 337:. https://doi.org/10.1126/science.1220893

11. Pino J, Kos L (2013) MC1R, EDNRB and kit signaling in pigmentation regulation related disorders. In: Skin Pigmentation: Genetics, Geographic Variation and Disorders

12. Duhl DMJ, Vrieling H, Miller KA, et al (1994) Neomorphic agouti mutations in obese yellow mice. Nat Genet. https://doi.org/10.1038/ng0994-59

13. Siracusa LD (1994) The agouti gene: turned on to yellow. Trends Genet.

14. Anthony JF Griffiths, Jeffrey H Miller, David T Suzuki, Richard C Lewontin and WMG (2000) An Introduction to Genetic Analysis, 7th edition

15. Liu F, Visser M, Duffy DL, et al (2015) Genetics of skin color variation in Europeans: genome-wide association studies with functional follow-up. Hum Genet 134:823–835. https://doi.org/10.1007/s00439-015-1559-0

16. Walsh S, Chaitanya L, Breslin K, et al (2017) Global skin colour prediction from DNA. Hum Genet 136:847–863. https://doi.org/10.1007/s00439-017-1808-5

17. Morgan MD, Pairo-Castineira E, Rawlik K, et al (2018) Genome-wide study of hair colour in UK Biobank explains most of the SNP heritability. Nat Commun 9:5271. https://doi.org/10.1038/s41467-018-07691-z

18. Carbone MA, Llopart A, DeAngelis M, et al (2005) Quantitative trait loci affecting the difference in pigmentation between Drosophila yakuba and D. santomea. Genetics 171:. https://doi.org/10.1534/genetics.105.044412

19. Mundy NI (2007) Coloration and the Genetics of Adaptation. PLoS Biol 5:. https://doi.org/10.1371/journal.pbio.0050250

20. Albertson RC, Powder KE, Hu Y, et al (2014) Genetic basis of continuous variation in the levels and modular inheritance of pigmentation in cichlid fishes. Mol Ecol 23:. https://doi.org/10.1111/mec.12900

21. Wollstein A, Walsh S, Liu F, et al (2017) Novel quantitative pigmentation phenotyping enhances genetic association, epistasis, and prediction of human eye colour. Sci Rep 7:. https://doi.org/10.1038/srep43359

22. Ritchie MD, Van Steen K (2018) The search for gene-gene interactions in genome-wide association studies: challenges in abundance of methods, practical considerations, and biological interpretation. Ann Transl Med 6:157. https://doi.org/10.21037/atm.2018.04.05

23. Branicki W, Brudnik U, Wojas-Pelc A (2009) Interactions between HERC2, OCA2 and MC1R may influence human pigmentation phenotype. Ann Hum Genet 73:160–70. https://doi.org/10.1111/j.1469-1809.2009.00504.x

24. Blanchard SG, Harris CO, Ittoop ORR, et al (1995) Agouti Antagonism of Melanocortin Binding and Action in the B16F10 Murine Melanoma Cell Line. Biochemistry. https://doi.org/10.1021/bi00033a012

25. Suzuki I, Tada A, Ollmann MM, et al (1997) Agouti signaling protein inhibits melanogenesis and the response of human melanocytes to α-melanotropin. J Invest Dermatol. https://doi.org/10.1111/1523-1747.ep12292572

26. Hepp D, Gonçalves GL, Moreira GRP, De Freitas TRO (2016) Epistatic interaction of the melanocortin 1 receptor and agouti signaling protein genes modulates wool color in the brazilian creole sheep. J Hered. https://doi.org/10.1093/jhered/esw037

27. Fontanesi L, Scotti E, Allain D, Dall’Olio S (2014) A frameshift mutation in the melanophilin gene causes the dilute coat colour in rabbit (Oryctolagus cuniculus) breeds. Anim Genet. https://doi.org/10.1111/age.12104

28. Utzeri VJ, Ribani A, Fontanesi L (2014) A premature stop codon in the TYRP1 gene is associated with brown coat colour in the European rabbit (Oryctolagus cuniculus). Anim Genet. https://doi.org/10.1111/age.12171

29. Aigner B, Besenfelder U, Müller M, Brem G (2000) Tyrosinase gene variants in different rabbit strains. Mamm Genome 11:700–702. https://doi.org/10.1007/s003350010120

30. Fontanesi L, Forestier L, Allain D, et al (2010) Characterization of the rabbit agouti signaling protein (ASIP) gene: Transcripts and phylogenetic analyses and identification of the causative mutation of the nonagouti black coat colour. Genomics. https://doi.org/10.1016/j.ygeno.2009.11.003

31. Stephens M (2017) False discovery rates: A new deal. Biostatistics. https://doi.org/10.1093/biostatistics/kxw041

32. Paterson EK, Fielder TJ, MacGregor GR, et al (2015) Tyrosinase depletion prevents the maturation of melanosomes in the mouse hair follicle. PLoS One 10:. https://doi.org/10.1371/journal.pone.0143702

33. Cooper MP, Fretwell N, Bailey SJ, Lyons LA (2006) White spotting in the domestic cat (Felis catus) maps near KIT on feline chromosome B1. Anim Genet 37:163–165. https://doi.org/10.1111/j.1365-2052.2005.01389.x

34. Haase B, Rieder S, Leeb T (2015) Two variants in the KIT gene as candidate causative mutations for a dominant white and a white spotting phenotype in the donkey. Anim Genet 46:321–324. https://doi.org/10.1111/age.12282

35. Holl H, Isaza R, Mohamoud Y, et al (2017) A frameshift mutation in KIT is associated with white spotting in the Arabian camel. Genes (Basel) 8:. https://doi.org/10.3390/genes8030102

36. Hauswirth R, Jude R, Haase B, et al (2013) Novel variants in the KIT and PAX3 genes in horses with white-spotted coat colour phenotypes. Anim Genet 44:763–765. https://doi.org/10.1111/age.12057

37. Fontanesi L, Vargiolu M, Scotti E, et al (2014) The kit gene is associated with the english spotting coat color locus and congenital megacolon in checkered giant rabbits (oryctolagus cuniculus). PLoS One 9:. https://doi.org/10.1371/journal.pone.0093750

38. Dürig N, Jude R, Holl H, et al (2017) Whole genome sequencing reveals a novel deletion variant in the KIT gene in horses with white spotted coat colour phenotypes. Anim Genet 48:483–485. https://doi.org/10.1111/age.12556

39. Silvers WK, Silvers WK (1979) The Agouti and Extension Series of Alleles, Umbrous, and Sable. In: The Coat Colors of Mice

40. Norris BJ, Whan VA (2008) A gene duplication affecting expression of the ovine ASIP gene is responsible for white and black sheep. Genome Res. https://doi.org/10.1101/gr.072090.107

41. Robic A, Morisson M, Leroux S, et al (2019) Two new structural mutations in the 5′ region of the ASIP gene cause diluted feather color phenotypes in Japanese quail. Genet Sel Evol. https://doi.org/10.1186/s12711-019-0458-6

42. Letko A, Ammann B, Jagannathan V, et al (2020) A deletion spanning the promoter and first exon of the hair cycle-specific ASIP transcript isoform in black and tan rabbits. Anim Genet 51:. https://doi.org/10.1111/age.12881

43. Henkel J, Saif R, Jagannathan V, et al (2019) Selection signatures in goats reveal copy number variants underlying breed-defining coat color phenotypes. PLOS Genet 15:e1008536. https://doi.org/10.1371/journal.pgen.1008536

44. Fontanesi L, Tazzoli M, Beretti F, Russo V (2006) Mutations in the melanocortin 1 receptor (MC1R) gene are associated with coat colours in the domestic rabbit (Oryctolagus cuniculus). Anim Genet. https://doi.org/10.1111/j.1365-2052.2006.01494.x

45. Fontanesi L, Scotti E, Colombo M, et al (2010) A composite six bp in-frame deletion in the melanocortin 1 receptor (MC1R) gene is associated with the Japanese brindling coat colour in rabbits (Oryctolagus cuniculus). BMC Genet. https://doi.org/10.1186/1471-2156-11-59

46. Gross JB, Weagley J, Stahl BA, et al (2018) A local duplication of the Melanocortin receptor 1 locus in Astyanax. Genome 61:. https://doi.org/10.1139/gen-2017-0049

47. Zhang Z, Harrison P, Gerstein M (2002) Identification and analysis of over 2000 ribosomal protein pseudogenes in the human genome. Genome Res 12:1466–1482. https://doi.org/10.1101/gr.331902

48. Balasubramanian S, Zheng D, Liu YJ, et al (2009) Comparative analysis of processed ribosomal protein pseudogenes in four mammalian genomes. Genome Biol 10:R2. https://doi.org/10.1186/gb-2009-10-1-r2

49. Balasubramanian S, Zheng D, Liu YJ, et al (2009) Comparative analysis of processed ribosomal protein pseudogenes in four mammalian genomes. Genome Biol 10:. https://doi.org/10.1186/gb-2009-10-1-r2

50. Tonner P, Srinivasasainagendra V, Zhang S, Zhi D (2012) Detecting transcription of ribosomal protein pseudogenes in diverse human tissues from RNA-seq data. BMC Genomics 13:. https://doi.org/10.1186/1471-2164-13-412

51. Tonner P, Srinivasasainagendra V, Zhang S, Zhi D (2012) Detecting transcription of ribosomal protein pseudogenes in diverse human tissues from RNA-seq data. BMC Genomics 13:412. https://doi.org/10.1186/1471-2164-13-412

52. López S, Smith-Zubiaga I, De Galdeano AG, et al (2015) Comparison of the transcriptional profiles of melanocytes from dark and light skinned individuals under basal conditions and following ultraviolet-B irradiation. PLoS One 10:. https://doi.org/10.1371/journal.pone.0134911

53. Ear J, Hsueh J, Nguyen M, et al (2016) A Zebrafish Model of 5q-Syndrome Using CRISPR/Cas9 Targeting RPS14 Reveals a p53-Independent and p53-Dependent Mechanism of Erythroid Failure. J Genet Genomics 43:307–318. https://doi.org/10.1016/j.jgg.2016.03.007

54. McGowan KA, Li JZ, Park CY, et al (2008) Ribosomal mutations cause p53-mediated dark skin and pleiotropic effects. Nat Genet 40:963–970. https://doi.org/10.1038/ng.188

55. Walker G, Box N (2008) Ribosomal stress, p53 activation and the tanning response. Expert Rev Dermatol 3:649–656. https://doi.org/10.1586/17469872.3.6.649

56. Arnheiter H, Bharti K (2008) Ribosomes and p53 - a new KIT for skin darkening. Pigment Cell Melanoma Res 21:501–502. https://doi.org/10.1111/j.1755-148X.2008.00488.x

57. Steiner CC, Weber JN, Hoekstra HE (2008) Erratum: Adaptive variation in beach mice produced by two interacting pigmentation genes (PloS Biology (2007) 5, 9, DOI: 10.1371/journal.pbio.0050219). PLoS Biol. 6:0418

58. Hepp D, Gonçalves GL, Moreira GRP, de Freitas TRO (2016) Epistatic Interaction of the Melanocortin 1 Receptor and Agouti Signaling Protein Genes Modulates Wool Color in the Brazilian Creole Sheep. J Hered 107:544–552. https://doi.org/10.1093/jhered/esw037

59. Okamura K, Suzuki T (2020) Current landscape of Oculocutaneous Albinism in Japan. Pigment Cell Melanoma Res.

60. Mériot M, Hitte C, Rimbault M, et al (2020) Donskoy cats as a new model of oculocutaneous albinism with the identification of a splice-site variant in Hermansky–Pudlak Syndrome 5 gene. Pigment Cell Melanoma Res 33:814–825. https://doi.org/10.1111/pcmr.12906

61. Daly CMS, Willer J, Gregg R, Gross JM (2013) Snow white, a model of Hermansky-Pudlak syzndrome type 5. Genetics 195:481–494. https://doi.org/10.1534/genetics.113.154898

62. Shi J, Yang XR, Ballew B, et al (2014) Rare missense variants in POT1 predispose to familial cutaneous malignant melanoma. Nat Genet 46:482–486. https://doi.org/10.1038/ng.2941

63. Robles-Espinoza CD, Harland M, Ramsay AJ, et al (2014) POT1 loss-of-function variants predispose to familial melanoma. Nat Genet 46:478–481. https://doi.org/10.1038/ng.2947

64. Trigueros-Motos L (2014) Mutations in POT1 predispose to familial cutaneous malignant melanoma. Clin Genet 86:217–218. https://doi.org/10.1111/cge.12416

65. Larue L (2020) Centenary theme section: SKIN MALIGNANCIES SIGNIFICANCE. https://doi.org/10.2340/00015555-3494

66. Newton-Bishop JA, Bishop DT, Harland M (2020) Melanoma genomics. Acta Derm. Venereol. 100:266–271

67. Sarthy VP, Pignataro L, Pannicke T, et al (2005) Glutamate transport by retinal Müller cells in glutamate/aspartate transporter-knockout mice. Glia 49:184–196. https://doi.org/10.1002/glia.20097

68. Chintala S, Li W, Lamoreux ML, et al (2005) Slc7a11 gene controls production of pheomelanin pigment and proliferation of cultured cells. Proc Natl Acad Sci U S A 102:10964–10969. https://doi.org/10.1073/pnas.0502856102

69. Tuson M, Marfany G, Gonzàlez-Duarte R (2004) Mutation of CERKL, a Novel Human Ceramide Kinase Gene, Causes Autosomal Recessive Retinitis Pigmentosa (RP26). Am J Hum Genet 74:128–138. https://doi.org/10.1086/381055

70. Aleman TS, Soumittra N, Cideciyan A V., et al (2009) CERKL mutations cause an autosomal recessive cone-rod dystrophy with inner retinopathy. Investig Ophthalmol Vis Sci 50:5944–5954. https://doi.org/10.1167/iovs.09-3982

71. Avila-Fernandez A, Riveiro-Alvarez R, Vallespin E, et al (2008) CERKL mutations and associated phenotypes in seven Spanish families with autosomal recessive retinitis pigmentosa. Investig Ophthalmol Vis Sci 49:2709–2713. https://doi.org/10.1167/iovs.07-0865

72. Matsui R, McGuigan DB, Gruzensky ML, et al (2016) SPATA7: Evolving phenotype from cone-rod dystrophy to retinitis pigmentosa. Ophthalmic Genet 37:333–338. https://doi.org/10.3109/13816810.2015.1130154

73. Eblimit A, Agrawal SA, Thomas K, et al (2018) Conditional loss of Spata7 in photoreceptors causes progressive retinal degeneration in mice. Exp Eye Res 166:120–130. https://doi.org/10.1016/j.exer.2017.10.015

74. Sengillo JD, Lee W, Bilancia CG, et al (2018) Phenotypic expansion and progression of SPATA7-associated retinitis pigmentosa. Doc Ophthalmol 136:125–133. https://doi.org/10.1007/s10633-018-9626-1

75. Salmaninejad A, Bedoni N, Ravesh Z, et al (2020) Whole exome sequencing and homozygosity mapping reveals genetic defects in consanguineous Iranian families with inherited retinal dystrophies. Sci Rep 10:. https://doi.org/10.1038/s41598-020-75841-9

76. Riazuddin SA, Iqbal M, Wang Y, et al (2010) A Splice-Site Mutation in a Retina-Specific Exon of BBS8 Causes Nonsyndromic Retinitis Pigmentosa. Am J Hum Genet 86:805–812. https://doi.org/10.1016/j.ajhg.2010.04.001

77. Downs LM, Wallin-Håkansson B, Bergström T, Mellersh CS (2014) A novel mutation in TTC8 is associated with progressive retinal atrophy in the golden retriever. Canine Genet Epidemiol 1:4. https://doi.org/10.1186/2052-6687-1-4

78. Larzul C, De Rochambeau H (2005) Selection for residual feed consumption in the rabbit. Livest Prod Sci. https://doi.org/10.1016/j.livprodsci.2004.12.007

79. Drouilhet L, Gilbert H, Balmisse E, et al (2013) Genetic parameters for two selection criteria for feed efficiency in rabbits. J Anim Sci. https://doi.org/10.2527/jas.2012-6176

80. Garreau H, Ruesche J, Gilbert H, et al (2019) Estimating direct genetic and maternal effects affecting rabbit growth and feed efficiency with a factorial design. J Anim Breed Genet. https://doi.org/10.1111/jbg.12380

81. Miguel C, Carl-Johan R, Palma D, et al (2014) Rabbit genome analysis reveals a polygenic basis for phenotypic change during domestication. Science (80-)

82. Sargolzaei M, Chesnais JP, Schenkel FS (2014) A new approach for efficient genotype imputation using information from relatives. BMC Genomics. https://doi.org/10.1186/1471-2164-15-478

83. Behrouzi P, Wit EC (2019) De novo construction of polyploid linkage maps using discrete graphical models. Bioinformatics. https://doi.org/10.1093/bioinformatics/bty777

84. Zhou X, Stephens M (2012) Genome-wide efficient mixed-model analysis for association studies. Nat Genet. https://doi.org/10.1038/ng.2310

85. Zhou X, Carbonetto P, Stephens M (2013) Polygenic Modeling with Bayesian Sparse Linear Mixed Models. PLoS Genet. https://doi.org/10.1371/journal.pgen.1003264

86. Wang G, Sarkar A, Carbonetto P, Stephens M (2020) A simple new approach to variable selection in regression, with application to genetic fine mapping. J R Stat Soc Ser B Stat Methodol. https://doi.org/10.1111/rssb.12388

87. Chang CC, Chow CC, Tellier LC, et al (2015) Second-generation PLINK: rising to the challenge of larger and richer datasets. Gigascience 4:7. https://doi.org/10.1186/s13742-015-0047-8

88. Breiman L, Friedman J, Stone C, Olshen R (1984) Classification and Regression Trees (Wadsworth Statistics/Probability). New York CRC Press

89. Venables WN, Ripley BD (2002) Modern Applied Statistics with S Fourth edition by

90. Wickham H (2016) ggplot2 Elegant Graphics for Data Analysis (Use R!)

91. D. Turner S (2018) qqman: an R package for visualizing GWAS results using Q-Q and manhattan plots. J Open Source Softw. https://doi.org/10.21105/joss.00731

92. Therneau T, Atkinson B, Ripley B (2015) Package ‘rpart’

93. Cui Y, Chen X, Luo H, et al (2016) BioCircos.js: An interactive Circos JavaScript library for biological data visualization on web applications. Bioinformatics. https://doi.org/10.1093/bioinformatics/btw041

94. Shin J-H, Blay S, Graham J, McNeney B (2006) LDheatmap: An R Function for Graphical Display of Pairwise Linkage Disequilibria Between Single Nucleotide Polymorphisms. J Stat Softw. https://doi.org/10.18637/jss.v016.c03

95. Thorvaldsdóttir H, Robinson JT, Mesirov JP (2013) Integrative Genomics Viewer (IGV): High-performance genomics data visualization and exploration. Brief Bioinform 14:. https://doi.org/10.1093/bib/bbs017

