## Additional Tables for "A genome-wide epistatic network underlies the molecular architecture of continuous color variation of body extremities: a rabbit model"

**Table S1: Best SNP marker associated with extremities coat coloration in rabbits** **2**
**Table S2: Segregation of known mutations within the experimental design** **3**
**Table S3: Fine mapping of regions involved in the determinism of extremities colored rabbits** **4**
**Table S4: Linkage map of both regions of the credible set located on Ocu14** **5**
**Table S5: Pairwise epistatic interaction between the set of best associated SNP markers** **6**
**Table S6: Significant epistatic interaction with the best SNP marker of the Ocu1$_{RPS14}$ locus** **7**
**Table S7: Significant epistatic interaction with the best SNP marker of the Ocu1$_{TYR}$ locus** **9**
**Table S8: Significant epistatic interaction with the best SNP marker of the Ocu4$_{ASIP}$ locus** **10**
**Table S9: Significant epistatic interaction with the best SNP marker of the GL018965$_{MC1R}$ locus** **14**
**Table S10: Primers and PCR conditions for additional manual genotyping** **20**

**Table S1: Best SNP marker associated with extremities coat coloration in rabbits**
The whole experimental design considering all animals from white (P1=1) to dark brown (P6=6) was analysed using a linear mixed model under an additive determinism assumption.

| Marker | Chromosome | Position | p-value |
|---|---|---|---|
| AX-146986391 | 1 | 125,766,001 | 2.36E-56 |
| AX-147059932 | 3 | 131,847,470 | 9.70E-06 |
| AX-147169681 | 4 | 7,186,175 | 4.84E-06 |
| AX-146983797 | 15 | 93,913,201 | 6.80E-11 |
| AX-147179313 | GL018754 | 18,452 | 2.83E-06 |
| AX-147173908 | GL018965 | 86,908 | 5.49E-05 |

**Table S2: Segregation of known mutations within the experimental design**

Genotypic distribution of known coloration mutations with genotypes coded 0/0, 0/1 and 1/1, 0 and 1 being major and minor alleles, respectively. While 6 out of 7 variants were present on the Affymetrix® AxiomOrcun™ SNP Array, the ASIP-agouti mutation was manually genotyped.

| | | **P1 (n=34)** | **P2 (n=32)** | **P3 (n=47)** | **P4 (n=125)** | **P5 (n=193)** | **P6 (n=255)** |
|---|---|---|---|---|---|---|---|
| Ocu1 | 0/0 | 34 | 31 | 43 | 122 | 191 | 250 |
| AX-146983952 41360196 | 0/1 | 0 | 1 | 4 | 3 | 2 | 5 |
| *TYRP1_brown* | 1/1 | 0 | 0 | 0 | 0 | 0 | 0 |
| Ocu1 | 0/0 | 0 | 18 | 14 | 46 | 102 | 234 |
| AX-146982536 127636997 | 0/1 | 4 | 14 | 33 | 79 | 91 | 21 |
| *TYR_albino* | 1/1 | 30 | 0 | 0 | 0 | 0 | 0 |
| Ocu1 | 0/0 | 0 | 18 | 13 | 45 | 104 | 237 |
| AX-146983594 127650960 | 0/1 | 4 | 14 | 34 | 80 | 89 | 18 |
| *TYR_himalayan* | 1/1 | 30 | 0 | 0 | 0 | 0 | 0 |
| Ocu4 | 0/0 | 15 | 14 | 13 | 58 | 122 | 182 |
| 5435400 | 0/1 | 15 | 12 | 28 | 55 | 61 | 62 |
| *ASIP_agouti* | 1/1 | 4 | 6 | 6 | 12 | 10 | 11 |
| GL018840 | 0/0 | 33 | 31 | 44 | 120 | 190 | 253 |
| AX-146983586 549853 | 0/1 | 1 | 1 | 3 | 5 | 3 | 1 |
| *MLPH_dilute* | 1/1 | 0 | 0 | 0 | 0 | 0 | 1 |
| GL018965 | 0/0 | 26 | 25 | 17 | 87 | 166 | 233 |
| AX-147194100 28267 | 0/1 | 7 | 3 | 28 | 38 | 27 | 20 |
| *MC1R_black* | 1/1 | 1 | 4 | 2 | 0 | 0 | 2 |
| GL018965 | 0/0 | 34 | 32 | 47 | 125 | 193 | 253 |
| AX-147194101 28268 | 0/1 | 0 | 0 | 0 | 0 | 0 | 1 |
| *MC1R_japanese* | 1/1 | 0 | 0 | 0 | 0 | 0 | 1 |

**Table S3: Fine mapping of regions involved in the determinism of extremities colored rabbits**

Only individuals carrying coloration at extremities, from cream/beige (P3) to dark chocolate/brown (P6) were considered and analysed using a Bayesian method which captures an effect variable allowing the fine-mapping of significant detected regions that are called Credible Set

| | | **Credible Set** | | **Best-Marker** | |
|---|---|---|---|---|---|
| **Name** | **Location** | **Position (bp)** | **Size (bp)** | **Marker** | **Position (bp)** |
| Ocu1 $_{RPS14}$ | Ocu1 | 60,434,668 | | AX-146995791 | 60,434,668 |
| Ocu1 $_{TYR}$ | Ocu1 | 125,705,538-127,952,493 | 2,246,955 | AX-147073566 | 126,663,954 |
| Ocu4 $_{ASIP}$ | Ocu4 | 5,429,684-7,354,537 | 1,924,853 | AX-147097074 | 6,334,054 |
| Ocu14 $_{RPS20}$ | Ocu14 | 4,983,223-5,568,243 and 28,770,644-29,027,099 | 585,020 + 256,455 | AX-147006836 and AX-147162382 | 5,217,509 and 28,926,645 |
| GL018965 $_{MC1R}$ | GL018965-GL018998 | 18,824-219,701 and 27,411-256,343 | 200,877 + 228,932 | AX-147194100 and AX-147082818 | 152,612 and 77,688 |

**<u>Table S4</u>: Linkage map of both regions of the credible set located on Ocu14**

| SNP | Position (cM) | Position on Ocu14 (bp) |
|---|---|---|
| AX-147102187 | 0.000000 | 5,515,735 |
| AX-147104064 | 0.143886 | 5,383,396 |
| AX-146982543 | 1.013474 | 28,895,431 |
| AX-147117705 | 1.013475 | 29,027,099 |
| AX-146998270 | 2.029476 | 5,553,685 |
| AX-147120707 | 3.045477 | 28,876,218 |
| AX-147059043 | 3.915066 | 4,996,898 |
| AX-147156329 | 4.78465 | 28,850,566 |
| AX-146983123 | 4.784655 | 28,904,781 |
| AX-147094312 | 5.654243 | 5,286,756 |
| AX-146994017 | 5.798129 | 5,501,834 |
| AX-146983467 | 6.814130 | 28,898,875 |
| AX-147012064 | 7.683718 | 5,224,952 |
| AX-147155811 | 8.553306 | 28,889,346 |
| AX-147132312 | 9.569307 | 5,471,323 |
| **AX-147006836** | 10.292910 | 5,217,509 |
| AX-146983562 | 11.750740 | 28,992,704 |
| AX-147180319 | 12.913584 | 5,292,788 |
| AX-147139656 | 13.201770 | 5,246,126 |
| AX-147069141 | 13.345656 | 5,568,243 |
| AX-147008516 | 14.361658 | 28,930,213 |
| AX-146990107 | 15.231246 | 5,043,566 |
| AX-147029148 | 16.100834 | 28,770,644 |
| AX-147178304 | 16.100835 | 28,996,427 |
| AX-147002203 | 16.678877 | 5,253,674 |
| AX-147005242 | 17.256919 | 28,871,253 |
| AX-147117704 | 17.256920 | 28,904,706 |
| AX-147033896 | 18.126508 | 5,330,354 |
| AX-147031484 | 18.126509 | 5,032,093 |
| AX-147169547 | 18.996097 | 28,937,642 |
| AX-147152923 | 18.996098 | 28,934,363 |
| AX-147158867 | 18.996099 | 28,854,954 |
| AX-147164764 | 18.996100 | 28,971,556 |
| AX-147061540 | 19.865689 | 4,983,223 |
| AX-147135673 | 20.735277 | 28,979,381 |
| AX-147098370 | 20.735278 | 28,909,079 |
| AX-147081143 | 20.735279 | 29,012,593 |
| AX-147059044 | 21.751280 | 5,462,683 |
| AX-147029125 | 21.895166 | 5,304,504 |

**Table S5: Pairwise epistatic interaction between the set of best associated SNP markers**

The whole experimental design considering all animals from white (P1) to dark brown (P6) was considered. Pairwise epistasis between the set of the 7 best associated SNP markers was evaluated using a linear regression model. beta int: regression coefficient, stat: chi-square statistic and pval: chi-square p-value, Un : chromosome Unknown. Significant interactions (p-value < 0.05) are in bold while suggestive interaction are in italic.

| Locus1 | SNP1 | Locus2 | SNP2 | Beta int | stat | p-value |
|---|---|---|---|---|---|---|
| Ocu1$_{RPS14}$ | **AX-146995791** | **1** | **AX-147087415** | **0.762039** | **5.85968** | **0.01549** |
| | **AX-146995791** | **1** | **AX-147073566** | **0.899964** | **6.51296** | **0.01071** |
| | AX-146995791 | 4 | AX-147097074 | 0.247889 | 0.188675 | 0.664 |
| | AX-146995791 | 14 | AX-147006836 | -0.199664 | 0.176629 | 0.6743 |
| | AX-146995791 | 15 | AX-146983797 | 0.230966 | 0.471242 | 0.4924 |
| | AX-146995791 | Un | AX-147194100 | 0.208486 | 0.19114 | 0.662 |
| Ocu1$_{TYR}$ | **AX-147087415** | **1** | **AX-146995791** | **0.762039** | **5.85968** | **0.01549** |
| | **AX-147087415** | **1** | **AX-147073566** | **-1.09457** | **82.444** | **1.103e-19** |
| | AX-147087415 | 4 | AX-147097074 | -0.00269295 | 0.000262619 | 0.9871 |
| | AX-147087415 | 14 | AX-147006836 | -0.131134 | 1.16329 | 0.2808 |
| | AX-147087415 | 15 | AX-146983797 | 0.131269 | 1.21629 | 0.2701 |
| | AX-147087415 | Un | AX-147194100 | 0.120022 | 0.607097 | 0.4359 |
| Ocu1$_{TYR}$ | **AX-147073566** | **1** | **AX-146995791** | **0.899964** | **6.51296** | **0.01071** |
| | **AX-147073566** | **1** | **AX-147087415** | **-1.09457** | **82.444** | **1.103e-19** |
| | AX-147073566 | 4 | AX-147097074 | -0.037269 | 0.0456237 | 0.8309 |
| | AX-147073566 | 14 | AX-147006836 | -0.151838 | 1.46439 | 0.2262 |
| | AX-147073566 | 15 | AX-146983797 | 0.131301 | 1.14583 | 0.2844 |
| | AX-147073566 | Un | AX-147194100 | 0.108285 | 0.473117 | 0.4916 |
| Ocu4$_{ASIP}$ | AX-147097074 | 1 | AX-146995791 | 0.247889 | 0.188675 | 0.664 |
| | AX-147097074 | 1 | AX-147087415 | -0.00269295 | 0.000262619 | 0.9871 |
| | AX-147097074 | 1 | AX-147073566 | -0.037269 | 0.0456237 | 0.8309 |
| | *AX-147097074* | *14* | *AX-147006836* | *0.38897* | *3.03937* | *0.08127* |
| | AX-147097074 | 15 | AX-146983797 | -0.0308148 | 0.0241143 | 0.8766 |
| | *AX-147097074* | *Un* | *AX-147194100* | *-0.400949* | *2.61836* | *0.1056* |
| Ocu14$_{RPS20}$ | AX-147006836 | 1 | AX-146995791 | -0.199664 | 0.176629 | 0.6743 |
| | AX-147006836 | 1 | AX-147087415 | -0.131134 | 1.16329 | 0.2808 |
| | AX-147006836 | 1 | AX-147073566 | -0.151838 | 1.46439 | 0.2262 |
| | *AX-147006836* | *4* | *AX-147097074* | *0.38897* | *3.03937* | *0.08127* |
| | **AX-147006836** | **15** | **AX-146983797** | **0.399583** | **5.84738** | **0.0156** |
| | AX-147006836 | Un | AX-147194100 | -0.193767 | 0.90847 | 0.3405 |
| Ocu15$_{KIT}$ | AX-146983797 | 1 | AX-146995791 | 0.230966 | 0.471242 | 0.4924 |
| | AX-146983797 | 1 | AX-147087415 | 0.131269 | 1.21629 | 0.2701 |
| | AX-146983797 | 1 | AX-147073566 | 0.131301 | 1.14583 | 0.2844 |
| | AX-146983797 | 4 | AX-147097074 | -0.0308148 | 0.0241143 | 0.8766 |
| | **AX-146983797** | **14** | **AX-147006836** | **0.399583** | **5.84738** | **0.0156** |
| | AX-146983797 | Un | AX-147194100 | 0.160126 | 0.562804 | 0.4531 |
| GL018965$_{MC1R}$ | AX-147194100 | 1 | AX-146995791 | 0.208486 | 0.19114 | 0.662 |
| | AX-147194100 | 1 | AX-147087415 | 0.120022 | 0.607097 | 0.4359 |
| | AX-147194100 | 1 | AX-147073566 | 0.108285 | 0.473117 | 0.4916 |
| | *AX-147194100* | *4* | *AX-147097074* | *-0.400949* | *2.61836* | *0.1056* |
| | AX-147194100 | 14 | AX-147006836 | -0.193767 | 0.90847 | 0.3405 |
| | AX-147194100 | 15 | AX-146983797 | 0.160126 | 0.562804 | 0.4531 |

**Table S6: Significant epistatic interaction with the best SNP marker of the Ocu1$_{RPS14}$ locus**

Only individuals carrying coloration at extremities, from cream/beige (P3) to dark chocolate/brown (P6) were considered and analysed using an adaptive shrinkage method. lfsr: local false sign rate, lfdr: local false discovery rate, beta: interaction effect and se: standard error

| SNP1 | Location | SNP2 | Ocu | Position | Gene | lfsr | lfdr | pval | beta | se |
|---|---|---|---|---|---|---|---|---|---|---|
| AX-146995791 | Ocu1$_{RPS14}$ | AX-147013489 | 1 | 38213144 | | 5.4233E-04 | 2.0790E-04 | 2.3913e-7 | -0.9611 | 0.1839 |
| AX-146995791 | Ocu1$_{RPS14}$ | AX-147103631 | 1 | 38218436 | | 5.1058E-04 | 1.9605E-04 | 2.2139e-7 | 0.9632 | 0.1838 |
| AX-146995791 | Ocu1$_{RPS14}$ | AX-147048245 | 1 | 38259888 | | 4.6036E-05 | 1.9016E-05 | 1.4948e-8 | -0.9926 | 0.1729 |
| AX-146995791 | Ocu1$_{RPS14}$ | AX-147147565 | 1 | 38267896 | | 5.2189E-04 | 2.0023E-04 | 2.2679e-7 | -0.9631 | 0.1839 |
| AX-146995791 | Ocu1$_{RPS14}$ | AX-147091738 | 1 | 38302698 | | 5.3941E-04 | 2.0679E-04 | 2.3709e-7 | -0.9616 | 0.1839 |
| AX-146995791 | Ocu1$_{RPS14}$ | AX-147087343 | 1 | 38324143 | | 5.4233E-04 | 2.0790E-04 | 2.3913e-7 | -0.9611 | 0.1839 |
| AX-146995791 | Ocu1$_{RPS14}$ | AX-147125939 | 1 | 38358432 | | 5.5104E-04 | 2.1115E-04 | 2.4414e-7 | -0.9604 | 0.1839 |
| AX-146995791 | Ocu1$_{RPS14}$ | AX-146999150 | 1 | 38386484 | | 5.4974E-04 | 2.1066E-04 | 2.4332e-7 | -0.9606 | 0.1839 |
| AX-146995791 | Ocu1$_{RPS14}$ | AX-147073489 | 1 | 38402700 | | 5.8871E-04 | 2.2519E-04 | 2.6609e-7 | -0.9579 | 0.1840 |
| AX-146995791 | Ocu1$_{RPS14}$ | AX-147008077 | 1 | 38431439 | | 5.4233E-04 | 2.0790E-04 | 2.3913e-7 | -0.9611 | 0.1839 |
| AX-146995791 | Ocu1$_{RPS14}$ | AX-147050817 | 1 | 38441877 | | 5.4233E-04 | 2.0790E-04 | 2.3913e-7 | -0.9611 | 0.1839 |
| AX-146995791 | Ocu1$_{RPS14}$ | AX-147068576 | 1 | 38456467 | | 5.4233E-04 | 2.0790E-04 | 2.3913e-7 | -0.9611 | 0.1839 |
| AX-146995791 | Ocu1$_{RPS14}$ | AX-146993778 | 1 | 38534949 | | 5.4233E-04 | 2.0790E-04 | 2.3913e-7 | -0.9611 | 0.1839 |
| AX-146995791 | Ocu1$_{RPS14}$ | AX-147030909 | 1 | 38564408 | | 5.4233E-04 | 2.0790E-04 | 2.3913e-7 | -0.9611 | 0.1839 |
| AX-146995791 | Ocu1$_{RPS14}$ | AX-147071051 | 1 | 38572373 | | 5.4233E-04 | 2.0790E-04 | 2.3913e-7 | -0.9611 | 0.1839 |
| AX-146995791 | Ocu1$_{RPS14}$ | AX-147008078 | 1 | 38578041 | | 4.5580E-04 | 1.7564E-04 | 1.9317e-7 | 0.9656 | 0.1833 |
| AX-146995791 | Ocu1$_{RPS14}$ | AX-147068577 | 1 | 38611851 | | 4.5580E-04 | 1.7564E-04 | 1.9317e-7 | 0.9656 | 0.1833 |
| AX-146995791 | Ocu1$_{RPS14}$ | AX-147089557 | 1 | 38727648 | | 5.3507E-04 | 2.0520E-04 | 2.3506e-7 | -0.9615 | 0.1839 |
| AX-146995791 | Ocu1$_{RPS14}$ | AX-147117324 | 1 | 38741596 | | 5.3507E-04 | 2.0520E-04 | 2.3506e-7 | -0.9615 | 0.1839 |
| AX-146995791 | Ocu1$_{RPS14}$ | AX-147068578 | 1 | 38747021 | | 5.2636E-04 | 2.0193E-04 | 2.2985e-7 | -0.9624 | 0.1839 |
| AX-146995791 | Ocu1$_{RPS14}$ | AX-147028510 | 1 | 38756395 | | 5.2636E-04 | 2.0193E-04 | 2.2985e-7 | -0.9624 | 0.1839 |
| AX-146995791 | Ocu1$_{RPS14}$ | AX-147166054 | 1 | 38769156 | | 4.5580E-04 | 1.7564E-04 | 1.9317e-7 | 0.9656 | 0.1833 |
| AX-146995791 | Ocu1$_{RPS14}$ | AX-147118901 | 1 | 38786430 | | 4.5580E-04 | 1.7564E-04 | 1.9317e-7 | 0.9656 | 0.1833 |
| AX-146995791 | Ocu1$_{RPS14}$ | AX-147148281 | 1 | 38805501 | | 5.3507E-04 | 2.0520E-04 | 2.3506e-7 | -0.9615 | 0.1839 |
| AX-146995791 | Ocu1$_{RPS14}$ | AX-147073490 | 1 | 38828879 | | 5.3507E-04 | 2.0520E-04 | 2.3506e-7 | -0.9615 | 0.1839 |
| AX-146995791 | Ocu1$_{RPS14}$ | AX-147015475 | 1 | 38894456 | | 7.4161E-06 | 3.0949E-06 | 1.0243e-9 | -1.1322 | 0.1825 |
| AX-146995791 | Ocu1$_{RPS14}$ | AX-147000425 | 1 | 38902959 | | 5.0422E-04 | 1.9364E-04 | 2.1723e-7 | -0.9642 | 0.1838 |
| AX-146995791 | Ocu1$_{RPS14}$ | AX-147128518 | 1 | 39199029 | | 2.9093E-04 | 1.1016E-04 | 6.3196e-8 | 1.0810 | 0.1973 |
| AX-146995791 | Ocu1$_{RPS14}$ | AX-147004839 | 1 | 39246389 | | 2.7628E-04 | 1.0472E-04 | 5.8751e-8 | 1.0841 | 0.1974 |
| AX-146995791 | Ocu1$_{RPS14}$ | AX-147053369 | 1 | 39253832 | | 2.3130E-04 | 8.7809E-05 | 4.4015e-8 | 1.1012 | 0.1986 |
| AX-146995791 | Ocu1$_{RPS14}$ | AX-146984698 | 1 | 39267169 | | 2.0490E-04 | 7.8015E-05 | 3.7693e-8 | 1.1056 | 0.1983 |
| AX-146995791 | Ocu1$_{RPS14}$ | AX-147030911 | 1 | 39273878 | | 2.0490E-04 | 7.8015E-05 | 3.7693e-8 | 1.1056 | 0.1983 |
| AX-146995791 | Ocu1$_{RPS14}$ | AX-147015477 | 1 | 39296903 | | 2.1413E-04 | 8.1454E-05 | 3.9999e-8 | 1.1034 | 0.1984 |
| AX-146995791 | Ocu1$_{RPS14}$ | AX-147048247 | 1 | 39310514 | | 2.0490E-04 | 7.8015E-05 | 3.7693e-8 | 1.1056 | 0.1983 |
| AX-146995791 | Ocu1$_{RPS14}$ | AX-147045684 | 1 | 39329536 | | 2.2520E-04 | 8.5566E-05 | 4.2714e-8 | 1.1014 | 0.1984 |
| AX-146995791 | Ocu1$_{RPS14}$ | AX-147045685 | 1 | 39339945 | | 7.8254E-04 | 2.8939E-04 | 2.4000e-7 | 1.0349 | 0.1980 |
| AX-146995791 | Ocu1$_{RPS14}$ | AX-147008079 | 1 | 39362818 | | 2.0490E-04 | 7.8015E-05 | 3.7693e-8 | 1.1056 | 0.1983 |
| AX-146995791 | Ocu1$_{RPS14}$ | AX-147109111 | 1 | 39370513 | | 2.4263E-04 | 9.2075E-05 | 4.7623e-8 | 1.0964 | 0.1982 |
| AX-146995791 | Ocu1$_{RPS14}$ | AX-147110836 | 1 | 39415268 | | 2.6554E-04 | 1.0060E-04 | 5.4150e-8 | 1.0909 | 0.1981 |
| AX-146995791 | Ocu1$_{RPS14}$ | AX-147173500 | 1 | 39425288 | | 2.9232E-04 | 1.1053E-04 | 6.1773e-8 | 1.0859 | 0.1981 |
| AX-146995791 | Ocu1$_{RPS14}$ | AX-147120350 | 1 | 39442193 | | 2.6907E-04 | 1.0192E-04 | 5.5206e-8 | 1.0900 | 0.1981 |
| AX-146995791 | Ocu1$_{RPS14}$ | AX-147004840 | 1 | 39456040 | | 2.6907E-04 | 1.0192E-04 | 5.5206e-8 | 1.0900 | 0.1981 |
| AX-146995791 | Ocu1$_{RPS14}$ | AX-147040634 | 1 | 39464457 | | 2.7196E-04 | 1.0299E-04 | 5.5975e-8 | 1.0896 | 0.1981 |
| AX-146995791 | Ocu1$_{RPS14}$ | AX-147136491 | 1 | 39471467 | | 2.9351E-04 | 1.1096E-04 | 6.2015e-8 | 1.0860 | 0.1981 |
| AX-146995791 | Ocu1$_{RPS14}$ | AX-147132065 | 1 | 39482352 | | 2.9401E-04 | 1.1107E-04 | 6.1235e-8 | -1.0885 | 0.1985 |
| AX-146995791 | Ocu1$_{RPS14}$ | AX-147135048 | 7 | 69856571 | | 3.9609E-04 | 1.5917E-04 | 2.8789e-7 | 0.8614 | 0.1660 |
| AX-146995791 | Ocu1$_{RPS14}$ | AX-147077463 | 7 | 69862659 | | 4.3476E-04 | 1.7433E-04 | 3.2605e-7 | 0.8574 | 0.1660 |
| AX-146995791 | Ocu1$_{RPS14}$ | AX-147101097 | 7 | 69868549 | | 4.3476E-04 | 1.7433E-04 | 3.2605e-7 | 0.8574 | 0.1660 |
| AX-146995791 | Ocu1$_{RPS14}$ | AX-147111946 | 7 | 69885359 | | 4.3476E-04 | 1.7433E-04 | 3.2605e-7 | 0.8574 | 0.1660 |

| | | | | | | | | | |
|---|---|---|---|---|---|---|---|---|---|
| AX-146995791 | Ocu1$_{RPS14}$ | AX-147091052 | 7 | 69893238 | 4.3476E-04 | 1.7433E-04 | 3.2605e-7 | 0.8574 | 0.1660 |
| AX-146995791 | Ocu1$_{RPS14}$ | AX-147057699 | 7 | 70042930 | 4.1148E-04 | 1.6517E-04 | 3.0195e-7 | 0.8604 | 0.1661 |
| AX-146995791 | Ocu1$_{RPS14}$ | AX-147039879 | 7 | 70066319 | 4.1765E-04 | 1.6757E-04 | 3.0740e-7 | 0.8601 | 0.1661 |
| AX-146995791 | Ocu1$_{RPS14}$ | AX-146993531 | 7 | 85582174 | 6.7354E-04 | 2.8816E-04 | 2.1876e-6 | 0.6476 | 0.1354 |
| AX-146995791 | Ocu1$_{RPS14}$ | AX-147170179 | 7 | 159620836 | 3.5683E-04 | 1.4515E-04 | 2.8767e-7 | 0.8406 | 0.1619 |
| AX-146995791 | Ocu1$_{RPS14}$ | AX-147181728 | 9 | 98983387 | 7.1852E-04 | 2.7565E-04 | 3.9563e-7 | -0.9247 | 0.1803 |
| AX-146995791 | Ocu1$_{RPS14}$ | AX-147022140 | 13 | 121883005 | 6.8542E-04 | 3.0778E-04 | 5.1342e-6 | 0.5453 | 0.1185 |
| AX-146995791 | Ocu1$_{RPS14}$ | AX-147012461 | 19 | 35428557 | 4.7183E-04 | 2.0406E-04 | 1.3291e-6 | -0.6604 | 0.1352 |
| AX-146995791 | Ocu1$_{RPS14}$ | AX-147079530 | 20 | 32406003 | 9.8077E-04 | 3.5952E-04 | 3.0798e-7 | 1.0351 | 0.1999 |
| AX-146995791 | Ocu1$_{RPS14}$ | AX-147147933 | 20 | 32411599 | 9.9411E-04 | 3.6427E-04 | 3.1369e-7 | 1.0345 | 0.1999 |
| AX-146995791 | Ocu1$_{RPS14}$ | AX-147106939 | GL018699 | 968 | 5.1604E-04 | 1.8968E-04 | 9.5646e-8 | 1.1192 | 0.2071 |
| AX-146995791 | Ocu1$_{RPS14}$ | AX-147120113 | GL018712 | 8247 | 4.6850E-04 | 2.1027E-04 | 2.5726e-6 | -0.5802 | 0.1222 |
| AX-146995791 | Ocu1$_{RPS14}$ | AX-146995650 | GL018712 | 8248 | 6.8547E-04 | 3.0307E-04 | 4.0571e-6 | -0.5747 | 0.1235 |
| AX-146995791 | Ocu1$_{RPS14}$ | AX-147087022 | GL018712 | 8252 | 1.3040E-04 | 5.7664E-05 | 1.8214e-7 | -0.7345 | 0.1391 |
| AX-146995791 | Ocu1$_{RPS14}$ | AX-147125709 | GL018712 | 8256 | 2.1151E-04 | 9.6317E-05 | 7.6957e-7 | -0.6226 | 0.1245 |
| AX-146995791 | Ocu1$_{RPS14}$ | AX-147037779 | GL018712 | 8308 | 6.3705E-04 | 2.8236E-04 | 3.6899e-6 | -0.5766 | 0.1234 |
| AX-146995791 | Ocu1$_{RPS14}$ | AX-147075521 | GL018712 | 8348 | 7.1481E-04 | 3.1584E-04 | 4.3336e-6 | -0.5720 | 0.1233 |
| AX-146995791 | Ocu1$_{RPS14}$ | AX-147125712 | GL018712 | 8367 | 4.9169E-04 | 2.1052E-04 | 1.1990e-6 | -0.6799 | 0.1385 |
| AX-146995791 | Ocu1$_{RPS14}$ | AX-147150173 | GL018715 | 9873 | 9.3312E-04 | 3.6639E-04 | 9.1009e-7 | 0.8270 | 0.1667 |
| AX-146995791 | Ocu1$_{RPS14}$ | AX-147136324 | GL018715 | 9875 | 8.7930E-04 | 3.4560E-04 | 8.2997e-7 | 0.8315 | 0.1669 |
| AX-146995791 | Ocu1$_{RPS14}$ | AX-147058136 | GL018715 | 9877 | 8.4809E-04 | 3.3385E-04 | 7.9566e-7 | 0.8316 | 0.1667 |
| AX-146995791 | Ocu1$_{RPS14}$ | AX-147128301 | GL018715 | 9878 | 6.6226E-04 | 2.6279E-04 | 5.7833e-7 | 0.8388 | 0.1660 |
| AX-146995791 | Ocu1$_{RPS14}$ | AX-147163141 | GL018749 | 17835 | 9.4265E-04 | 3.3507E-04 | 1.3603e-7 | 1.1714 | 0.2196 |

**Table S7: Significant epistatic interaction with the best SNP marker of the Ocu1$_{TYR}$ locus**

Only individuals carrying coloration at extremities, from cream/beige (P3) to dark chocolate/brown (P6) were considered and analysed using an adaptive shrinkage method. lfsr: local false sign rate, lfdr: local false discovery rate, beta: interaction effect and se: standard error

| SNP1 | Location | SNP2 | Ocu | Position | Gene | lfsr | lfdr | pval | beta | se |
|---|---|---|---|---|---|---|---|---|---|---|
| AX-147073566 | Ocu1$_{TYR}$ | AX-147092879 | 20 | 31689099 | | 2.0952E-04 | 1.0904E-04 | 3.7385e-6 | -0.4136 | 0.0886 |
| AX-147073566 | Ocu1$_{TYR}$ | AX-147001181 | 20 | 31903383 | | 6.5540E-04 | 3.3108E-04 | 1.3291e-5 | -0.3917 | 0.0892 |
| AX-147073566 | Ocu1$_{TYR}$ | AX-147064939 | 20 | 31910012 | | 6.5540E-04 | 3.3108E-04 | 1.3291e-5 | -0.3917 | 0.0892 |
| AX-147073566 | Ocu1$_{TYR}$ | AX-147079529 | 20 | 31919490 | | 6.5540E-04 | 3.3108E-04 | 1.3291e-5 | -0.3917 | 0.0892 |
| AX-147073566 | Ocu1$_{TYR}$ | AX-147067461 | 20 | 31939727 | | 6.5540E-04 | 3.3108E-04 | 1.3291e-5 | -0.3917 | 0.0892 |
| AX-147073566 | Ocu1$_{TYR}$ | AX-147052196 | 20 | 31944944 | | 6.5540E-04 | 3.3108E-04 | 1.3291e-5 | -0.3917 | 0.0892 |
| AX-147073566 | Ocu1$_{TYR}$ | AX-147064940 | 20 | 31960219 | | 6.5540E-04 | 3.3108E-04 | 1.3291e-5 | -0.3917 | 0.0892 |
| AX-147073566 | Ocu1$_{TYR}$ | AX-147027396 | 20 | 31967461 | | 6.5540E-04 | 3.3108E-04 | 1.3291e-5 | -0.3917 | 0.0892 |
| AX-147073566 | Ocu1$_{TYR}$ | AX-146999850 | 20 | 31981614 | | 6.5540E-04 | 3.3108E-04 | 1.3291e-5 | -0.3917 | 0.0892 |
| AX-147073566 | Ocu1$_{TYR}$ | AX-147114958 | 20 | 32076869 | | 6.5540E-04 | 3.3108E-04 | 1.3291e-5 | -0.3917 | 0.0892 |
| AX-147073566 | Ocu1$_{TYR}$ | AX-147172818 | 20 | 32087428 | | 2.4997E-04 | 1.2921E-04 | 4.4890e-6 | -0.4129 | 0.0892 |
| AX-147073566 | Ocu1$_{TYR}$ | AX-147164888 | 20 | 32092947 | | 7.2196E-04 | 3.6347E-04 | 1.4764e-5 | -0.3904 | 0.0894 |
| AX-147073566 | Ocu1$_{TYR}$ | AX-147178794 | 20 | 33187157 | | 8.6337E-04 | 4.3568E-04 | 1.8574e-5 | -0.3796 | 0.0879 |

**Table S8: Significant epistatic interaction with the best SNP marker of the Ocu4$_{ASIP}$ locus**

Only individuals carrying coloration at extremities, from cream/beige (P3) to dark chocolate/brown (P6) were considered and analysed using an adaptive shrinkage method. lfsr: local false sign rate, lfdr: local false discovery rate, beta: interaction effect and se: standard error

| SNP1 | Location | SNP2 | Ocu | Position | Gene | lfsr | lfdr | pval | beta | se |
|---|---|---|---|---|---|---|---|---|---|---|
| AX-147097074 | Ocu4$_{ASIP}$ | AX-147056054 | 1 | 157632880 | | 1.9152E-04 | 8.8328E-05 | 8.0176e-7 | -0.6042 | 0.1211 |
| AX-147097074 | Ocu4$_{ASIP}$ | AX-147095962 | 1 | 157845651 | | 1.3025E-04 | 6.0734E-05 | 4.8595e-7 | -0.6141 | 0.1207 |
| AX-147097074 | Ocu4$_{ASIP}$ | AX-147058560 | 1 | 157896081 | | 1.4499E-04 | 6.7405E-05 | 5.5859e-7 | -0.6113 | 0.1208 |
| AX-147097074 | Ocu4$_{ASIP}$ | AX-147045792 | 1 | 157921078 | | 2.6158E-04 | 1.1992E-04 | 1.2751e-6 | -0.5893 | 0.1204 |
| AX-147097074 | Ocu4$_{ASIP}$ | AX-147028641 | 1 | 158024874 | | 1.4499E-04 | 6.7405E-05 | 5.5859e-7 | -0.6113 | 0.1208 |
| AX-147097074 | Ocu4$_{ASIP}$ | AX-147080695 | 1 | 158038939 | | 2.4868E-04 | 1.1633E-04 | 1.6310e-6 | -0.5544 | 0.1145 |
| AX-147097074 | Ocu4$_{ASIP}$ | AX-147101842 | 1 | 158046548 | | 2.5667E-04 | 1.1996E-04 | 1.6983e-6 | -0.5536 | 0.1145 |
| AX-147097074 | Ocu4$_{ASIP}$ | AX-146992129 | 1 | 158056875 | | 4.4411E-04 | 2.0468E-04 | 3.5182e-6 | -0.5352 | 0.1143 |
| AX-147097074 | Ocu4$_{ASIP}$ | AX-147143006 | 1 | 158062459 | | 4.2584E-04 | 1.9649E-04 | 3.3308e-6 | -0.5365 | 0.1143 |
| AX-147097074 | Ocu4$_{ASIP}$ | AX-147161273 | 1 | 158132219 | | 4.4411E-04 | 2.0468E-04 | 3.5182e-6 | -0.5352 | 0.1143 |
| AX-147097074 | Ocu4$_{ASIP}$ | AX-147101843 | 1 | 158149694 | | 4.4411E-04 | 2.0468E-04 | 3.5182e-6 | -0.5352 | 0.1143 |
| AX-147097074 | Ocu4$_{ASIP}$ | AX-147050932 | 1 | 158157039 | | 4.4411E-04 | 2.0468E-04 | 3.5182e-6 | -0.5352 | 0.1143 |
| AX-147097074 | Ocu4$_{ASIP}$ | AX-147109190 | 1 | 158164032 | | 4.4411E-04 | 2.0468E-04 | 3.5182e-6 | -0.5352 | 0.1143 |
| AX-147097074 | Ocu4$_{ASIP}$ | AX-147170548 | 2 | 20102246 | | 5.2929E-04 | 2.4403E-04 | 4.7378e-6 | 0.5209 | 0.1128 |
| AX-147097074 | Ocu4$_{ASIP}$ | AX-147041805 | 2 | 20124589 | | 7.5090E-04 | 3.4938E-04 | 9.1428e-6 | 0.4811 | 0.1075 |
| AX-147097074 | Ocu4$_{ASIP}$ | AX-147057142 | 2 | 20158990 | | 4.1020E-04 | 1.9423E-04 | 4.3470e-6 | 0.4973 | 0.1072 |
| AX-147097074 | Ocu4$_{ASIP}$ | AX-147022691 | 2 | 20167714 | | 2.7423E-04 | 1.3165E-04 | 2.7359e-6 | 0.5034 | 0.1063 |
| AX-147097074 | Ocu4$_{ASIP}$ | AX-147022692 | 2 | 20233498 | | 4.1020E-04 | 1.9423E-04 | 4.3470e-6 | 0.4973 | 0.1072 |
| AX-147097074 | Ocu4$_{ASIP}$ | AX-147138906 | 2 | 20267336 | | 4.2972E-04 | 2.0321E-04 | 4.6050e-6 | 0.4960 | 0.1072 |
| AX-147097074 | Ocu4$_{ASIP}$ | AX-147132546 | 2 | 20495263 | | 6.2857E-04 | 2.9365E-04 | 7.2585e-6 | 0.4877 | 0.1077 |
| AX-147097074 | Ocu4$_{ASIP}$ | AX-147039297 | 2 | 20508309 | | 4.0027E-04 | 1.8963E-04 | 4.2120e-6 | 0.4981 | 0.1072 |
| AX-147097074 | Ocu4$_{ASIP}$ | AX-146995218 | 2 | 20514362 | | 4.1927E-04 | 1.9838E-04 | 4.4615e-6 | 0.4968 | 0.1073 |
| AX-147097074 | Ocu4$_{ASIP}$ | AX-147079342 | 2 | 20530150 | | 4.0027E-04 | 1.8963E-04 | 4.2120e-6 | 0.4981 | 0.1072 |
| AX-147097074 | Ocu4$_{ASIP}$ | AX-147100729 | 2 | 20543793 | | 5.4689E-04 | 2.5674E-04 | 6.1673e-6 | 0.4902 | 0.1075 |
| AX-147097074 | Ocu4$_{ASIP}$ | AX-147034407 | 2 | 20550013 | | 4.3706E-04 | 2.0654E-04 | 4.6914e-6 | 0.4958 | 0.1073 |
| AX-147097074 | Ocu4$_{ASIP}$ | AX-147012493 | 2 | 20569931 | | 4.0027E-04 | 1.8963E-04 | 4.2120e-6 | 0.4981 | 0.1072 |
| AX-147097074 | Ocu4$_{ASIP}$ | AX-147104476 | 2 | 20592178 | | 3.7557E-04 | 1.7828E-04 | 3.9015e-6 | 0.4995 | 0.1072 |
| AX-147097074 | Ocu4$_{ASIP}$ | AX-146995219 | 2 | 20603573 | | 2.8933E-04 | 1.3760E-04 | 2.6488e-6 | 0.5147 | 0.1085 |
| AX-147097074 | Ocu4$_{ASIP}$ | AX-147131387 | 2 | 26801380 | | 8.8599E-04 | 4.1673E-04 | 1.2870e-5 | 0.4554 | 0.1035 |
| AX-147097074 | Ocu4$_{ASIP}$ | AX-147139855 | 2 | 29195251 | | 7.3358E-04 | 3.3145E-04 | 6.2860e-6 | 0.5287 | 0.1160 |
| AX-147097074 | Ocu4$_{ASIP}$ | AX-147092732 | 2 | 29481817 | | 9.5272E-04 | 4.2694E-04 | 8.7978e-6 | 0.5209 | 0.1162 |
| AX-147097074 | Ocu4$_{ASIP}$ | AX-146994884 | 12 | 82013581 | | 4.8085E-04 | 2.1936E-04 | 3.4808e-6 | 0.5469 | 0.1167 |
| AX-147097074 | Ocu4$_{ASIP}$ | AX-147146270 | 13 | 64510533 | | 9.4697E-04 | 4.3455E-04 | 1.1377e-5 | -0.4845 | 0.1094 |
| AX-147097074 | Ocu4$_{ASIP}$ | AX-147043648 | 13 | 64526616 | | 9.4697E-04 | 4.3455E-04 | 1.1377e-5 | -0.4845 | 0.1094 |
| AX-147097074 | Ocu4$_{ASIP}$ | AX-147083389 | 13 | 64544712 | | 9.4697E-04 | 4.3455E-04 | 1.1377e-5 | -0.4845 | 0.1094 |
| AX-147097074 | Ocu4$_{ASIP}$ | AX-147022091 | 13 | 64557339 | | 9.0524E-04 | 4.1594E-04 | 1.0748e-5 | -0.4859 | 0.1094 |
| AX-147097074 | Ocu4$_{ASIP}$ | AX-146984271 | 13 | 64565476 | | 9.4697E-04 | 4.3455E-04 | 1.1377e-5 | -0.4845 | 0.1094 |
| AX-147097074 | Ocu4$_{ASIP}$ | AX-147169990 | 13 | 65330868 | | 4.6055E-04 | 2.1653E-04 | 4.8035e-6 | -0.5000 | 0.1083 |
| AX-147097074 | Ocu4$_{ASIP}$ | AX-147026600 | 13 | 65582366 | | 4.6055E-04 | 2.1653E-04 | 4.8035e-6 | -0.5000 | 0.1083 |
| AX-147097074 | Ocu4$_{ASIP}$ | AX-147015884 | 13 | 65594870 | | 4.6055E-04 | 2.1653E-04 | 4.8035e-6 | -0.5000 | 0.1083 |
| AX-147097074 | Ocu4$_{ASIP}$ | AX-146984724 | 13 | 65609960 | | 8.0611E-04 | 3.5690E-04 | 5.5651e-6 | -0.5569 | 0.1215 |
| AX-147097074 | Ocu4$_{ASIP}$ | AX-147000683 | 13 | 65621337 | | 8.7664E-05 | 4.3011E-05 | 5.9871e-7 | -0.5444 | 0.1079 |
| AX-147097074 | Ocu4$_{ASIP}$ | AX-147006798 | 13 | 65652188 | | 1.0851E-04 | 5.2992E-05 | 7.8923e-7 | -0.5381 | 0.1078 |
| AX-147097074 | Ocu4$_{ASIP}$ | AX-147017915 | 13 | 65667044 | | 1.0851E-04 | 5.2992E-05 | 7.8923e-7 | -0.5381 | 0.1078 |
| AX-147097074 | Ocu4$_{ASIP}$ | AX-146987492 | 13 | 65672135 | | 1.0851E-04 | 5.2992E-05 | 7.8923e-7 | -0.5381 | 0.1078 |
| AX-147097074 | Ocu4$_{ASIP}$ | AX-147081041 | 13 | 65677357 | | 1.0851E-04 | 5.2992E-05 | 7.8923e-7 | -0.5381 | 0.1078 |
| AX-147097074 | Ocu4$_{ASIP}$ | AX-147022092 | 13 | 65690098 | | 1.0392E-04 | 5.0795E-05 | 7.4627e-7 | -0.5394 | 0.1078 |
| AX-147097074 | Ocu4$_{ASIP}$ | AX-147022093 | 13 | 65697786 | | 3.9997E-05 | 1.9941E-05 | 2.1335e-7 | -0.5682 | 0.1082 |
| AX-147097074 | Ocu4$_{ASIP}$ | AX-147175689 | 13 | 65722107 | | 3.1394E-04 | 1.4944E-04 | 3.0486e-6 | -0.5075 | 0.1077 |

| | | | | | | | | | |
|---|---|---|---|---|---|---|---|---|---|
| AX-147097074 | Ocu4$_{ASIP}$ | AX-147036196 | 13 | 65727651 | 3.2657E-04 | 1.5532E-04 | 3.2112e-6 | -0.5061 | 0.1076 |
| AX-147097074 | Ocu4$_{ASIP}$ | AX-147058967 | 13 | 65734390 | 1.4979E-04 | 7.2236E-05 | 1.1073e-6 | -0.5376 | 0.1092 |
| AX-147097074 | Ocu4$_{ASIP}$ | AX-146988401 | 13 | 65740088 | 1.5848E-04 | 7.6308E-05 | 1.1866e-6 | -0.5364 | 0.1093 |
| AX-147097074 | Ocu4$_{ASIP}$ | AX-146998216 | 13 | 65750335 | 2.9236E-04 | 1.3945E-04 | 2.7931e-6 | -0.5093 | 0.1076 |
| AX-147097074 | Ocu4$_{ASIP}$ | AX-147102121 | 13 | 65771540 | 2.9236E-04 | 1.3945E-04 | 2.7931e-6 | -0.5093 | 0.1076 |
| AX-147097074 | Ocu4$_{ASIP}$ | AX-147100242 | 13 | 65784623 | 1.2735E-04 | 6.1491E-05 | 8.6349e-7 | -0.5469 | 0.1100 |
| AX-147097074 | Ocu4$_{ASIP}$ | AX-147010173 | 13 | 65808409 | 2.9236E-04 | 1.3945E-04 | 2.7931e-6 | -0.5093 | 0.1076 |
| AX-147097074 | Ocu4$_{ASIP}$ | AX-147096285 | 13 | 65814593 | 2.9236E-04 | 1.3945E-04 | 2.7931e-6 | -0.5093 | 0.1076 |
| AX-147097074 | Ocu4$_{ASIP}$ | AX-146988893 | 13 | 65820680 | 2.9236E-04 | 1.3945E-04 | 2.7931e-6 | -0.5093 | 0.1076 |
| AX-147097074 | Ocu4$_{ASIP}$ | AX-147087769 | 13 | 65833120 | 2.9236E-04 | 1.3945E-04 | 2.7931e-6 | -0.5093 | 0.1076 |
| AX-147097074 | Ocu4$_{ASIP}$ | AX-147022094 | 13 | 65847915 | 3.6765E-04 | 1.7416E-04 | 3.6857e-6 | -0.5041 | 0.1079 |
| AX-147097074 | Ocu4$_{ASIP}$ | AX-147111150 | 13 | 65855798 | 2.9236E-04 | 1.3945E-04 | 2.7931e-6 | -0.5093 | 0.1076 |
| AX-147097074 | Ocu4$_{ASIP}$ | AX-147081042 | 13 | 65867442 | 1.1375E-04 | 5.5539E-05 | 8.4998e-7 | -0.5351 | 0.1075 |
| AX-147097074 | Ocu4$_{ASIP}$ | AX-147041125 | 13 | 65882168 | 2.9236E-04 | 1.3945E-04 | 2.7931e-6 | -0.5093 | 0.1076 |
| AX-147097074 | Ocu4$_{ASIP}$ | AX-147109452 | 13 | 65888854 | 2.9236E-04 | 1.3945E-04 | 2.7931e-6 | -0.5093 | 0.1076 |
| AX-147097074 | Ocu4$_{ASIP}$ | AX-147107653 | 13 | 65895133 | 2.9236E-04 | 1.3945E-04 | 2.7931e-6 | -0.5093 | 0.1076 |
| AX-147097074 | Ocu4$_{ASIP}$ | AX-147061474 | 13 | 65903743 | 9.1294E-04 | 4.2156E-04 | 1.1435e-5 | -0.4782 | 0.1080 |
| AX-147097074 | Ocu4$_{ASIP}$ | AX-147013871 | 13 | 65910218 | 2.9236E-04 | 1.3945E-04 | 2.7931e-6 | -0.5093 | 0.1076 |
| AX-147097074 | Ocu4$_{ASIP}$ | AX-147141448 | 13 | 65915470 | 2.9236E-04 | 1.3945E-04 | 2.7931e-6 | -0.5093 | 0.1076 |
| AX-147097074 | Ocu4$_{ASIP}$ | AX-146987493 | 13 | 65935475 | 2.9236E-04 | 1.3945E-04 | 2.7931e-6 | -0.5093 | 0.1076 |
| AX-147097074 | Ocu4$_{ASIP}$ | AX-147144746 | 13 | 65940555 | 2.9236E-04 | 1.3945E-04 | 2.7931e-6 | -0.5093 | 0.1076 |
| AX-147097074 | Ocu4$_{ASIP}$ | AX-147149758 | 13 | 65951823 | 5.8556E-04 | 2.7450E-04 | 6.7479e-6 | -0.4876 | 0.1073 |
| AX-147097074 | Ocu4$_{ASIP}$ | AX-147136665 | 13 | 66017280 | 2.9236E-04 | 1.3945E-04 | 2.7931e-6 | -0.5093 | 0.1076 |
| AX-147097074 | Ocu4$_{ASIP}$ | AX-146987927 | 13 | 66024131 | 2.9236E-04 | 1.3945E-04 | 2.7931e-6 | -0.5093 | 0.1076 |
| AX-147097074 | Ocu4$_{ASIP}$ | AX-146985111 | 13 | 66032301 | 2.9236E-04 | 1.3945E-04 | 2.7931e-6 | -0.5093 | 0.1076 |
| AX-147097074 | Ocu4$_{ASIP}$ | AX-147019996 | 13 | 66042897 | 2.3682E-04 | 1.1349E-04 | 2.1256e-6 | -0.5162 | 0.1078 |
| AX-147097074 | Ocu4$_{ASIP}$ | AX-147143130 | 13 | 66050552 | 6.5060E-04 | 3.0018E-04 | 6.6830e-6 | -0.5040 | 0.1109 |
| AX-147097074 | Ocu4$_{ASIP}$ | AX-147092162 | 13 | 66063625 | 2.9236E-04 | 1.3945E-04 | 2.7931e-6 | -0.5093 | 0.1076 |
| AX-147097074 | Ocu4$_{ASIP}$ | AX-147024312 | 13 | 66090967 | 3.1767E-04 | 1.5104E-04 | 3.0627e-6 | -0.5085 | 0.1079 |
| AX-147097074 | Ocu4$_{ASIP}$ | AX-147038647 | 13 | 66098683 | 2.9236E-04 | 1.3945E-04 | 2.7931e-6 | -0.5093 | 0.1076 |
| AX-147097074 | Ocu4$_{ASIP}$ | AX-147085589 | 13 | 66112983 | 2.8998E-04 | 1.3835E-04 | 2.7668e-6 | -0.5094 | 0.1076 |
| AX-147097074 | Ocu4$_{ASIP}$ | AX-147085590 | 13 | 66119346 | 3.0867E-04 | 1.4700E-04 | 2.9850e-6 | -0.5079 | 0.1077 |
| AX-147097074 | Ocu4$_{ASIP}$ | AX-147117650 | 13 | 66141511 | 2.8998E-04 | 1.3835E-04 | 2.7668e-6 | -0.5094 | 0.1076 |
| AX-147097074 | Ocu4$_{ASIP}$ | AX-147116084 | 13 | 66148282 | 2.8998E-04 | 1.3835E-04 | 2.7668e-6 | -0.5094 | 0.1076 |
| AX-147097074 | Ocu4$_{ASIP}$ | AX-147145551 | 13 | 66156448 | 2.8998E-04 | 1.3835E-04 | 2.7668e-6 | -0.5094 | 0.1076 |
| AX-147097074 | Ocu4$_{ASIP}$ | AX-147114455 | 13 | 66167707 | 2.8998E-04 | 1.3835E-04 | 2.7668e-6 | -0.5094 | 0.1076 |
| AX-147097074 | Ocu4$_{ASIP}$ | AX-146987101 | 13 | 66181898 | 3.1779E-04 | 1.5118E-04 | 3.0849e-6 | -0.5076 | 0.1078 |
| AX-147097074 | Ocu4$_{ASIP}$ | AX-147131095 | 13 | 66191475 | 2.8998E-04 | 1.3835E-04 | 2.7668e-6 | -0.5094 | 0.1076 |
| AX-147097074 | Ocu4$_{ASIP}$ | AX-147053930 | 13 | 66198252 | 3.1860E-04 | 1.5151E-04 | 3.0837e-6 | -0.5080 | 0.1078 |
| AX-147097074 | Ocu4$_{ASIP}$ | AX-147005180 | 13 | 66207313 | 2.8998E-04 | 1.3835E-04 | 2.7668e-6 | -0.5094 | 0.1076 |
| AX-147097074 | Ocu4$_{ASIP}$ | AX-147073937 | 13 | 66219378 | 2.8998E-04 | 1.3835E-04 | 2.7668e-6 | -0.5094 | 0.1076 |
| AX-147097074 | Ocu4$_{ASIP}$ | AX-147022095 | 13 | 66229746 | 2.8998E-04 | 1.3835E-04 | 2.7668e-6 | -0.5094 | 0.1076 |
| AX-147097074 | Ocu4$_{ASIP}$ | AX-147013872 | 13 | 66246964 | 2.8998E-04 | 1.3835E-04 | 2.7668e-6 | -0.5094 | 0.1076 |
| AX-147097074 | Ocu4$_{ASIP}$ | AX-147031414 | 13 | 66873358 | 1.9652E-04 | 9.5155E-05 | 1.8087e-6 | -0.5122 | 0.1062 |
| AX-147097074 | Ocu4$_{ASIP}$ | AX-147015886 | 13 | 66884993 | 1.9652E-04 | 9.5155E-05 | 1.8087e-6 | -0.5122 | 0.1062 |
| AX-147097074 | Ocu4$_{ASIP}$ | AX-146988402 | 13 | 66899231 | 1.9652E-04 | 9.5155E-05 | 1.8087e-6 | -0.5122 | 0.1062 |
| AX-147097074 | Ocu4$_{ASIP}$ | AX-147002143 | 13 | 66905606 | 1.9652E-04 | 9.5155E-05 | 1.8087e-6 | -0.5122 | 0.1062 |
| AX-147097074 | Ocu4$_{ASIP}$ | AX-147090023 | 13 | 66919393 | 1.9652E-04 | 9.5155E-05 | 1.8087e-6 | -0.5122 | 0.1062 |
| AX-147097074 | Ocu4$_{ASIP}$ | AX-147133409 | 13 | 66947852 | 1.9652E-04 | 9.5155E-05 | 1.8087e-6 | -0.5122 | 0.1062 |
| AX-147097074 | Ocu4$_{ASIP}$ | AX-147096287 | 13 | 67033092 | 1.8618E-04 | 9.0260E-05 | 1.6889e-6 | -0.5138 | 0.1062 |
| AX-147097074 | Ocu4$_{ASIP}$ | AX-147019998 | 13 | 67042258 | 1.7155E-04 | 8.3301E-05 | 1.5161e-6 | -0.5167 | 0.1063 |
| AX-147097074 | Ocu4$_{ASIP}$ | AX-147109453 | 13 | 67059620 | 1.8618E-04 | 9.0260E-05 | 1.6889e-6 | -0.5138 | 0.1062 |
| AX-147097074 | Ocu4$_{ASIP}$ | AX-147096288 | 13 | 67081983 | 1.8618E-04 | 9.0260E-05 | 1.6889e-6 | -0.5138 | 0.1062 |
| AX-147097074 | Ocu4$_{ASIP}$ | AX-147012005 | 13 | 67089536 | 1.8618E-04 | 9.0260E-05 | 1.6889e-6 | -0.5138 | 0.1062 |
| AX-147097074 | Ocu4$_{ASIP}$ | AX-147036197 | 13 | 67096254 | 7.8828E-05 | 3.8965E-05 | 5.6569e-7 | -0.5383 | 0.1064 |

| | | | | | | | | | | |
|---|---|---|---|---|---|---|---|---|---|---|
| AX-147097074 | Ocu4$_{ASIP}$ | AX-147094246 | 13 | 67102289 | | 1.8618E-04 | 9.0260E-05 | 1.6889e-6 | -0.5138 | 0.1062 |
| AX-147097074 | Ocu4$_{ASIP}$ | AX-147029047 | 13 | 67114445 | | 1.8618E-04 | 9.0260E-05 | 1.6889e-6 | -0.5138 | 0.1062 |
| AX-147097074 | Ocu4$_{ASIP}$ | AX-147038649 | 13 | 67122247 | | 1.8618E-04 | 9.0260E-05 | 1.6889e-6 | -0.5138 | 0.1062 |
| AX-147097074 | Ocu4$_{ASIP}$ | AX-146993123 | 13 | 67135131 | | 1.8618E-04 | 9.0260E-05 | 1.6889e-6 | -0.5138 | 0.1062 |
| AX-147097074 | Ocu4$_{ASIP}$ | AX-147073940 | 13 | 67150197 | | 2.8271E-04 | 1.3558E-04 | 2.8315e-6 | -0.5030 | 0.1064 |
| AX-147097074 | Ocu4$_{ASIP}$ | AX-146999454 | 13 | 67163138 | | 1.8618E-04 | 9.0260E-05 | 1.6889e-6 | -0.5138 | 0.1062 |
| AX-147097074 | Ocu4$_{ASIP}$ | AX-147076338 | 13 | 67169427 | | 1.8618E-04 | 9.0260E-05 | 1.6889e-6 | -0.5138 | 0.1062 |
| AX-147097074 | Ocu4$_{ASIP}$ | AX-147053931 | 13 | 67177147 | | 1.4691E-04 | 7.1623E-05 | 1.2531e-6 | -0.5204 | 0.1063 |
| AX-147097074 | Ocu4$_{ASIP}$ | AX-147128760 | 13 | 67194024 | | 1.8618E-04 | 9.0260E-05 | 1.6889e-6 | -0.5138 | 0.1062 |
| AX-147097074 | Ocu4$_{ASIP}$ | AX-147111151 | 13 | 67208025 | | 3.5588E-04 | 1.7051E-04 | 3.9837e-6 | -0.4892 | 0.1051 |
| AX-147097074 | Ocu4$_{ASIP}$ | AX-147019999 | 13 | 67213863 | | 1.8618E-04 | 9.0260E-05 | 1.6889e-6 | -0.5138 | 0.1062 |
| AX-147097074 | Ocu4$_{ASIP}$ | AX-147143133 | 13 | 68385779 | | 4.8065E-04 | 2.3040E-04 | 6.2506e-6 | -0.4692 | 0.1029 |
| AX-147097074 | Ocu4$_{ASIP}$ | AX-147010174 | 13 | 68401557 | | 3.9794E-04 | 1.9192E-04 | 5.0267e-6 | -0.4726 | 0.1026 |
| AX-147097074 | Ocu4$_{ASIP}$ | AX-147136666 | 13 | 68421057 | | 6.5172E-04 | 3.1001E-04 | 9.0767e-6 | -0.4603 | 0.1028 |
| AX-147097074 | Ocu4$_{ASIP}$ | AX-147159333 | 13 | 68443494 | | 3.9794E-04 | 1.9192E-04 | 5.0267e-6 | -0.4726 | 0.1026 |
| AX-147097074 | Ocu4$_{ASIP}$ | AX-147046241 | 13 | 68493862 | | 3.9794E-04 | 1.9192E-04 | 5.0267e-6 | -0.4726 | 0.1026 |
| AX-147097074 | Ocu4$_{ASIP}$ | AX-147006799 | 13 | 68515774 | | 3.7088E-04 | 1.7916E-04 | 4.6058e-6 | -0.4748 | 0.1026 |
| AX-147097074 | Ocu4$_{ASIP}$ | AX-147002145 | 13 | 68581460 | | 4.1646E-04 | 2.0051E-04 | 5.2855e-6 | -0.4721 | 0.1027 |
| AX-147097074 | Ocu4$_{ASIP}$ | AX-147000686 | 13 | 68745544 | | 3.8473E-04 | 1.8575E-04 | 4.8330e-6 | -0.4733 | 0.1026 |
| AX-147097074 | Ocu4$_{ASIP}$ | AX-147096301 | 13 | 82097700 | | 3.3029E-04 | 1.6341E-04 | 4.8219e-6 | 0.4500 | 0.0975 |
| AX-147097074 | Ocu4$_{ASIP}$ | AX-147076352 | 13 | 82108305 | | 1.7237E-04 | 8.6358E-05 | 2.1766e-6 | -0.4698 | 0.0982 |
| AX-147097074 | Ocu4$_{ASIP}$ | AX-146994954 | 13 | 82116856 | | 1.8492E-04 | 9.1952E-05 | 2.2341e-6 | -0.4762 | 0.0997 |
| AX-147097074 | Ocu4$_{ASIP}$ | AX-147031434 | 13 | 82122481 | | 1.7237E-04 | 8.6358E-05 | 2.1766e-6 | -0.4698 | 0.0982 |
| AX-147097074 | Ocu4$_{ASIP}$ | AX-147071576 | 13 | 82151617 | | 1.9255E-04 | 9.6181E-05 | 2.4732e-6 | -0.4676 | 0.0983 |
| AX-147097074 | Ocu4$_{ASIP}$ | AX-147109462 | 13 | 82184551 | | 1.7237E-04 | 8.6358E-05 | 2.1766e-6 | -0.4698 | 0.0982 |
| AX-147097074 | Ocu4$_{ASIP}$ | AX-147064099 | 13 | 82253671 | | 1.7237E-04 | 8.6358E-05 | 2.1766e-6 | -0.4698 | 0.0982 |
| AX-147097074 | Ocu4$_{ASIP}$ | AX-147015903 | 13 | 82263720 | | 1.7237E-04 | 8.6358E-05 | 2.1766e-6 | -0.4698 | 0.0982 |
| AX-147097074 | Ocu4$_{ASIP}$ | AX-147139643 | 13 | 85343198 | | 1.1111E-04 | 5.5666E-05 | 1.1533e-6 | 0.4950 | 0.1007 |
| AX-147097074 | Ocu4$_{ASIP}$ | AX-147083413 | 13 | 85362023 | | 3.5668E-04 | 1.7182E-04 | 4.2304e-6 | 0.4812 | 0.1036 |
| AX-147097074 | Ocu4$_{ASIP}$ | AX-147087792 | 13 | 85370010 | | 1.1111E-04 | 5.5666E-05 | 1.1533e-6 | 0.4950 | 0.1007 |
| AX-147097074 | Ocu4$_{ASIP}$ | AX-146998232 | 13 | 85381053 | | 1.1086E-04 | 5.5432E-05 | 1.1237e-6 | 0.4981 | 0.1012 |
| AX-147097074 | Ocu4$_{ASIP}$ | AX-147041147 | 13 | 85388370 | | 1.1111E-04 | 5.5666E-05 | 1.1533e-6 | 0.4950 | 0.1007 |
| AX-147097074 | Ocu4$_{ASIP}$ | AX-147085615 | 13 | 87951250 | | 6.1283E-04 | 2.9322E-04 | 8.7277e-6 | 0.4565 | 0.1018 |
| AX-147097074 | Ocu4$_{ASIP}$ | AX-147064105 | 13 | 87959662 | | 6.4361E-04 | 3.0861E-04 | 9.5161e-6 | 0.4508 | 0.1009 |
| AX-147097074 | Ocu4$_{ASIP}$ | AX-147073962 | 13 | 88115864 | | 6.1283E-04 | 2.9322E-04 | 8.7277e-6 | 0.4565 | 0.1018 |
| AX-147097074 | Ocu4$_{ASIP}$ | AX-147165670 | 13 | 88568374 | | 3.5211E-04 | 1.7160E-04 | 4.6384e-6 | 0.4660 | 0.1008 |
| AX-147097074 | Ocu4$_{ASIP}$ | AX-147022110 | 13 | 88617113 | | 5.5434E-04 | 2.6517E-04 | 7.5437e-6 | 0.4631 | 0.1025 |
| AX-147097074 | Ocu4$_{ASIP}$ | AX-147033831 | 13 | 88622302 | | 6.1283E-04 | 2.9322E-04 | 8.7277e-6 | 0.4565 | 0.1018 |
| AX-147097074 | Ocu4$_{ASIP}$ | AX-147061500 | 13 | 88629643 | | 7.4355E-04 | 3.5520E-04 | 1.1325e-5 | 0.4467 | 0.1009 |
| AX-147097074 | Ocu4$_{ASIP}$ | AX-147085617 | 13 | 89166403 | | 5.4755E-04 | 2.6197E-04 | 7.4219e-6 | 0.4636 | 0.1025 |
| AX-147097074 | Ocu4$_{ASIP}$ | AX-147053969 | 13 | 89176260 | | 5.4755E-04 | 2.6197E-04 | 7.4219e-6 | 0.4636 | 0.1025 |
| AX-147097074 | Ocu4$_{ASIP}$ | AX-147046262 | 13 | 89182555 | | 5.4755E-04 | 2.6197E-04 | 7.4219e-6 | 0.4636 | 0.1025 |
| AX-147097074 | Ocu4$_{ASIP}$ | AX-147041437 | 15 | 55262235 | | 3.7831E-04 | 1.9728E-04 | 7.7768e-6 | 0.3853 | 0.0854 |
| AX-147097074 | Ocu4$_{ASIP}$ | AX-147069634 | 18 | 33325902 | | 8.2746E-04 | 4.0301E-04 | 1.4887e-5 | -0.4179 | 0.0957 |
| AX-147097074 | Ocu4$_{ASIP}$ | AX-147070592 | GL018699 | 116 | | 6.4941E-04 | 2.7398E-04 | 1.5644e-6 | 0.6849 | 0.1412 |
| AX-147097074 | Ocu4$_{ASIP}$ | AX-147125765 | GL018733 | 14917 | | 6.9982E-04 | 3.1067E-04 | 4.4904e-6 | -0.5649 | 0.1220 |
| AX-147097074 | Ocu4$_{ASIP}$ | AX-147045403 | GL018733 | 14919 | | 6.9754E-04 | 3.0986E-04 | 4.5080e-6 | -0.5639 | 0.1218 |
| AX-147097074 | Ocu4$_{ASIP}$ | AX-147063247 | GL018733 | 14920 | | 8.6753E-04 | 3.8160E-04 | 5.7708e-6 | -0.5625 | 0.1229 |
| AX-147097074 | Ocu4$_{ASIP}$ | AX-147152079 | GL018733 | 14922 | | 3.5569E-04 | 1.6140E-04 | 1.8705e-6 | -0.5831 | 0.1211 |
| AX-147097074 | Ocu4$_{ASIP}$ | AX-147172515 | GL018733 | 14925 | | 6.3892E-04 | 2.8456E-04 | 4.0037e-6 | -0.5669 | 0.1218 |
| AX-147097074 | Ocu4$_{ASIP}$ | AX-147117134 | GL018733 | 14926 | | 7.0744E-04 | 3.1415E-04 | 4.5998e-6 | -0.5633 | 0.1218 |
| AX-147097074 | Ocu4$_{ASIP}$ | AX-147011438 | GL018733 | 14949 | *HPS5* | 1.2448E-04 | 5.8173E-05 | 4.6718e-7 | -0.6132 | 0.1203 |
| AX-147097074 | Ocu4$_{ASIP}$ | AX-146999046 | GL018733 | 14963 | *HPS5* | 4.2851E-04 | 1.9726E-04 | 3.2652e-6 | -0.5402 | 0.1149 |
| AX-147097074 | Ocu4$_{ASIP}$ | AX-147099642 | GL018733 | 14965 | *HPS5* | 2.1633E-04 | 9.8775E-05 | 8.3410e-7 | -0.6147 | 0.1234 |
| AX-147097074 | Ocu4$_{ASIP}$ | AX-147068298 | GL018733 | 14979 | | 1.9152E-04 | 8.8328E-05 | 8.0176e-7 | -0.6042 | 0.1211 |

| | | | | | | | | | | |
|---|---|---|---|---|---|---|---|---|---|---|
| AX-147097074 | Ocu4$_{ASIP}$ | AX-147107074 | GL018733 | 14987 | | 2.0812E-04 | 9.5899E-05 | 9.1887e-7 | -0.5988 | 0.1207 |
| AX-147097074 | Ocu4$_{ASIP}$ | AX-147173844 | GL018733 | 14990 | | 4.1177E-04 | 1.8750E-04 | 2.5793e-6 | -0.5634 | 0.1187 |
| AX-147097074 | Ocu4$_{ASIP}$ | AX-147171265 | GL018941 | 27698 | | 9.5660E-05 | 4.1058E-05 | 7.3084e-8 | -0.8307 | 0.1523 |
| AX-147097074 | Ocu4$_{ASIP}$ | MC1R | GL018965 | 14 | *MC1R* | 4.1948E-04 | 1.7351E-04 | 4.8883e-7 | -0.7817 | 0.1536 |
| AX-147097074 | Ocu4$_{ASIP}$ | AX-147171269 | GL018965 | 28258 | | 8.9187E-05 | 3.8316E-05 | 6.6372e-8 | -0.8343 | 0.1525 |
| AX-147097074 | Ocu4$_{ASIP}$ | AX-147176467 | GL018965 | 28260 | | 1.0255E-04 | 4.3834E-05 | 7.7193e-8 | -0.8347 | 0.1534 |
| AX-147097074 | Ocu4$_{ASIP}$ | AX-147171270 | GL018965 | 28265 | | 9.7162E-05 | 4.1610E-05 | 7.2487e-8 | -0.8346 | 0.1530 |
| AX-147097074 | Ocu4$_{ASIP}$ | AX-147144567 | GL018965 | 28272 | | 1.6276E-04 | 6.7422E-05 | 1.0482e-7 | -0.8706 | 0.1617 |
| AX-147097074 | Ocu4$_{ASIP}$ | AX-147170337 | GL018965 | 28277 | | 4.6733E-05 | 2.0436E-05 | 3.1758e-8 | -0.8446 | 0.1507 |
| AX-147097074 | Ocu4$_{ASIP}$ | AX-147136433 | GL018998 | 28677 | | 9.2822E-05 | 3.9785E-05 | 6.8398e-8 | -0.8365 | 0.1531 |
| AX-147097074 | Ocu4$_{ASIP}$ | AX-147078087 | GL018998 | 28678 | | 9.7072E-05 | 4.1512E-05 | 7.1149e-8 | -0.8381 | 0.1536 |
| AX-147097074 | Ocu4$_{ASIP}$ | AX-147173462 | GL018998 | 28680 | | 1.0299E-04 | 4.4025E-05 | 7.7623e-8 | -0.8340 | 0.1533 |
| AX-147097074 | Ocu4$_{ASIP}$ | AX-147175161 | GL018998 | 28686 | | 3.8159E-05 | 1.6517E-05 | 2.1613e-8 | -0.8843 | 0.1557 |
| AX-147097074 | Ocu4$_{ASIP}$ | AX-147168938 | GL018998 | 28687 | | 9.8290E-05 | 4.2070E-05 | 7.3300e-8 | -0.8349 | 0.1532 |
| AX-147097074 | Ocu4$_{ASIP}$ | AX-147173479 | GL019360 | 30535 | | 2.4215E-04 | 1.0067E-04 | 2.0669e-7 | -0.8243 | 0.1568 |
| AX-147097074 | Ocu4$_{ASIP}$ | AX-147159226 | GL019360 | 30536 | | 6.5184E-04 | 2.5705E-04 | 5.1486e-7 | -0.8575 | 0.1689 |
| AX-147097074 | Ocu4$_{ASIP}$ | AX-147173053 | AAGW02081354 | 31035 | | 9.5062E-05 | 4.0670E-05 | 6.9399e-8 | -0.8388 | 0.1536 |

**Table S9: Significant epistatic interaction with the best SNP marker of the GL018965$_{MC1R}$ locus**
Only individuals carrying coloration at extremities, from cream/beige (P3) to dark chocolate/brown (P6) were considered and analysed using an adaptive shrinkage method. lfsr: local false sign rate, lfdr: local false discovery rate, beta: interaction effect and se: standard error

| SNP1 | Location | SNP2 | Ocu | Position | Gene | lfsr | lfdr | pval | beta | se |
|---|---|---|---|---|---|---|---|---|---|---|
| AX-147194100 | GL018965$_{MC1R}$ | AX-147081710 | 2 | 56390169 | | 5.6349E-04 | 2.5898E-04 | 5.0444e-6 | -0.5213 | 0.1132 |
| AX-147194100 | GL018965$_{MC1R}$ | AX-147029724 | 2 | 56466476 | | 6.3814E-04 | 2.9227E-04 | 5.9278e-6 | -0.5173 | 0.1132 |
| AX-147194100 | GL018965$_{MC1R}$ | AX-147052039 | 2 | 56514716 | | 6.4555E-04 | 2.9558E-04 | 6.0201e-6 | -0.5168 | 0.1132 |
| AX-147194100 | GL018965$_{MC1R}$ | AX-147143336 | 2 | 56571740 | | 5.6349E-04 | 2.5898E-04 | 5.0444e-6 | -0.5213 | 0.1132 |
| AX-147194100 | GL018965$_{MC1R}$ | AX-147016688 | 3 | 94335608 | | 9.1234E-04 | 4.1272E-04 | 9.1807e-6 | -0.5090 | 0.1138 |
| AX-147194100 | GL018965$_{MC1R}$ | AX-147167768 | 3 | 96034485 | | 5.5068E-04 | 2.3840E-04 | 1.8340e-6 | 0.6419 | 0.1332 |
| AX-147194100 | GL018965$_{MC1R}$ | AX-147147971 | 3 | 96039991 | | 6.1678E-04 | 2.6610E-04 | 2.1500e-6 | 0.6377 | 0.1332 |
| AX-147194100 | GL018965$_{MC1R}$ | AX-147116661 | 3 | 96095657 | | 4.9667E-04 | 2.1556E-04 | 1.5702e-6 | 0.6469 | 0.1333 |
| AX-147194100 | GL018965$_{MC1R}$ | AX-147044581 | 3 | 96142908 | | 4.5996E-04 | 2.0014E-04 | 1.4181e-6 | 0.6490 | 0.1332 |
| AX-147194100 | GL018965$_{MC1R}$ | AX-146998701 | 3 | 96168690 | | 5.9258E-04 | 2.5588E-04 | 2.0198e-6 | 0.6399 | 0.1333 |
| AX-147194100 | GL018965$_{MC1R}$ | AX-146997500 | 3 | 96232275 | | 4.9667E-04 | 2.1556E-04 | 1.5702e-6 | 0.6469 | 0.1333 |
| AX-147194100 | GL018965$_{MC1R}$ | AX-147007417 | 3 | 96248777 | | 5.2331E-04 | 2.2678E-04 | 1.6916e-6 | 0.6449 | 0.1333 |
| AX-147194100 | GL018965$_{MC1R}$ | AX-147145030 | 3 | 96396410 | | 2.2503E-05 | 1.0842E-05 | 4.0033e-8 | 0.6834 | 0.1228 |
| AX-147194100 | GL018965$_{MC1R}$ | AX-147090801 | 3 | 96476622 | | 4.7115E-04 | 2.0479E-04 | 1.4564e-6 | 0.6490 | 0.1333 |
| AX-147194100 | GL018965$_{MC1R}$ | AX-147088623 | 3 | 96498770 | | 4.7115E-04 | 2.0479E-04 | 1.4564e-6 | 0.6490 | 0.1333 |
| AX-147194100 | GL018965$_{MC1R}$ | AX-147065044 | 3 | 96506770 | | 6.4465E-04 | 2.7472E-04 | 1.8616e-6 | 0.6616 | 0.1374 |
| AX-147194100 | GL018965$_{MC1R}$ | AX-147178006 | 3 | 96513741 | | 3.4842E-05 | 1.6494E-05 | 6.0969e-8 | 0.6893 | 0.1256 |
| AX-147194100 | GL018965$_{MC1R}$ | AX-147039576 | 3 | 96548211 | | 2.9884E-05 | 1.4195E-05 | 4.9902e-8 | 0.6936 | 0.1256 |
| AX-147194100 | GL018965$_{MC1R}$ | AX-147113412 | 3 | 96633386 | | 8.0100E-04 | 3.3943E-04 | 2.5855e-6 | 0.6508 | 0.1371 |
| AX-147194100 | GL018965$_{MC1R}$ | AX-147042096 | 3 | 96709899 | | 9.4570E-04 | 3.9860E-04 | 3.2688e-6 | 0.6445 | 0.1372 |
| AX-147194100 | GL018965$_{MC1R}$ | AX-147062445 | 3 | 96719329 | | 9.4570E-04 | 3.9860E-04 | 3.2688e-6 | 0.6445 | 0.1372 |
| AX-147194100 | GL018965$_{MC1R}$ | AX-146984235 | 3 | 101753403 | *SLC7A13/PSKH2* | 5.1963E-04 | 2.5106E-04 | 7.4820e-6 | 0.4540 | 0.1004 |
| AX-147194100 | GL018965$_{MC1R}$ | AX-147042100 | 3 | 102183637 | *SLC7A13/PSKH2* | 1.9890E-04 | 1.0000E-04 | 2.7428e-6 | 0.4572 | 0.0966 |
| AX-147194100 | GL018965$_{MC1R}$ | AX-147119701 | 3 | 102214044 | *SLC7A13/PSKH2* | 3.2812E-04 | 1.6426E-04 | 5.2165e-6 | 0.4360 | 0.0948 |
| AX-147194100 | GL018965$_{MC1R}$ | AX-147032347 | 3 | 102254753 | *SLC7A13/PSKH2* | 6.0748E-05 | 3.1456E-05 | 7.0041e-7 | 0.4800 | 0.0957 |
| AX-147194100 | GL018965$_{MC1R}$ | AX-147180030 | 3 | 102296886 | *SLC7A13/PSKH2* | 1.0874E-04 | 5.5493E-05 | 1.3631e-6 | 0.4697 | 0.0962 |
| AX-147194100 | GL018965$_{MC1R}$ | AX-146993443 | 3 | 102374529 | | 3.3323E-04 | 1.6675E-04 | 5.3096e-6 | 0.4356 | 0.0948 |
| AX-147194100 | GL018965$_{MC1R}$ | AX-147034694 | 3 | 102382918 | | 3.3706E-04 | 1.6868E-04 | 5.3928e-6 | 0.4350 | 0.0947 |
| AX-147194100 | GL018965$_{MC1R}$ | AX-147074821 | 3 | 102428761 | | 3.7227E-04 | 1.8583E-04 | 6.0470e-6 | 0.4327 | 0.0948 |
| AX-147194100 | GL018965$_{MC1R}$ | AX-147150003 | 3 | 102434864 | | 3.7227E-04 | 1.8583E-04 | 6.0470e-6 | 0.4327 | 0.0948 |
| AX-147194100 | GL018965$_{MC1R}$ | AX-147160043 | 3 | 113264239 | | 7.6122E-04 | 3.5445E-04 | 9.4046e-6 | -0.4791 | 0.1072 |
| AX-147194100 | GL018965$_{MC1R}$ | AX-147102836 | 3 | 113315639 | | 1.1096E-04 | 5.5143E-05 | 1.0432e-6 | 0.5075 | 0.1028 |
| AX-147194100 | GL018965$_{MC1R}$ | AX-147088636 | 3 | 113354882 | | 9.9883E-05 | 4.9752E-05 | 9.1495e-7 | 0.5104 | 0.1029 |
| AX-147194100 | GL018965$_{MC1R}$ | AX-147104696 | 3 | 115785150 | | 7.9621E-05 | 3.9982E-05 | 7.1979e-7 | 0.5113 | 0.1020 |
| AX-147194100 | GL018965$_{MC1R}$ | AX-147171057 | 3 | 115808922 | | 8.3079E-05 | 4.1645E-05 | 7.5044e-7 | 0.5115 | 0.1023 |
| AX-147194100 | GL018965$_{MC1R}$ | AX-147034706 | 3 | 115991989 | | 9.2990E-05 | 4.6503E-05 | 8.6394e-7 | 0.5084 | 0.1022 |
| AX-147194100 | GL018965$_{MC1R}$ | AX-147176700 | 3 | 116044519 | | 9.2990E-05 | 4.6503E-05 | 8.6394e-7 | 0.5084 | 0.1022 |
| AX-147194100 | GL018965$_{MC1R}$ | AX-147126651 | 3 | 116060146 | | 9.2990E-05 | 4.6503E-05 | 8.6394e-7 | 0.5084 | 0.1022 |
| AX-147194100 | GL018965$_{MC1R}$ | AX-147115050 | 3 | 116067589 | | 9.2990E-05 | 4.6503E-05 | 8.6394e-7 | 0.5084 | 0.1022 |
| AX-147194100 | GL018965$_{MC1R}$ | AX-147016706 | 3 | 116072212 | | 9.2990E-05 | 4.6503E-05 | 8.6394e-7 | 0.5084 | 0.1022 |
| AX-147194100 | GL018965$_{MC1R}$ | AX-147130345 | 3 | 116079011 | | 9.2990E-05 | 4.6503E-05 | 8.6394e-7 | 0.5084 | 0.1022 |
| AX-147194100 | GL018965$_{MC1R}$ | AX-147129168 | 3 | 116086979 | | 9.2990E-05 | 4.6503E-05 | 8.6394e-7 | 0.5084 | 0.1022 |
| AX-147194100 | GL018965$_{MC1R}$ | AX-147047239 | 3 | 117898430 | | 1.2654E-04 | 6.1428E-05 | 9.2718e-7 | 0.5379 | 0.1085 |
| AX-147194100 | GL018965$_{MC1R}$ | AX-147116676 | 3 | 117925678 | | 1.2654E-04 | 6.1428E-05 | 9.2718e-7 | 0.5379 | 0.1085 |
| AX-147194100 | GL018965$_{MC1R}$ | AX-147153108 | 4 | 3758580 | | 8.9617E-05 | 3.8679E-05 | 7.0686e-8 | -0.8230 | 0.1508 |
| AX-147194100 | GL018965$_{MC1R}$ | AX-146984655 | 4 | 3768175 | | 7.3819E-05 | 3.2113E-05 | 5.8193e-8 | -0.8206 | 0.1494 |
| AX-147194100 | GL018965$_{MC1R}$ | AX-147074883 | 4 | 4154002 | | 8.5008E-04 | 3.3722E-04 | 9.0204e-7 | -0.8106 | 0.1633 |
| AX-147194100 | GL018965$_{MC1R}$ | AX-147016737 | 4 | 4860188 | | 2.9473E-04 | 1.2097E-04 | 2.3774e-7 | -0.8371 | 0.1601 |
| AX-147194100 | GL018965$_{MC1R}$ | AX-147029987 | 4 | 4873287 | | 2.9473E-04 | 1.2097E-04 | 2.3774e-7 | -0.8371 | 0.1601 |

| | | | | | | | | | | |
|---|---|---|---|---|---|---|---|---|---|---|
| AX-147194100 | GL018965$_{MC1R}$ | AX-147161027 | 4 | 4885152 | | 2.7613E-04 | 1.1401E-04 | 2.3190e-7 | -0.8287 | 0.1584 |
| AX-147194100 | GL018965$_{MC1R}$ | AX-147072521 | 4 | 4897773 | | 2.7613E-04 | 1.1401E-04 | 2.3190e-7 | -0.8287 | 0.1584 |
| AX-147194100 | GL018965$_{MC1R}$ | AX-147025249 | 4 | 4972440 | | 2.7613E-04 | 1.1401E-04 | 2.3190e-7 | -0.8287 | 0.1584 |
| AX-147194100 | GL018965$_{MC1R}$ | AX-147077276 | 4 | 4981328 | | 2.7613E-04 | 1.1401E-04 | 2.3190e-7 | -0.8287 | 0.1584 |
| AX-147194100 | GL018965$_{MC1R}$ | AX-147088674 | 4 | 4988564 | | 2.7613E-04 | 1.1401E-04 | 2.3190e-7 | -0.8287 | 0.1584 |
| AX-147194100 | GL018965$_{MC1R}$ | AX-147127978 | 4 | 4994098 | | 2.9473E-04 | 1.2097E-04 | 2.3774e-7 | -0.8371 | 0.1601 |
| AX-147194100 | GL018965$_{MC1R}$ | AX-147168744 | 4 | 5108085 | | 5.4728E-06 | 2.8171E-06 | 1.3846e-8 | -0.6353 | 0.1104 |
| AX-147194100 | GL018965$_{MC1R}$ | AX-147070051 | 4 | 5111463 | | 5.7656E-06 | 2.9648E-06 | 1.4792e-8 | -0.6342 | 0.1104 |
| AX-147194100 | GL018965$_{MC1R}$ | AX-147175834 | 4 | 5172769 | | 1.2812E-06 | 6.8016E-07 | 2.4792e-9 | -0.6561 | 0.1084 |
| AX-147194100 | GL018965$_{MC1R}$ | AX-147141770 | 4 | 5355432 | | 2.2529E-04 | 9.3525E-05 | 1.7952e-7 | -0.8347 | 0.1580 |
| AX-147194100 | GL018965$_{MC1R}$ | AX-147156483 | 4 | 5412648 | | 7.1256E-09 | 3.9766E-09 | 6.1243e-12 | -0.7829 | 0.1116 |
| AX-147194100 | GL018965$_{MC1R}$ | AX-147029988 | 4 | 5429684 | | 5.3895E-09 | 3.0376E-09 | 4.8015e-12 | -0.7731 | 0.1096 |
| AX-147194100 | GL018965$_{MC1R}$ | AX-146983875 | 4 | 5435197 | *ASIP* | 4.6420E-08 | 2.5238E-08 | 4.0516e-11 | -0.7647 | 0.1137 |
| AX-147194100 | GL018965$_{MC1R}$ | AX-146983878 | 4 | 5435206 | *ASIP* | 2.4131E-07 | 1.2966E-07 | 2.6829e-10 | -0.7176 | 0.1117 |
| AX-147194100 | GL018965$_{MC1R}$ | AS-P5435370 | 4 | 5435370 | *ASIP* | 3.6618E-08 | 2.0236E-08 | 3.5190e-11 | -0.7417 | 0.1099 |
| AX-147194100 | GL018965$_{MC1R}$ | AX-146983848 | 4 | 5435391 | *ASIP* | 1.3906E-04 | 6.0846E-05 | 1.7189e-7 | -0.7533 | 0.1424 |
| AX-147194100 | GL018965$_{MC1R}$ | ASIP | 4 | 5435400 | *ASIP* | 1.9445E-09 | 1.1152E-09 | 1.7978e-12 | -0.7728 | 0.1073 |
| AX-147194100 | GL018965$_{MC1R}$ | AX-146983571 | 4 | 5439656 | *ASIP* | 4.8689E-08 | 2.6687E-08 | 4.5687e-11 | -0.7467 | 0.1113 |
| AX-147194100 | GL018965$_{MC1R}$ | AX-146983570 | 4 | 5439677 | *ASIP* | 4.8689E-08 | 2.6687E-08 | 4.5687e-11 | -0.7467 | 0.1113 |
| AX-147194100 | GL018965$_{MC1R}$ | AX-147090864 | 4 | 5501377 | | 2.4818E-09 | 1.4202E-09 | 2.2947e-12 | -0.7692 | 0.1073 |
| AX-147194100 | GL018965$_{MC1R}$ | AX-147047284 | 4 | 5550419 | | 1.4671E-09 | 8.4288E-10 | 1.3541e-12 | -0.7783 | 0.1074 |
| AX-147194100 | GL018965$_{MC1R}$ | AX-147001274 | 4 | 5599202 | | 7.2197E-10 | 4.2043E-10 | 6.9472e-13 | -0.7736 | 0.1054 |
| AX-147194100 | GL018965$_{MC1R}$ | AX-147059966 | 4 | 5602431 | | 7.2197E-10 | 4.2043E-10 | 6.9472e-13 | -0.7736 | 0.1054 |
| AX-147194100 | GL018965$_{MC1R}$ | AX-147059967 | 4 | 5633343 | | 9.6466E-10 | 5.6080E-10 | 9.2821e-13 | -0.7680 | 0.1052 |
| AX-147194100 | GL018965$_{MC1R}$ | AX-147115077 | 4 | 5647773 | | 9.4079E-08 | 5.2275E-08 | 1.2397e-10 | -0.6908 | 0.1054 |
| AX-147194100 | GL018965$_{MC1R}$ | AX-147082020 | 4 | 5655567 | | 7.2197E-10 | 4.2043E-10 | 6.9472e-13 | -0.7736 | 0.1054 |
| AX-147194100 | GL018965$_{MC1R}$ | AX-147136018 | 4 | 5660694 | | 1.5906E-09 | 9.1928E-10 | 1.5159e-12 | -0.7638 | 0.1057 |
| AX-147194100 | GL018965$_{MC1R}$ | AX-147161987 | 4 | 5665991 | | 1.3814E-10 | 8.1246E-11 | 1.3750e-13 | -0.8030 | 0.1060 |
| AX-147194100 | GL018965$_{MC1R}$ | AX-147132705 | 4 | 5673715 | | 7.2197E-10 | 4.2043E-10 | 6.9472e-13 | -0.7736 | 0.1054 |
| AX-147194100 | GL018965$_{MC1R}$ | AX-147067643 | 4 | 5701317 | | 6.7587E-10 | 3.9350E-10 | 6.4886e-13 | -0.7760 | 0.1056 |
| AX-147194100 | GL018965$_{MC1R}$ | AX-147129189 | 4 | 5710216 | | 9.1582E-10 | 5.3264E-10 | 8.8155e-13 | -0.7688 | 0.1052 |
| AX-147194100 | GL018965$_{MC1R}$ | AX-147022988 | 4 | 5732438 | | 9.1582E-10 | 5.3264E-10 | 8.8155e-13 | -0.7688 | 0.1052 |
| AX-147194100 | GL018965$_{MC1R}$ | AX-147168745 | 4 | 5749725 | | 9.6553E-10 | 5.6107E-10 | 9.2725e-13 | -0.7688 | 0.1053 |
| AX-147194100 | GL018965$_{MC1R}$ | AX-147034762 | 4 | 5776903 | | 9.3635E-10 | 5.4454E-10 | 9.0166e-13 | -0.7682 | 0.1052 |
| AX-147194100 | GL018965$_{MC1R}$ | AX-147097073 | 4 | 5831757 | | 7.2163E-10 | 4.2025E-10 | 6.9446e-13 | -0.7735 | 0.1054 |
| AX-147194100 | GL018965$_{MC1R}$ | AX-147044647 | 4 | 5843986 | | 7.8686E-10 | 4.5638E-10 | 7.4546e-13 | -0.7790 | 0.1063 |
| AX-147194100 | GL018965$_{MC1R}$ | AX-147067644 | 4 | 5850185 | | 7.2163E-10 | 4.2025E-10 | 6.9446e-13 | -0.7735 | 0.1054 |
| AX-147194100 | GL018965$_{MC1R}$ | AX-147110148 | 4 | 5857504 | | 1.3313E-10 | 7.8325E-11 | 1.3270e-13 | -0.8035 | 0.1060 |
| AX-147194100 | GL018965$_{MC1R}$ | AX-147164912 | 4 | 5862771 | | 1.7783E-10 | 1.0448E-10 | 1.7589e-13 | -0.7978 | 0.1058 |
| AX-147194100 | GL018965$_{MC1R}$ | AX-147119738 | 4 | 5872863 | | 9.3635E-10 | 5.4454E-10 | 9.0166e-13 | -0.7682 | 0.1052 |
| AX-147194100 | GL018965$_{MC1R}$ | AX-147174144 | 4 | 5897338 | | 1.6945E-10 | 9.9518E-11 | 1.6746e-13 | -0.7999 | 0.1060 |
| AX-147194100 | GL018965$_{MC1R}$ | AX-147175406 | 4 | 5903530 | | 1.7783E-10 | 1.0448E-10 | 1.7589e-13 | -0.7978 | 0.1058 |
| AX-147194100 | GL018965$_{MC1R}$ | AX-147151895 | 4 | 5958610 | | 9.3934E-10 | 5.4432E-10 | 8.8986e-13 | -0.7753 | 0.1061 |
| AX-147194100 | GL018965$_{MC1R}$ | AX-147153729 | 4 | 5993707 | | 2.7931E-08 | 1.5542E-08 | 2.7261e-11 | -0.7386 | 0.1088 |
| AX-147194100 | GL018965$_{MC1R}$ | AX-147113450 | 4 | 6047198 | | 8.1032E-09 | 4.5364E-09 | 7.1496e-12 | -0.7722 | 0.1104 |
| AX-147194100 | GL018965$_{MC1R}$ | AX-147133844 | 4 | 6060380 | | 3.7290E-09 | 2.1098E-09 | 3.3198e-12 | -0.7781 | 0.1094 |
| AX-147194100 | GL018965$_{MC1R}$ | AX-147018712 | 4 | 6066489 | | 7.5272E-09 | 4.2329E-09 | 6.7808e-12 | -0.7659 | 0.1094 |
| AX-147194100 | GL018965$_{MC1R}$ | AX-147131545 | 4 | 6132488 | | 2.7931E-08 | 1.5542E-08 | 2.7261e-11 | -0.7386 | 0.1088 |
| AX-147194100 | GL018965$_{MC1R}$ | AX-147072522 | 4 | 6140446 | | 2.7931E-08 | 1.5542E-08 | 2.7261e-11 | -0.7386 | 0.1088 |
| AX-147194100 | GL018965$_{MC1R}$ | AX-147111822 | 4 | 6162134 | | 7.5272E-09 | 4.2329E-09 | 6.7808e-12 | -0.7659 | 0.1094 |
| AX-147194100 | GL018965$_{MC1R}$ | AX-147148704 | 4 | 6178276 | | 9.5506E-05 | 4.0781E-05 | 6.8109e-8 | -0.8430 | 0.1543 |
| AX-147194100 | GL018965$_{MC1R}$ | AX-147104722 | 4 | 6247254 | | 9.5506E-05 | 4.0781E-05 | 6.8109e-8 | -0.8430 | 0.1543 |
| AX-147194100 | GL018965$_{MC1R}$ | AX-147118241 | 4 | 6256260 | | 9.5506E-05 | 4.0781E-05 | 6.8109e-8 | -0.8430 | 0.1543 |
| AX-147194100 | GL018965$_{MC1R}$ | AX-147116701 | 4 | 6263090 | | 1.5595E-04 | 6.5886E-05 | 1.2638e-7 | -0.8258 | 0.1544 |
| AX-147194100 | GL018965$_{MC1R}$ | AX-147178424 | 4 | 6299192 | | 9.9959E-05 | 4.2370E-05 | 6.6595e-8 | -0.8569 | 0.1567 |

| | | | | | | | | | |
|---|---|---|---|---|---|---|---|---|---|
| AX-147194100 | GL018965$_{MC1R}$ | AX-147180423 | 4 | 6409251 | 1.5761E-04 | 6.7196E-05 | 1.4519e-7 | -0.8031 | 0.1509 |
| AX-147194100 | GL018965$_{MC1R}$ | AX-147086481 | 4 | 6571042 | 1.6581E-04 | 7.2089E-05 | 2.0990e-7 | -0.7523 | 0.1432 |
| AX-147194100 | GL018965$_{MC1R}$ | AX-147133845 | 4 | 6583970 | 1.0125E-04 | 4.4570E-05 | 1.1112e-7 | -0.7672 | 0.1428 |
| AX-147194100 | GL018965$_{MC1R}$ | AX-147047285 | 4 | 6609333 | 1.6581E-04 | 7.2089E-05 | 2.0990e-7 | -0.7523 | 0.1432 |
| AX-147194100 | GL018965$_{MC1R}$ | AX-147138110 | 4 | 6621523 | 2.6459E-04 | 1.1409E-04 | 4.1838e-7 | -0.7272 | 0.1421 |
| AX-147194100 | GL018965$_{MC1R}$ | AX-147002769 | 4 | 6628954 | 1.6581E-04 | 7.2089E-05 | 2.0990e-7 | -0.7523 | 0.1432 |
| AX-147194100 | GL018965$_{MC1R}$ | AX-147108384 | 4 | 6665508 | 6.3833E-09 | 3.6172E-09 | 5.9492e-12 | -0.7566 | 0.1077 |
| AX-147194100 | GL018965$_{MC1R}$ | AX-147141771 | 4 | 6691032 | 6.3833E-09 | 3.6172E-09 | 5.9492e-12 | -0.7566 | 0.1077 |
| AX-147194100 | GL018965$_{MC1R}$ | AX-147165829 | 4 | 6832350 | 1.4097E-09 | 8.0659E-10 | 1.2758e-12 | -0.7878 | 0.1086 |
| AX-147194100 | GL018965$_{MC1R}$ | AX-147111824 | 4 | 6837895 | 6.3833E-09 | 3.6172E-09 | 5.9492e-12 | -0.7566 | 0.1077 |
| AX-147194100 | GL018965$_{MC1R}$ | AX-147145039 | 4 | 6845381 | 1.3152E-09 | 7.5353E-10 | 1.1953e-12 | -0.7873 | 0.1084 |
| AX-147194100 | GL018965$_{MC1R}$ | AX-147169236 | 4 | 6852795 | 1.0902E-09 | 6.2553E-10 | 9.9293e-13 | -0.7904 | 0.1084 |
| AX-147194100 | GL018965$_{MC1R}$ | AX-147042159 | 4 | 6867188 | 6.3833E-09 | 3.6172E-09 | 5.9492e-12 | -0.7566 | 0.1077 |
| AX-147194100 | GL018965$_{MC1R}$ | AX-147166292 | 4 | 6910263 | 6.3833E-09 | 3.6172E-09 | 5.9492e-12 | -0.7566 | 0.1077 |
| AX-147194100 | GL018965$_{MC1R}$ | AX-147174991 | 4 | 6945657 | 6.3833E-09 | 3.6172E-09 | 5.9492e-12 | -0.7566 | 0.1077 |
| AX-147194100 | GL018965$_{MC1R}$ | AX-147054938 | 4 | 6952681 | 4.9970E-09 | 2.8360E-09 | 4.6091e-12 | -0.7620 | 0.1079 |
| AX-147194100 | GL018965$_{MC1R}$ | AX-147090865 | 4 | 7034409 | 5.2565E-09 | 2.9648E-09 | 4.6944e-12 | -0.7726 | 0.1095 |
| AX-147194100 | GL018965$_{MC1R}$ | AX-147141772 | 4 | 7057509 | 6.3833E-09 | 3.6172E-09 | 5.9492e-12 | -0.7566 | 0.1077 |
| AX-147194100 | GL018965$_{MC1R}$ | AX-147144266 | 4 | 7076001 | 6.0774E-09 | 3.4426E-09 | 5.6272e-12 | -0.7590 | 0.1079 |
| AX-147194100 | GL018965$_{MC1R}$ | AX-147079699 | 4 | 7081578 | 4.8288E-09 | 2.7401E-09 | 4.4387e-12 | -0.7634 | 0.1080 |
| AX-147194100 | GL018965$_{MC1R}$ | AX-147150659 | 4 | 7093781 | 4.9970E-09 | 2.8360E-09 | 4.6091e-12 | -0.7620 | 0.1079 |
| AX-147194100 | GL018965$_{MC1R}$ | AX-147169681 | 4 | 7186175 | 4.8201E-04 | 1.9787E-04 | 5.5324e-7 | -0.7859 | 0.1552 |
| AX-147194100 | GL018965$_{MC1R}$ | AX-147027565 | 4 | 7251253 | 2.3056E-08 | 1.2693E-08 | 2.0167e-11 | -0.7671 | 0.1122 |
| AX-147194100 | GL018965$_{MC1R}$ | AX-147170603 | 4 | 7264726 | 2.8500E-08 | 1.5655E-08 | 2.5159e-11 | -0.7634 | 0.1122 |
| AX-147194100 | GL018965$_{MC1R}$ | AX-147177567 | 4 | 7280887 | 2.8500E-08 | 1.5655E-08 | 2.5159e-11 | -0.7634 | 0.1122 |
| AX-147194100 | GL018965$_{MC1R}$ | AX-147170145 | 4 | 7286249 | 2.8500E-08 | 1.5655E-08 | 2.5159e-11 | -0.7634 | 0.1122 |
| AX-147194100 | GL018965$_{MC1R}$ | AX-147133846 | 4 | 7293733 | 3.1786E-08 | 1.7440E-08 | 2.8218e-11 | -0.7614 | 0.1122 |
| AX-147194100 | GL018965$_{MC1R}$ | AX-147054939 | 4 | 7299811 | 3.1786E-08 | 1.7440E-08 | 2.8218e-11 | -0.7614 | 0.1122 |
| AX-147194100 | GL018965$_{MC1R}$ | AX-147123987 | 4 | 7321664 | 3.1786E-08 | 1.7440E-08 | 2.8218e-11 | -0.7614 | 0.1122 |
| AX-147194100 | GL018965$_{MC1R}$ | AX-147133847 | 4 | 7327831 | 2.5871E-07 | 1.3819E-07 | 2.7274e-10 | -0.7264 | 0.1131 |
| AX-147194100 | GL018965$_{MC1R}$ | AX-147179605 | 4 | 7347783 | 3.1786E-08 | 1.7440E-08 | 2.8218e-11 | -0.7614 | 0.1122 |
| AX-147194100 | GL018965$_{MC1R}$ | AX-147150017 | 4 | 7354537 | 3.1786E-08 | 1.7440E-08 | 2.8218e-11 | -0.7614 | 0.1122 |
| AX-147194100 | GL018965$_{MC1R}$ | AX-147100956 | 4 | 7429282 | 5.4628E-05 | 2.6954E-05 | 2.9932e-7 | -0.5667 | 0.1093 |
| AX-147194100 | GL018965$_{MC1R}$ | AX-147097075 | 4 | 7451409 | 1.1081E-04 | 5.3494E-05 | 6.8080e-7 | -0.5574 | 0.1110 |
| AX-147194100 | GL018965$_{MC1R}$ | AX-147065114 | 4 | 7468597 | 5.4628E-05 | 2.6954E-05 | 2.9932e-7 | -0.5667 | 0.1093 |
| AX-147194100 | GL018965$_{MC1R}$ | AX-147047286 | 4 | 7485053 | 5.4628E-05 | 2.6954E-05 | 2.9932e-7 | -0.5667 | 0.1093 |
| AX-147194100 | GL018965$_{MC1R}$ | AX-147147994 | 4 | 7504341 | 5.4628E-05 | 2.6954E-05 | 2.9932e-7 | -0.5667 | 0.1093 |
| AX-147194100 | GL018965$_{MC1R}$ | AX-147118242 | 4 | 7523401 | 1.0061E-04 | 4.5683E-05 | 1.8180e-7 | -0.6964 | 0.1319 |
| AX-147194100 | GL018965$_{MC1R}$ | AX-147136019 | 4 | 7536509 | 1.0061E-04 | 4.5683E-05 | 1.8180e-7 | -0.6964 | 0.1319 |
| AX-147194100 | GL018965$_{MC1R}$ | AX-147168747 | 4 | 7554760 | 1.0061E-04 | 4.5683E-05 | 1.8180e-7 | -0.6964 | 0.1319 |
| AX-147194100 | GL018965$_{MC1R}$ | AX-147166806 | 4 | 7563636 | 1.0061E-04 | 4.5683E-05 | 1.8180e-7 | -0.6964 | 0.1319 |
| AX-147194100 | GL018965$_{MC1R}$ | AX-147146554 | 4 | 7578427 | 1.4390E-04 | 6.4170E-05 | 2.4981e-7 | -0.7060 | 0.1353 |
| AX-147194100 | GL018965$_{MC1R}$ | AX-147159539 | 4 | 7586674 | 2.0021E-04 | 8.6128E-05 | 2.4601e-7 | -0.7597 | 0.1455 |
| AX-147194100 | GL018965$_{MC1R}$ | AX-147093035 | 4 | 7822042 | 7.0594E-05 | 3.4663E-05 | 4.2346e-7 | -0.5580 | 0.1091 |
| AX-147194100 | GL018965$_{MC1R}$ | AX-147088683 | 4 | 17259391 | 1.8955E-04 | 9.0947E-05 | 1.5113e-6 | 0.5297 | 0.1090 |
| AX-147194100 | GL018965$_{MC1R}$ | AX-147059974 | 4 | 17266266 | 1.8955E-04 | 9.0947E-05 | 1.5113e-6 | 0.5297 | 0.1090 |
| AX-147194100 | GL018965$_{MC1R}$ | AX-147169684 | 4 | 17278633 | 1.8955E-04 | 9.0947E-05 | 1.5113e-6 | 0.5297 | 0.1090 |
| AX-147194100 | GL018965$_{MC1R}$ | AX-147097087 | 4 | 17288318 | 1.8955E-04 | 9.0947E-05 | 1.5113e-6 | 0.5297 | 0.1090 |
| AX-147194100 | GL018965$_{MC1R}$ | AX-147099072 | 4 | 17303972 | 1.8955E-04 | 9.0947E-05 | 1.5113e-6 | 0.5297 | 0.1090 |
| AX-147194100 | GL018965$_{MC1R}$ | AX-147172397 | 4 | 17310761 | 2.3683E-04 | 1.1270E-04 | 1.9342e-6 | 0.5279 | 0.1098 |
| AX-147194100 | GL018965$_{MC1R}$ | AX-147016740 | 4 | 17319356 | 2.4040E-04 | 1.1460E-04 | 2.0309e-6 | 0.5238 | 0.1092 |
| AX-147194100 | GL018965$_{MC1R}$ | AX-147062504 | 4 | 17330905 | 1.8955E-04 | 9.0947E-05 | 1.5113e-6 | 0.5297 | 0.1090 |
| AX-147194100 | GL018965$_{MC1R}$ | AX-147137046 | 4 | 17491611 | 2.0833E-04 | 9.9723E-05 | 1.7037e-6 | 0.5271 | 0.1090 |
| AX-147194100 | GL018965$_{MC1R}$ | AX-147002775 | 4 | 17496614 | 2.4706E-04 | 1.1774E-04 | 2.1135e-6 | 0.5224 | 0.1091 |
| AX-147194100 | GL018965$_{MC1R}$ | AX-147106569 | 4 | 17573103 | 2.0833E-04 | 9.9723E-05 | 1.7037e-6 | 0.5271 | 0.1090 |

| | | | | | | | | | | |
|---|---|---|---|---|---|---|---|---|---|---|
| AX-147194100 | GL018965$_{MC1R}$ | AX-147095071 | 4 | 17645970 | | 2.1536E-04 | 1.0300E-04 | 1.7774e-6 | 0.5261 | 0.1090 |
| AX-147194100 | GL018965$_{MC1R}$ | AX-147062505 | 4 | 17653776 | | 2.5518E-04 | 1.2149E-04 | 2.1969e-6 | 0.5218 | 0.1091 |
| AX-147194100 | GL018965$_{MC1R}$ | AX-147140921 | 4 | 17666468 | | 2.1536E-04 | 1.0300E-04 | 1.7774e-6 | 0.5261 | 0.1090 |
| AX-147194100 | GL018965$_{MC1R}$ | AX-147088684 | 4 | 17688360 | | 2.1536E-04 | 1.0300E-04 | 1.7774e-6 | 0.5261 | 0.1090 |
| AX-147194100 | GL018965$_{MC1R}$ | AX-147175408 | 4 | 17736623 | | 2.2554E-04 | 1.0777E-04 | 1.8905e-6 | 0.5245 | 0.1090 |
| AX-147194100 | GL018965$_{MC1R}$ | AX-147082029 | 4 | 17766006 | | 1.9595E-04 | 9.3941E-05 | 1.5768e-6 | 0.5288 | 0.1090 |
| AX-147194100 | GL018965$_{MC1R}$ | AX-147088685 | 4 | 17791678 | | 1.9595E-04 | 9.3941E-05 | 1.5768e-6 | 0.5288 | 0.1090 |
| AX-147194100 | GL018965$_{MC1R}$ | AX-147131555 | 4 | 17832711 | | 1.9595E-04 | 9.3941E-05 | 1.5768e-6 | 0.5288 | 0.1090 |
| AX-147194100 | GL018965$_{MC1R}$ | AX-147027572 | 4 | 17855444 | | 1.9963E-04 | 9.5656E-05 | 1.6127e-6 | 0.5284 | 0.1090 |
| AX-147194100 | GL018965$_{MC1R}$ | AX-147108392 | 4 | 17993039 | | 1.9595E-04 | 9.3941E-05 | 1.5768e-6 | 0.5288 | 0.1090 |
| AX-147194100 | GL018965$_{MC1R}$ | AX-147067655 | 4 | 18155509 | | 1.9595E-04 | 9.3941E-05 | 1.5768e-6 | 0.5288 | 0.1090 |
| AX-147194100 | GL018965$_{MC1R}$ | AX-147161484 | 4 | 18464637 | | 2.2291E-04 | 1.1790E-04 | 4.3768e-6 | 0.3937 | 0.0849 |
| AX-147194100 | GL018965$_{MC1R}$ | AX-147150664 | 4 | 19197281 | | 2.0388E-04 | 9.7648E-05 | 1.6582e-6 | 0.5277 | 0.1090 |
| AX-147194100 | GL018965$_{MC1R}$ | AX-147025254 | 4 | 19542374 | | 2.6372E-04 | 1.2564E-04 | 2.3383e-6 | 0.5182 | 0.1087 |
| AX-147194100 | GL018965$_{MC1R}$ | AX-147129198 | 4 | 19581862 | | 2.3596E-04 | 1.1271E-04 | 2.0254e-6 | 0.5217 | 0.1087 |
| AX-147194100 | GL018965$_{MC1R}$ | AX-147025255 | 4 | 19609491 | | 2.4010E-04 | 1.1467E-04 | 2.0773e-6 | 0.5208 | 0.1087 |
| AX-147194100 | GL018965$_{MC1R}$ | AX-147145041 | 4 | 19615377 | | 2.4010E-04 | 1.1467E-04 | 2.0773e-6 | 0.5208 | 0.1087 |
| AX-147194100 | GL018965$_{MC1R}$ | AX-147095072 | 4 | 19639397 | | 2.4010E-04 | 1.1467E-04 | 2.0773e-6 | 0.5208 | 0.1087 |
| AX-147194100 | GL018965$_{MC1R}$ | AX-147136027 | 4 | 19667206 | | 2.4010E-04 | 1.1467E-04 | 2.0773e-6 | 0.5208 | 0.1087 |
| AX-147194100 | GL018965$_{MC1R}$ | AX-147018721 | 4 | 19675540 | | 2.4730E-04 | 1.1802E-04 | 2.1559e-6 | 0.5200 | 0.1087 |
| AX-147194100 | GL018965$_{MC1R}$ | AX-147138117 | 4 | 19685108 | | 2.4010E-04 | 1.1467E-04 | 2.0773e-6 | 0.5208 | 0.1087 |
| AX-147194100 | GL018965$_{MC1R}$ | AX-147110161 | 4 | 19745235 | | 2.6372E-04 | 1.2564E-04 | 2.3383e-6 | 0.5182 | 0.1087 |
| AX-147194100 | GL018965$_{MC1R}$ | AX-147016741 | 4 | 19754842 | | 2.6372E-04 | 1.2564E-04 | 2.3383e-6 | 0.5182 | 0.1087 |
| AX-147194100 | GL018965$_{MC1R}$ | AX-147111835 | 4 | 19804192 | | 2.4010E-04 | 1.1467E-04 | 2.0773e-6 | 0.5208 | 0.1087 |
| AX-147194100 | GL018965$_{MC1R}$ | AX-147039678 | 4 | 19865298 | | 2.4010E-04 | 1.1467E-04 | 2.0773e-6 | 0.5208 | 0.1087 |
| AX-147194100 | GL018965$_{MC1R}$ | AX-147054953 | 4 | 19878939 | | 3.2724E-04 | 1.5553E-04 | 3.1931e-6 | 0.5071 | 0.1078 |
| AX-147194100 | GL018965$_{MC1R}$ | AX-147143427 | 4 | 19904847 | | 2.4010E-04 | 1.1467E-04 | 2.0773e-6 | 0.5208 | 0.1087 |
| AX-147194100 | GL018965$_{MC1R}$ | AX-147159543 | 4 | 19932860 | | 2.4010E-04 | 1.1467E-04 | 2.0773e-6 | 0.5208 | 0.1087 |
| AX-147194100 | GL018965$_{MC1R}$ | AX-147097089 | 4 | 19970204 | | 1.6107E-04 | 7.7599E-05 | 1.2291e-6 | 0.5342 | 0.1090 |
| AX-147194100 | GL018965$_{MC1R}$ | AX-147042174 | 4 | 19982775 | | 2.4010E-04 | 1.1467E-04 | 2.0773e-6 | 0.5208 | 0.1087 |
| AX-147194100 | GL018965$_{MC1R}$ | AX-147065121 | 4 | 20133915 | | 1.9837E-04 | 9.5169E-05 | 1.6248e-6 | 0.5267 | 0.1087 |
| AX-147194100 | GL018965$_{MC1R}$ | AX-147034773 | 4 | 20139374 | | 1.8338E-04 | 8.8132E-05 | 1.4651e-6 | 0.5293 | 0.1088 |
| AX-147194100 | GL018965$_{MC1R}$ | AX-147116710 | 4 | 20147777 | | 4.9765E-04 | 2.3261E-04 | 5.0740e-6 | 0.5034 | 0.1093 |
| AX-147194100 | GL018965$_{MC1R}$ | AX-147130380 | 4 | 41621806 | | 7.8337E-04 | 3.5568E-04 | 7.4655e-6 | 0.5153 | 0.1140 |
| AX-147194100 | GL018965$_{MC1R}$ | AX-147044679 | 4 | 42136130 | | 8.1980E-04 | 3.7359E-04 | 8.4009e-6 | 0.5058 | 0.1125 |
| AX-147194100 | GL018965$_{MC1R}$ | AX-147070078 | 4 | 42208014 | | 8.1980E-04 | 3.7359E-04 | 8.4009e-6 | 0.5058 | 0.1125 |
| AX-147194100 | GL018965$_{MC1R}$ | AX-147072559 | 4 | 42216083 | | 8.1980E-04 | 3.7359E-04 | 8.4009e-6 | 0.5058 | 0.1125 |
| AX-147194100 | GL018965$_{MC1R}$ | AX-147018733 | 4 | 42236407 | | 8.1980E-04 | 3.7359E-04 | 8.4009e-6 | 0.5058 | 0.1125 |
| AX-147194100 | GL018965$_{MC1R}$ | AX-147152508 | 4 | 42267703 | | 9.0257E-04 | 4.1014E-04 | 9.5009e-6 | 0.5028 | 0.1125 |
| AX-147194100 | GL018965$_{MC1R}$ | AX-147018734 | 4 | 42283609 | | 7.2064E-04 | 3.2952E-04 | 7.0908e-6 | 0.5104 | 0.1126 |
| AX-147194100 | GL018965$_{MC1R}$ | AX-147074900 | 4 | 42302021 | | 8.1980E-04 | 3.7359E-04 | 8.4009e-6 | 0.5058 | 0.1125 |
| AX-147194100 | GL018965$_{MC1R}$ | AX-147138126 | 4 | 42308759 | | 6.7148E-04 | 3.0758E-04 | 6.4538e-6 | 0.5131 | 0.1127 |
| AX-147194100 | GL018965$_{MC1R}$ | AX-147020878 | 4 | 42322040 | | 8.1980E-04 | 3.7359E-04 | 8.4009e-6 | 0.5058 | 0.1125 |
| AX-147194100 | GL018965$_{MC1R}$ | AX-147079727 | 4 | 42335738 | | 8.1980E-04 | 3.7359E-04 | 8.4009e-6 | 0.5058 | 0.1125 |
| AX-147194100 | GL018965$_{MC1R}$ | AX-147074901 | 4 | 42343074 | | 8.1980E-04 | 3.7359E-04 | 8.4009e-6 | 0.5058 | 0.1125 |
| AX-147194100 | GL018965$_{MC1R}$ | AX-147121222 | 4 | 42423613 | | 8.1980E-04 | 3.7359E-04 | 8.4009e-6 | 0.5058 | 0.1125 |
| AX-147194100 | GL018965$_{MC1R}$ | AX-147147251 | 4 | 42594926 | | 7.2520E-04 | 3.3165E-04 | 7.1749e-6 | 0.5097 | 0.1125 |
| AX-147194100 | GL018965$_{MC1R}$ | AX-147093064 | 4 | 57056982 | | 5.4260E-04 | 2.5092E-04 | 5.1255e-6 | 0.5140 | 0.1117 |
| AX-147194100 | GL018965$_{MC1R}$ | AX-147145848 | 4 | 57392538 | | 9.4033E-04 | 4.2682E-04 | 1.0021e-5 | 0.5014 | 0.1125 |
| AX-147194100 | GL018965$_{MC1R}$ | AX-147093065 | 4 | 57399288 | | 4.5593E-04 | 2.1184E-04 | 4.0962e-6 | 0.5195 | 0.1117 |
| AX-147194100 | GL018965$_{MC1R}$ | AX-147077314 | 4 | 57404747 | | 4.3117E-04 | 2.0066E-04 | 3.8172e-6 | 0.5210 | 0.1117 |
| AX-147194100 | GL018965$_{MC1R}$ | AX-147168255 | 4 | 57475952 | | 6.8725E-04 | 3.0817E-04 | 5.0553e-6 | 0.5476 | 0.1189 |
| AX-147194100 | GL018965$_{MC1R}$ | AX-147099194 | 7 | 18487673 | *POT1* | 6.7558E-04 | 3.4048E-04 | 1.3660e-5 | 0.3925 | 0.0895 |
| AX-147194100 | GL018965$_{MC1R}$ | AX-147144315 | 7 | 18935180 | *POT1* | 6.7439E-04 | 3.4159E-04 | 1.3927e-5 | 0.3877 | 0.0885 |
| AX-147194100 | GL018965$_{MC1R}$ | AX-147052496 | 7 | 19777217 | *POT1* | 7.9912E-04 | 3.8152E-04 | 1.2470e-5 | -0.4431 | 0.1005 |

| | | | | | | | | | | |
|---|---|---|---|---|---|---|---|---|---|---|
| AX-147194100 | GL018965$_{MC1R}$ | AX-146983344 | 7 | 23565806 | | 5.9016E-04 | 2.9842E-04 | 1.1724e-5 | 0.3957 | 0.0895 |
| AX-147194100 | GL018965$_{MC1R}$ | AX-147124124 | 7 | 122231177 | *CERKL* | 2.1092E-04 | 1.0309E-04 | 2.2563e-6 | 0.4932 | 0.1032 |
| AX-147194100 | GL018965$_{MC1R}$ | AX-146989746 | 7 | 122281634 | *CERKL* | 4.2737E-04 | 2.0993E-04 | 6.4921e-6 | 0.4446 | 0.0977 |
| AX-147194100 | GL018965$_{MC1R}$ | AX-147042415 | 7 | 122307378 | *CERKL* | 4.2737E-04 | 2.0993E-04 | 6.4921e-6 | 0.4446 | 0.0977 |
| AX-147194100 | GL018965$_{MC1R}$ | AX-147104912 | 7 | 122312400 | *CERKL* | 4.2737E-04 | 2.0993E-04 | 6.4921e-6 | 0.4446 | 0.0977 |
| AX-147194100 | GL018965$_{MC1R}$ | AX-147108549 | 7 | 122326907 | *CERKL* | 4.2737E-04 | 2.0993E-04 | 6.4921e-6 | 0.4446 | 0.0977 |
| AX-147194100 | GL018965$_{MC1R}$ | AX-147130487 | 7 | 122789755 | *CERKL* | 2.5568E-04 | 1.2608E-04 | 3.2743e-6 | 0.4690 | 0.0998 |
| AX-147194100 | GL018965$_{MC1R}$ | AX-147042416 | 7 | 123769296 | | 5.8267E-04 | 2.7234E-04 | 6.4890e-6 | 0.4923 | 0.1082 |
| AX-147194100 | GL018965$_{MC1R}$ | AX-147055205 | 7 | 123947622 | | 7.2842E-04 | 3.5218E-04 | 1.1992e-5 | 0.4334 | 0.0982 |
| AX-147194100 | GL018965$_{MC1R}$ | AX-147012957 | 7 | 124038246 | | 6.8745E-04 | 3.2938E-04 | 1.0378e-5 | 0.4478 | 0.1007 |
| AX-147194100 | GL018965$_{MC1R}$ | AX-147093243 | 7 | 124276367 | | 7.0550E-04 | 3.3797E-04 | 1.0748e-5 | 0.4464 | 0.1005 |
| AX-147194100 | GL018965$_{MC1R}$ | AX-147016943 | 7 | 124450559 | | 8.3756E-04 | 4.0364E-04 | 1.4186e-5 | 0.4292 | 0.0980 |
| AX-147194100 | GL018965$_{MC1R}$ | AX-147091088 | 7 | 124468761 | | 9.7912E-04 | 4.6971E-04 | 1.7019e-5 | 0.4254 | 0.0981 |
| AX-147194100 | GL018965$_{MC1R}$ | AX-147047527 | 7 | 124484859 | | 8.1063E-04 | 3.9103E-04 | 1.3655e-5 | 0.4300 | 0.0980 |
| AX-147194100 | GL018965$_{MC1R}$ | AX-146994454 | 7 | 124724425 | | 4.1204E-04 | 1.9998E-04 | 5.5979e-6 | 0.4620 | 0.1008 |
| AX-147194100 | GL018965$_{MC1R}$ | AX-147037439 | 7 | 124835395 | | 9.4555E-04 | 4.4775E-04 | 1.4866e-5 | 0.4428 | 0.1014 |
| AX-147194100 | GL018965$_{MC1R}$ | AX-147111975 | 7 | 124841635 | | 4.9853E-04 | 2.3629E-04 | 5.9160e-6 | -0.4826 | 0.1056 |
| AX-147194100 | GL018965$_{MC1R}$ | AX-147088907 | 7 | 125339331 | | 3.1752E-04 | 1.5488E-04 | 4.0336e-6 | 0.4709 | 0.1012 |
| AX-147194100 | GL018965$_{MC1R}$ | AX-147124126 | 7 | 125675080 | | 6.3684E-04 | 3.0163E-04 | 8.4546e-6 | 0.4675 | 0.1040 |
| AX-147194100 | GL018965$_{MC1R}$ | AX-147101138 | 7 | 126032136 | | 1.9367E-04 | 9.3955E-05 | 1.8120e-6 | 0.5101 | 0.1058 |
| AX-147194100 | GL018965$_{MC1R}$ | AX-147000520 | 10 | 27978262 | | 5.0375E-04 | 2.2771E-04 | 3.2897e-6 | -0.5599 | 0.1192 |
| AX-147194100 | GL018965$_{MC1R}$ | AX-147080769 | 10 | 35705012 | | 2.8536E-04 | 1.3103E-04 | 1.5338e-6 | -0.5787 | 0.1192 |
| AX-147194100 | GL018965$_{MC1R}$ | AX-147021837 | 10 | 35710262 | | 2.8536E-04 | 1.3103E-04 | 1.5338e-6 | -0.5787 | 0.1192 |
| AX-147194100 | GL018965$_{MC1R}$ | AX-147085326 | 10 | 35780647 | | 2.3564E-04 | 1.0871E-04 | 1.1764e-6 | -0.5857 | 0.1193 |
| AX-147194100 | GL018965$_{MC1R}$ | AX-147138602 | 10 | 35787696 | | 2.8536E-04 | 1.3103E-04 | 1.5338e-6 | -0.5787 | 0.1192 |
| AX-147194100 | GL018965$_{MC1R}$ | AX-147066293 | 10 | 35798248 | | 2.8536E-04 | 1.3103E-04 | 1.5338e-6 | -0.5787 | 0.1192 |
| AX-147194100 | GL018965$_{MC1R}$ | AX-147156763 | 10 | 35873975 | | 2.8536E-04 | 1.3103E-04 | 1.5338e-6 | -0.5787 | 0.1192 |
| AX-147194100 | GL018965$_{MC1R}$ | AX-147076047 | 10 | 35945454 | | 2.8536E-04 | 1.3103E-04 | 1.5338e-6 | -0.5787 | 0.1192 |
| AX-147194100 | GL018965$_{MC1R}$ | AX-147001940 | 10 | 35960043 | | 3.2393E-04 | 1.4823E-04 | 1.8176e-6 | -0.5746 | 0.1192 |
| AX-147194100 | GL018965$_{MC1R}$ | AX-147061119 | 10 | 35971599 | | 2.8536E-04 | 1.3103E-04 | 1.5338e-6 | -0.5787 | 0.1192 |
| AX-147194100 | GL018965$_{MC1R}$ | AX-147085327 | 10 | 36294963 | | 3.9848E-04 | 1.8123E-04 | 2.3857e-6 | -0.5685 | 0.1193 |
| AX-147194100 | GL018965$_{MC1R}$ | AX-147109239 | 10 | 36322653 | | 4.1819E-04 | 1.8994E-04 | 2.5459e-6 | -0.5669 | 0.1193 |
| AX-147194100 | GL018965$_{MC1R}$ | AX-147091912 | 10 | 36396275 | | 6.4688E-04 | 2.8874E-04 | 4.2354e-6 | -0.5616 | 0.1210 |
| AX-147194100 | GL018965$_{MC1R}$ | AX-147071251 | 10 | 36420720 | | 6.7909E-04 | 3.0265E-04 | 4.5153e-6 | -0.5601 | 0.1210 |
| AX-147194100 | GL018965$_{MC1R}$ | AX-147026289 | 10 | 36432792 | | 4.4452E-04 | 2.0145E-04 | 2.7390e-6 | -0.5659 | 0.1195 |
| AX-147194100 | GL018965$_{MC1R}$ | AX-147013722 | 11 | 84453876 | | 6.6195E-04 | 3.1167E-04 | 8.4778e-6 | 0.4730 | 0.1053 |
| AX-147194100 | GL018965$_{MC1R}$ | AX-147022140 | 13 | 121883005 | | 5.0260E-04 | 2.6794E-04 | 1.1688e-5 | -0.3533 | 0.0799 |
| AX-147194100 | GL018965$_{MC1R}$ | AX-147172253 | 13 | 125018521 | | 9.1399E-04 | 4.4266E-04 | 1.6434e-5 | -0.4187 | 0.0964 |
| AX-147194100 | GL018965$_{MC1R}$ | AX-147135656 | 13 | 125045081 | | 4.5613E-04 | 2.2561E-04 | 7.4444e-6 | -0.4327 | 0.0957 |
| AX-147194100 | GL018965$_{MC1R}$ | AX-147143979 | 13 | 125054916 | | 4.8816E-04 | 2.3979E-04 | 7.7769e-6 | -0.4372 | 0.0969 |
| AX-147194100 | GL018965$_{MC1R}$ | AX-147157331 | 13 | 125135433 | | 6.4401E-04 | 2.8982E-04 | 4.7497e-6 | -0.5468 | 0.1184 |
| AX-147194100 | GL018965$_{MC1R}$ | AX-147162840 | 13 | 125186251 | | 4.6997E-04 | 2.3228E-04 | 7.7077e-6 | -0.4320 | 0.0957 |
| AX-147194100 | GL018965$_{MC1R}$ | AX-147169535 | 13 | 125759786 | | 9.5806E-04 | 4.5178E-04 | 1.4682e-5 | 0.4472 | 0.1023 |
| AX-147194100 | GL018965$_{MC1R}$ | AX-146990796 | 13 | 130087549 | | 2.9424E-04 | 1.4473E-04 | 3.9074e-6 | -0.4641 | 0.0996 |
| AX-147194100 | GL018965$_{MC1R}$ | AX-147083441 | 13 | 130093238 | | 2.0452E-04 | 1.0147E-04 | 2.5246e-6 | -0.4735 | 0.0996 |
| AX-147194100 | GL018965$_{MC1R}$ | AX-147168584 | 13 | 130399123 | | 7.2975E-04 | 3.4819E-04 | 1.0931e-5 | -0.4493 | 0.1013 |
| AX-147194100 | GL018965$_{MC1R}$ | AX-147176146 | 13 | 131452813 | | 7.4449E-04 | 3.6512E-04 | 1.3482e-5 | -0.4161 | 0.0948 |
| AX-147194100 | GL018965$_{MC1R}$ | AX-147163795 | 13 | 131458395 | | 9.3323E-04 | 4.5484E-04 | 1.7496e-5 | -0.4107 | 0.0948 |
| AX-147194100 | GL018965$_{MC1R}$ | AX-147119230 | 14 | 50353708 | | 4.6767E-04 | 2.3289E-04 | 8.0370e-6 | -0.4236 | 0.0940 |
| AX-147194100 | GL018965$_{MC1R}$ | AX-147104090 | 14 | 50362060 | | 4.4131E-04 | 2.2010E-04 | 7.5220e-6 | -0.4248 | 0.0940 |
| AX-147194100 | GL018965$_{MC1R}$ | AX-147158354 | 14 | 50368832 | | 4.2296E-04 | 2.1117E-04 | 7.1617e-6 | -0.4259 | 0.0940 |
| AX-147194100 | GL018965$_{MC1R}$ | AX-147048868 | 14 | 50374787 | | 4.2296E-04 | 2.1117E-04 | 7.1617e-6 | -0.4259 | 0.0940 |
| AX-147194100 | GL018965$_{MC1R}$ | AX-147092266 | 14 | 50381230 | | 4.7946E-04 | 2.3862E-04 | 8.2730e-6 | -0.4230 | 0.0940 |
| AX-147194100 | GL018965$_{MC1R}$ | AX-147169093 | 14 | 50413907 | | 4.2296E-04 | 2.1117E-04 | 7.1617e-6 | -0.4259 | 0.0940 |
| AX-147194100 | GL018965$_{MC1R}$ | AX-147109548 | 14 | 50425012 | | 4.7946E-04 | 2.3862E-04 | 8.2730e-6 | -0.4230 | 0.0940 |

| | | | | | | | | | | |
|---|---|---|---|---|---|---|---|---|---|---|
| AX-147194100 | GL018965$_{MC1R}$ | AX-147085696 | 14 | 50431143 | | 4.7946E-04 | 2.3862E-04 | 8.2730e-6 | -0.4230 | 0.0940 |
| AX-147194100 | GL018965$_{MC1R}$ | AX-147051432 | 14 | 50448244 | | 4.2404E-04 | 2.1165E-04 | 7.1723e-6 | -0.4261 | 0.0941 |
| AX-147194100 | GL018965$_{MC1R}$ | AX-147120724 | 14 | 50458962 | | 4.4830E-04 | 2.2348E-04 | 7.6537e-6 | -0.4246 | 0.0940 |
| AX-147194100 | GL018965$_{MC1R}$ | AX-147054073 | 14 | 50467207 | | 4.7946E-04 | 2.3862E-04 | 8.2730e-6 | -0.4230 | 0.0940 |
| AX-147194100 | GL018965$_{MC1R}$ | AX-147158872 | 14 | 50473592 | | 4.7946E-04 | 2.3862E-04 | 8.2730e-6 | -0.4230 | 0.0940 |
| AX-147194100 | GL018965$_{MC1R}$ | AX-147096381 | 14 | 50489652 | | 4.7946E-04 | 2.3862E-04 | 8.2730e-6 | -0.4230 | 0.0940 |
| AX-147194100 | GL018965$_{MC1R}$ | AX-147143173 | 14 | 50502520 | | 4.7946E-04 | 2.3862E-04 | 8.2730e-6 | -0.4230 | 0.0940 |
| AX-147194100 | GL018965$_{MC1R}$ | AX-147122126 | 14 | 50515001 | | 4.7946E-04 | 2.3862E-04 | 8.2730e-6 | -0.4230 | 0.0940 |
| AX-147194100 | GL018965$_{MC1R}$ | AX-147102230 | 14 | 50769091 | | 7.9217E-04 | 3.9030E-04 | 1.4984e-5 | -0.4078 | 0.0934 |
| AX-147194100 | GL018965$_{MC1R}$ | AX-147100330 | 14 | 50803809 | | 6.8500E-04 | 3.3876E-04 | 1.2674e-5 | -0.4114 | 0.0934 |
| AX-147194100 | GL018965$_{MC1R}$ | AX-147132328 | 14 | 50816927 | | 9.2989E-04 | 4.5730E-04 | 1.8232e-5 | -0.4014 | 0.0929 |
| AX-147194100 | GL018965$_{MC1R}$ | AX-147168113 | 14 | 51531949 | | 8.6289E-04 | 4.1206E-04 | 1.3897e-5 | -0.4382 | 0.1000 |
| AX-147194100 | GL018965$_{MC1R}$ | AX-147071662 | 14 | 51543974 | | 8.5342E-04 | 4.0767E-04 | 1.3718e-5 | -0.4385 | 0.1000 |
| AX-147194100 | GL018965$_{MC1R}$ | AX-147157342 | 14 | 51636241 | | 6.7408E-04 | 3.3584E-04 | 1.2906e-5 | -0.4044 | 0.0919 |
| AX-147194100 | GL018965$_{MC1R}$ | AX-147003701 | 14 | 51666539 | | 6.7408E-04 | 3.3584E-04 | 1.2906e-5 | -0.4044 | 0.0919 |
| AX-147194100 | GL018965$_{MC1R}$ | AX-147172267 | 14 | 51726019 | | 9.5801E-04 | 4.7272E-04 | 1.9242e-5 | -0.3964 | 0.0920 |
| AX-147194100 | GL018965$_{MC1R}$ | AX-147088193 | 17 | 36238191 | | 9.4876E-04 | 3.7887E-04 | 1.2357e-6 | 0.7798 | 0.1591 |
| AX-147194100 | GL018965$_{MC1R}$ | AX-147005577 | 19 | 14344377 | | 6.9107E-04 | 3.1367E-04 | 6.0332e-6 | -0.5259 | 0.1151 |
| AX-147194100 | GL018965$_{MC1R}$ | AX-147126472 | 19 | 14359988 | | 6.9107E-04 | 3.1367E-04 | 6.0332e-6 | -0.5259 | 0.1151 |
| AX-147194100 | GL018965$_{MC1R}$ | AX-147175773 | 19 | 14387963 | | 6.9107E-04 | 3.1367E-04 | 6.0332e-6 | -0.5259 | 0.1151 |
| AX-147194100 | GL018965$_{MC1R}$ | AX-147116584 | 20 | 13978882 | *SLC24A4* | 5.2252E-04 | 2.1727E-04 | 7.9062e-7 | -0.7450 | 0.1493 |
| AX-147194100 | GL018965$_{MC1R}$ | AX-147145778 | 20 | 14039605 | | 5.2252E-04 | 2.1727E-04 | 7.9062e-7 | -0.7450 | 0.1493 |
| AX-147194100 | GL018965$_{MC1R}$ | AX-147178398 | 20 | 14475465 | | 6.4882E-04 | 2.7402E-04 | 1.5920e-6 | -0.6825 | 0.1408 |
| AX-147194100 | GL018965$_{MC1R}$ | AX-147173265 | 20 | 16497558 | | 2.2403E-04 | 1.0833E-04 | 2.1809e-6 | 0.5057 | 0.1057 |
| AX-147194100 | GL018965$_{MC1R}$ | AX-147135945 | 20 | 16510520 | *TTC8* | 2.2403E-04 | 1.0833E-04 | 2.1809e-6 | 0.5057 | 0.1057 |
| AX-147194100 | GL018965$_{MC1R}$ | AX-147148645 | 20 | 16519051 | *TTC8* | 2.2403E-04 | 1.0833E-04 | 2.1809e-6 | 0.5057 | 0.1057 |
| AX-147194100 | GL018965$_{MC1R}$ | AX-147036995 | 20 | 16659910 | *TTC8* | 3.6511E-04 | 1.7116E-04 | 3.2020e-6 | -0.5213 | 0.1108 |
| AX-147194100 | GL018965$_{MC1R}$ | AX-147147930 | 20 | 16853972 | | 3.2105E-04 | 1.5104E-04 | 2.7190e-6 | -0.5250 | 0.1108 |
| AX-147194100 | GL018965$_{MC1R}$ | AX-147135946 | 20 | 16860941 | | 5.1405E-04 | 2.2424E-04 | 1.8479e-6 | -0.6317 | 0.1311 |
| AX-147194100 | GL018965$_{MC1R}$ | AX-147131466 | 20 | 16911333 | | 5.1405E-04 | 2.2424E-04 | 1.8479e-6 | -0.6317 | 0.1311 |
| AX-147194100 | GL018965$_{MC1R}$ | AX-147178791 | 20 | 16920075 | | 5.1405E-04 | 2.2424E-04 | 1.8479e-6 | -0.6317 | 0.1311 |
| AX-147194100 | GL018965$_{MC1R}$ | AX-147108233 | 20 | 16926472 | | 5.1405E-04 | 2.2424E-04 | 1.8479e-6 | -0.6317 | 0.1311 |
| AX-147194100 | GL018965$_{MC1R}$ | AX-147100829 | 20 | 16938325 | | 5.1405E-04 | 2.2424E-04 | 1.8479e-6 | -0.6317 | 0.1311 |
| AX-147194100 | GL018965$_{MC1R}$ | AX-147049600 | 20 | 16953872 | *SPATA7* | 5.5919E-04 | 2.4333E-04 | 2.0795e-6 | -0.6286 | 0.1312 |
| AX-147194100 | GL018965$_{MC1R}$ | AX-147074687 | 20 | 16988376 | *SPATA7* | 1.3397E-04 | 6.5432E-05 | 1.1097e-6 | 0.5236 | 0.1064 |
| AX-147194100 | GL018965$_{MC1R}$ | AX-147057297 | 20 | 16998158 | *SPATA7* | 1.3397E-04 | 6.5432E-05 | 1.1097e-6 | -0.5236 | 0.1064 |
| AX-147194100 | GL018965$_{MC1R}$ | AX-147052184 | 20 | 17008788 | | 5.1405E-04 | 2.2424E-04 | 1.8479e-6 | -0.6317 | 0.1311 |
| AX-147194100 | GL018965$_{MC1R}$ | AX-147032232 | 20 | 17032215 | | 9.4242E-04 | 4.4478E-04 | 1.4436e-5 | -0.4471 | 0.1022 |
| AX-147194100 | GL018965$_{MC1R}$ | AX-147100830 | 20 | 18491550 | | 8.4044E-04 | 3.9069E-04 | 1.0751e-5 | -0.4746 | 0.1069 |
| AX-147194100 | GL018965$_{MC1R}$ | AX-147047073 | 20 | 18744378 | | 3.6268E-04 | 1.7440E-04 | 4.2578e-6 | -0.4827 | 0.1040 |
| AX-147194100 | GL018965$_{MC1R}$ | AX-147100831 | 20 | 19305852 | | 1.0146E-04 | 5.0993E-05 | 1.0467e-6 | -0.4956 | 0.1004 |
| AX-147194100 | GL018965$_{MC1R}$ | AX-147065735 | GL018710 | 7572 | | 9.9036E-04 | 4.6419E-04 | 1.4698e-5 | 0.4527 | 0.1036 |
| AX-147194100 | GL018965$_{MC1R}$ | AX-147009578 | GL018710 | 7578 | | 7.4775E-04 | 3.5603E-04 | 1.1142e-5 | 0.4505 | 0.1016 |
| AX-147194100 | GL018965$_{MC1R}$ | AX-147128287 | GL018710 | 7579 | | 8.4182E-04 | 3.9605E-04 | 1.2029e-5 | 0.4583 | 0.1038 |

**Table S10: Primers and PCR conditions for additional manual genotyping**

| Gene | Sequence | Size of PCR product | PCR conditions | Digestion Conditions | Genotyping method |
|---|---|---|---|---|---|
| ASIP a | CAGGAAGGCACATCCTCTTT | 418 | 56°C 42 cycles | | capillary electrophoresis |
| ASIP a | TTCCCAAACCAAAGAAGTCAA | 418 | 56°C 42 cycles | | capillary electrophoresis |
| ASIP | GCTTCGAAGAAGAAGGTGGCG | 193 | 57°C 42 cycles | | allele specific PCR |
| ASIP | ACTTCGAAGAAGAAGGTGGCG | 193 | 57°C 42 cycles | | allele specific PCR |
| ASIP | CTCAGCAGTTGGGGTTGAG | 193 | 57°C 42 cycles | | allele specific PCR |
| MC1R Ed e | CACCAGCCCCTTCCTGAT | 473-479-503-509 | 60°C 42 cycles | | capillary electrophoresis |
| MC1R Ed e | TCTTCTACGCACTGCGCTAC | 473-479-503-510 | 60°C 42 cycles | | capillary electrophoresis |
| TYR c | GATCCCTGTACCTGGGACATC | 225 | 55°C 42 cycles | Nci 37°C 15 min | RFLP PCR |
| TYR c | ACTACAATTGAAAGCCCGTTTCA | 225 | 55°C 42 cycles | Nci 37°C 15 min | RFLP PCR |
| TYR $C^h$ | TGAAATTGGCAGCTTTGTCCA | 195 | 55°C 42 cycles | BsaXI 37°C 15 min | RFLP PCR |
| TYR $C^h$ | AGCTCTGTCGGCTATTGTACT | 195 | 55°C 42 cycles | BsaXI 37°C 15 min | RFLP PCR |
