## Additional Figures for "A genome-wide epistatic network underlies the molecular architecture of continuous color variation of body extremities: a rabbit model"

**Figure S1: Coat coloration genetic determinism for white rabbits** **2**
**Figure S2: Coat coloration genetic determinism for spotted rabbits** **3**
**Figure S3: Linkage disequilibrium heatmap between both Ocu15$_{KIT}$ and GL018754 regions** **4**
**Figure S4: Coat coloration genetic determinism for remaining colored rabbits (P3 to P6 groups)** **5**
**Figure S5: Genetic components that contribute to the extremities coat color determinism** **6**
**Figure S6: Coat coloration phenotypic distribution within the whole experimental design** **7**

**<u>Figure S1</u>: Coat coloration genetic determinism for white rabbits**

**a.** Genotypic distribution of the best SNP marker associated located within the Ocu1$_{TYR}$ interval. **b.** Manhattan plot. GWAS were performed comparing white rabbits (phenotype 1 (P1)) versus the 5 remaining ones under different genetic models.

**a**

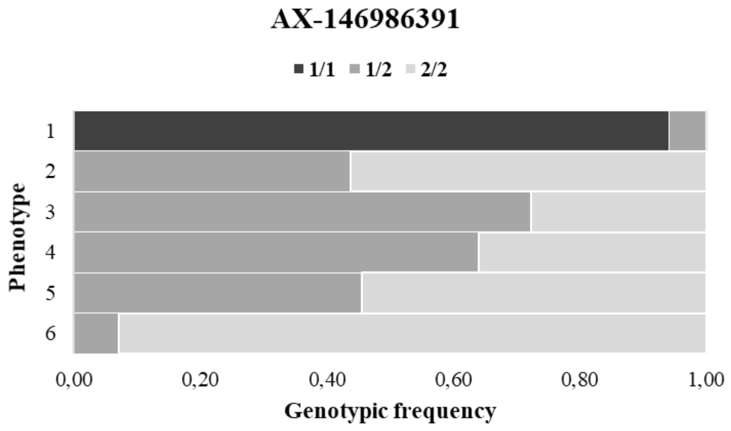


**b**

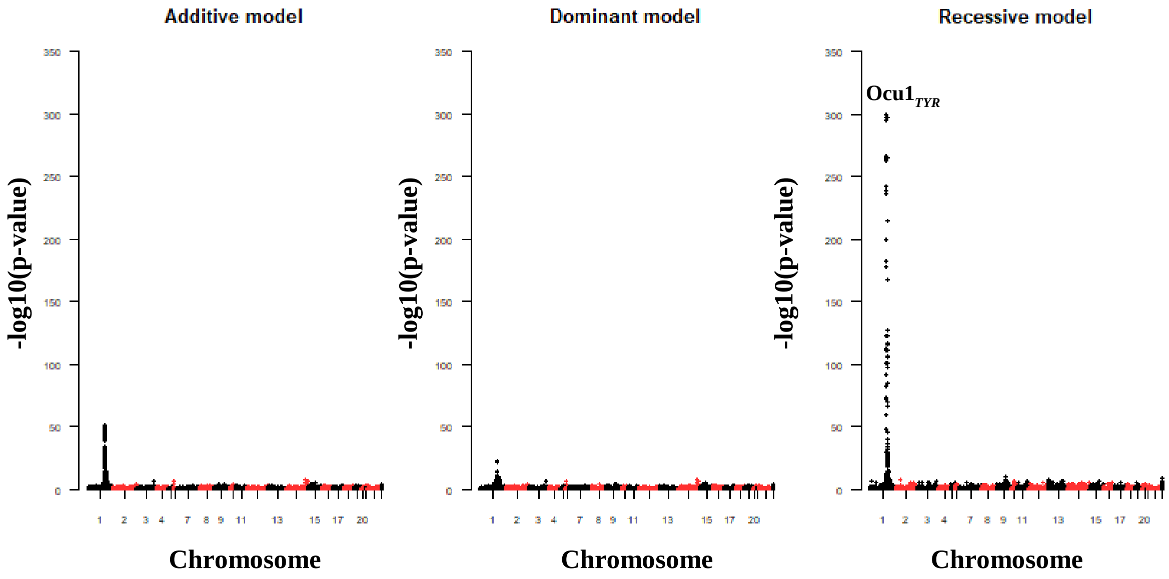

**Figure S2: Coat coloration genetic determinism for spotted rabbits**

**a.** Genotypic distribution of the best SNP marker associated located within the Ocu15$_{KIT}$ interval. **d.** Manhattan plot. GWAS were performed comparing spotted rabbits (phenotype 2 (P2)) to combined light to dark brown coat-colored rabbits (P3 to P6), but excluding white rabbits (phenotype 1 (P1))

**a**

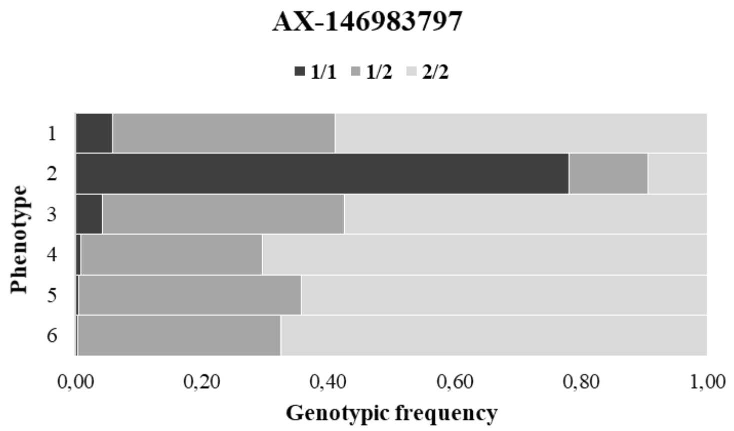


**b**

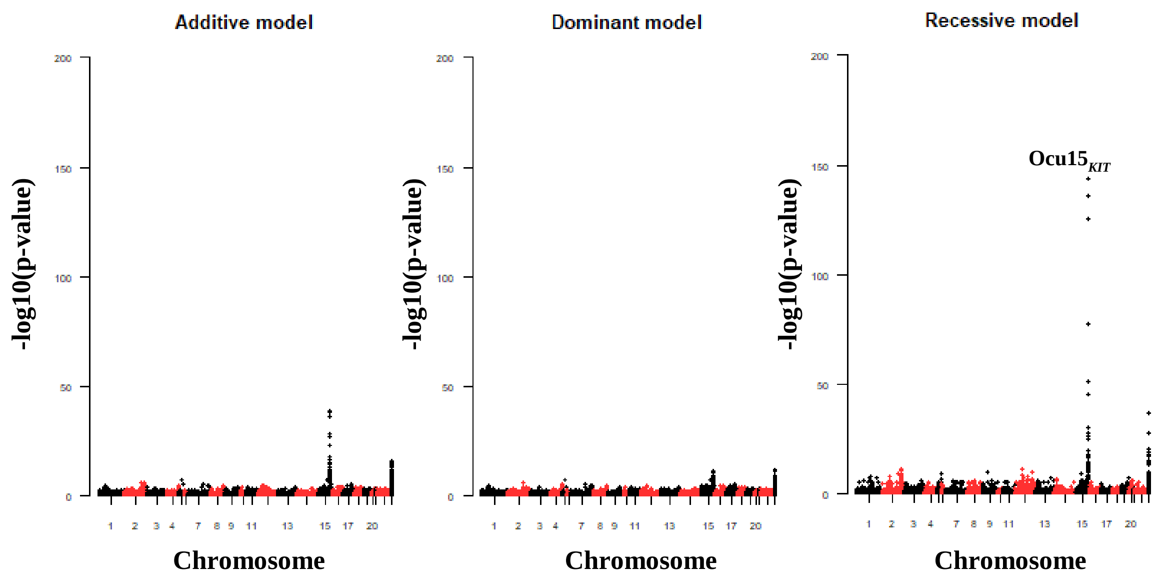

**<u>Figure S3</u>: Linkage disequilibrium heatmap between both Ocu15$_{KIT}$ and GL018754 regions**
The red box showed a strong LD between a block of SNP markers from the Ocu15$_{KIT}$ locus and the scaffold GL018754 strongly suggested that they are linked.

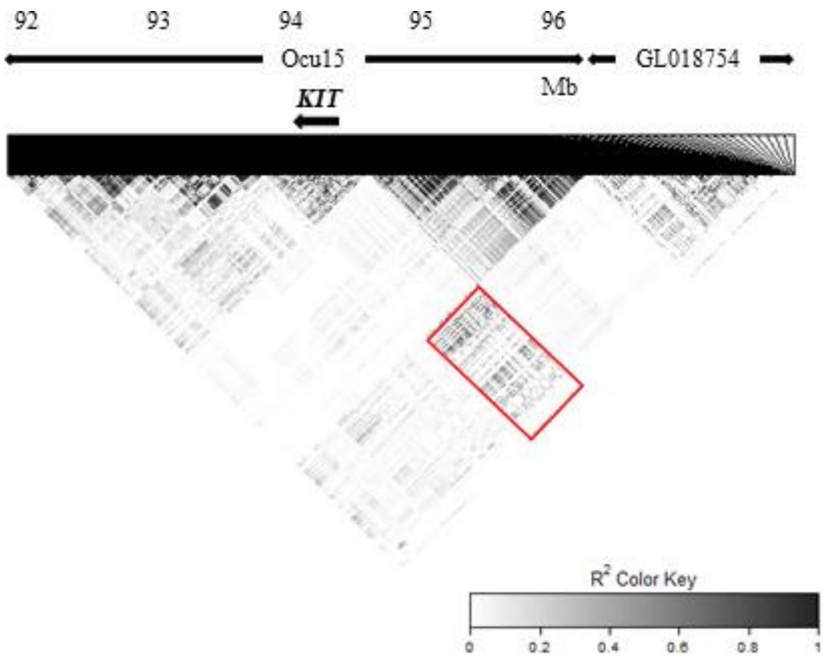

**Figure S4: Coat coloration genetic determinism for remaining colored rabbits (P3 to P6 groups)**

**a.** Estimation of number of QTL involved. **b.** Proportion of the phenotypic variance explained by QTL using the additive linear mixed model. **c.** Proportion of the genetic variance explained by QTL using the additive linear mixed model. **d**. The SNP effects were tested with a Bayesian sparse animal mixed model and the sparse probabilities evaluated from the total effect size for a given SNP on sliding windows containing 20 SNPs were plotted.

QTL: quantitative trait locus. Stars correspond to loci showing a probability above 0.15 for being a QTL.

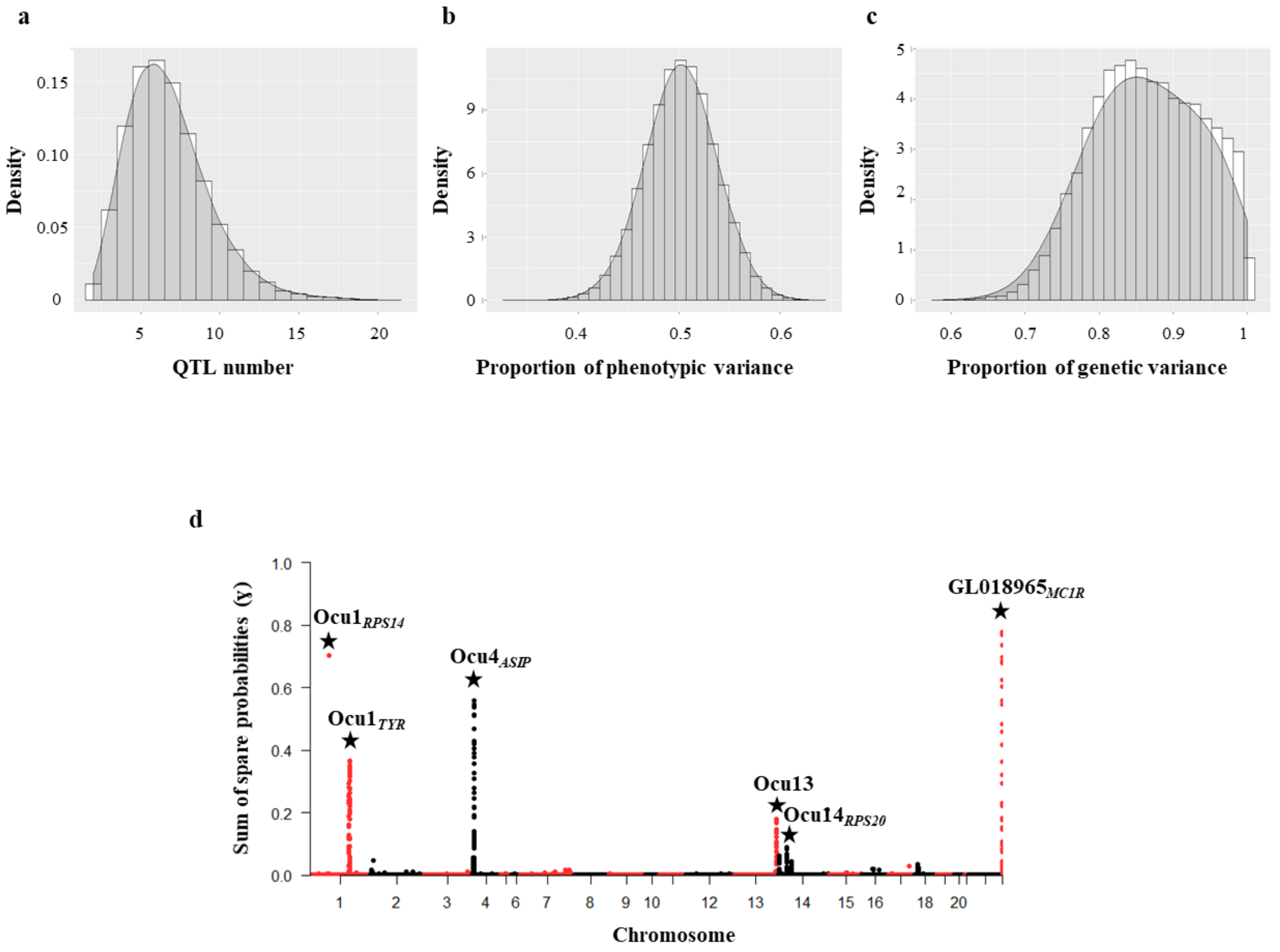

**<u>Figure S5</u>: Genetic components that contribute to the extremities coat color determinism**

**a**. Phenotypic prediction. The boxplot represents the predicted phenotype with the underlined fitted best model for individuals colored from light to dark (P3 to P6) since P1 and P2 are major genes. **b.** Genetic effects. Both individual and interaction effects on the genetic determinism of coat coloration for phenotypes P3 to P6 were evaluated (*, ** and *** for p-values < 0.1, 0.01 and 0.001, respectively).

**a**

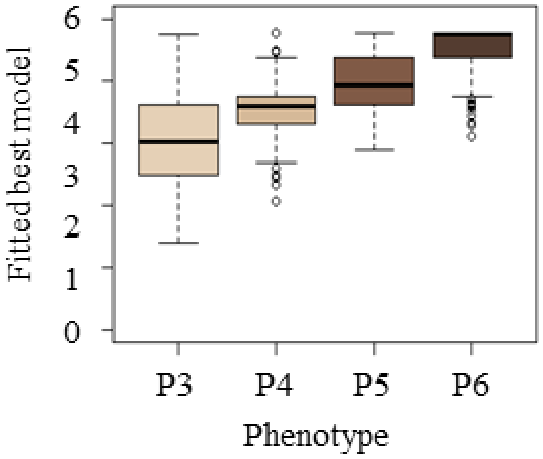


**b**

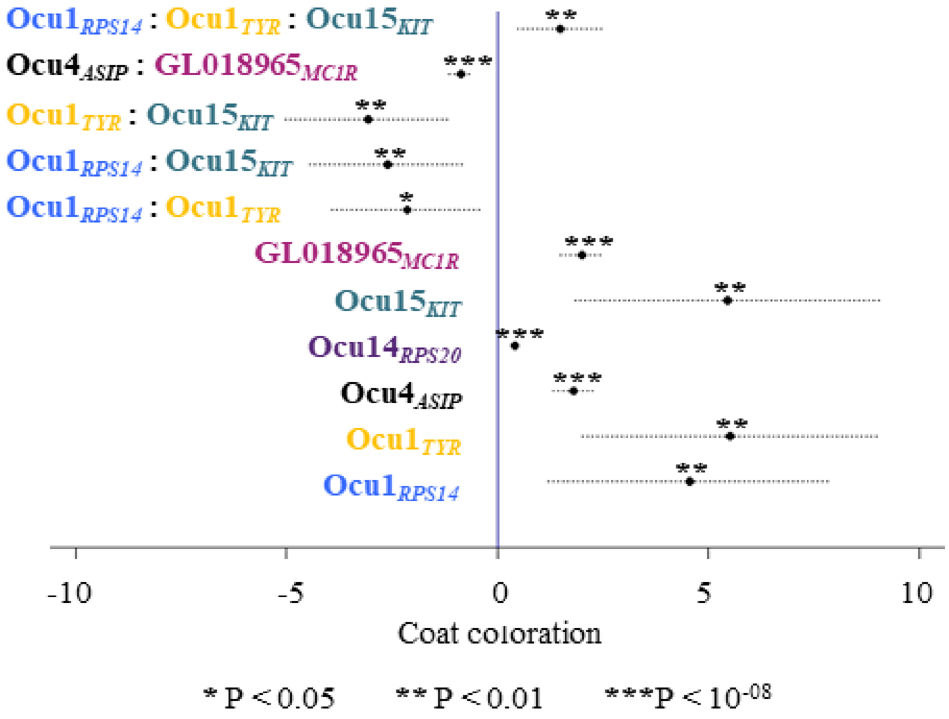


b

**<u>Figure S6</u>: Extremities coat coloration phenotypic classification within the whole experimental design**
**a**. Correlation between visual inspection of phenotypic coat coloration (classified in 5 groups from A to E) and luminescence measurement for the 5 groups. **b.** Distribution of coloration within the 20 sires' families. **c.** Distribution of parents' coloration within the 6 groups of color of body extremities. Filled colors are paternal colors and hashed colors are maternal colors.

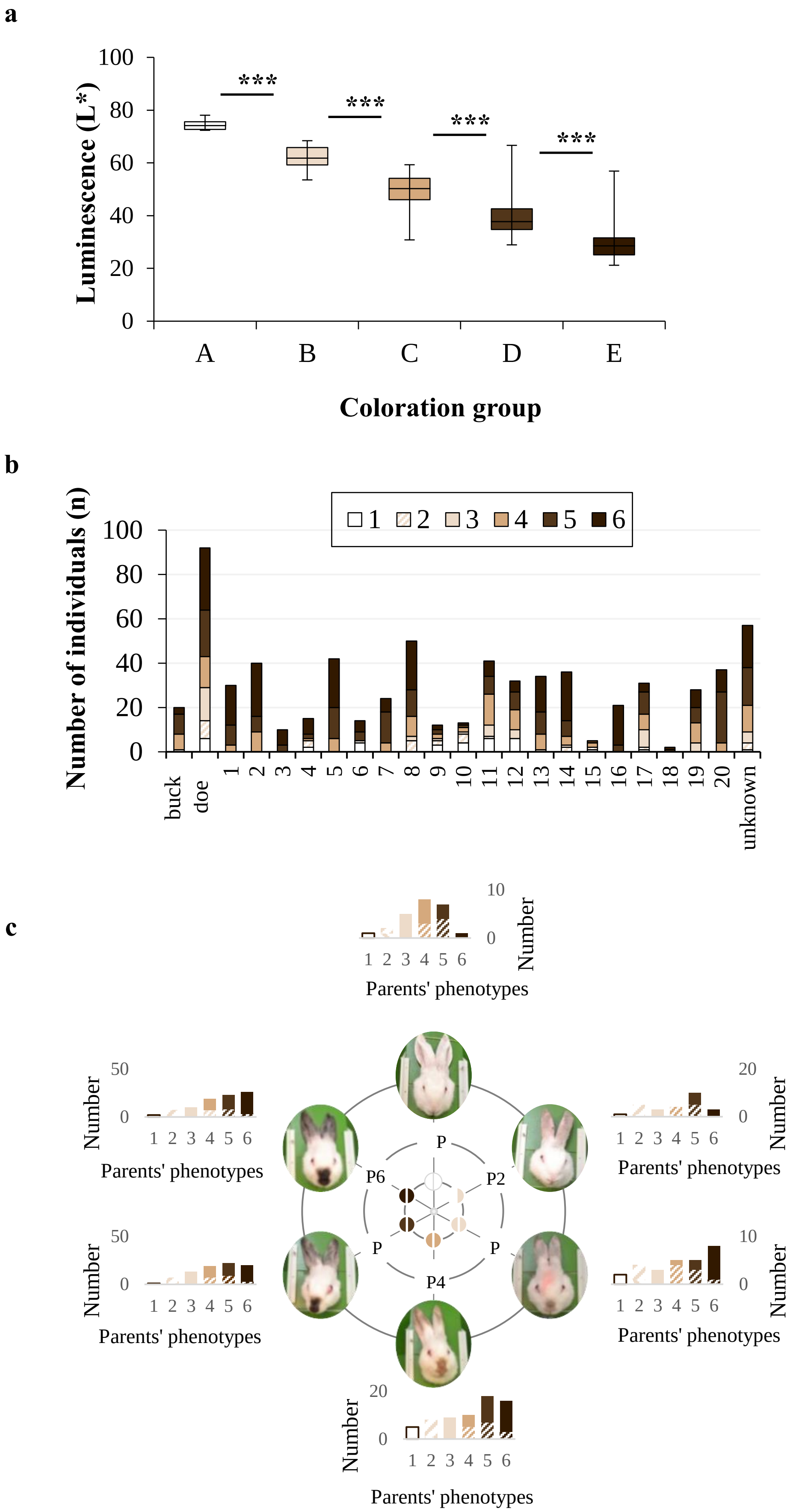
